# Complete chloroplast genome of African Baobab (*Adansonia digitata* L.): structural characterization, comparative genomics, and phylogenetic placement within Malvaceae

**DOI:** 10.64898/2026.02.26.708206

**Authors:** Oketch Fredrick Onyango, Zipporah Muchiri, Everlyne Osir Owiro, Mathews Wafula, Oscar Mwaura, Rose Kigathi

## Abstract

Chloroplast genomes are invaluable resources for plant genomic research, providing insights into genome evolution and molecular adaptation. With the growing scientific and economic interest in *Adansonia digitata*, a comprehensive characterization of its chloroplast is timely and necessary. A complete chloroplast genome of *A. digitata* was assembled, annotated, and characterized. Comparative structural analysis was conducted against other *Adansonia* species, and the assembly was validated through phylogenetic placement within Malvaceae. The assembled genome exhibits the canonical quadripartite organization, spanning 160,061 bp with a GC content of 36.88%, 79 protein-coding genes, 32 tRNAs, and 4 rRNAs. Repeat analysis identified 100 simple sequence repeat motifs, predominantly A/T-rich mononucleotide types (76%), alongside 50 long sequence repeats dominated by forward (26) and palindromic (17) repeats. Comparative analysis with other *Adansonia* species revealed conserved genome structure, with minor IR boundary shifts involving the ndhF gene, and ycf1 duplication in *A. gregorii* and *A. grandidieri.* Average nucleotide identity exceeded 99% across all *Adansonia* species, with near-complete similarity (ANI ≈ 99.96%) observed with the putative *A. kilima.* All predicted RNA editing events were nonsynonymous, dominated by C→U conversions (55.02%). Codon usage showed non-random synonymous preferences biased toward A/U-ending codons, driven primarily by mutational pressure with detectable gene-specific translational selection. Nucleotide diversity (π) was higher in intergenic spacers (0.00490 ± 0.00574) than in coding regions (0.00167 ± 0.00199), with the majority of genomic regions showing no sequence variation (π = 0). Substitution patterns indicated pervasive purifying selection, with relatively high but insignificant signals in *matK, ycf1, accD*, and *rpoB*. Phylogenomic analyses placed the assembled *A. digitata* chloroplast genome within the *Adansonia* lineage, consistent with its established systematic position. This study provides detailed insight into the chloroplast genome of *A. digitata*, and the findings will contribute towards advancing its genomic research.

## Introduction

The genus *Adansonia* comprises wild fruit tree species native to Madagascar, mainland Africa, and Australia^1^. The trees are believed to be of ancient lineages, with archaeological and paleobotanical evidence suggesting their origin and subsequent dispersal to the breakup of the supercontinent Gondwana ^1^. Recent evidence suggests that Madagascar is the center of baobab speciation, with *Adansonia digitata* in mainland Africa and *Adansonia gregorii* in Australia diverging from a common ancestral lineage in Madagascar ^1,2^. Globally, *A. digitata* is the most widespread species, while six others, *Adansonia grandidieri, A. rubrostipa, A. za, A. madagascariensis, A. perrieri,* and *A. suarezensis*, are endemic to Madagascar ^1,3^, whereas *A. gregorii* is native to Australia^1^. Although a ninth putative species, *A. kilima*, has been proposed to occur in sympatry with *A. digitata* in the highlands of mainland Africa, the evidence remains inconclusive, and the species has not been formally recognized ^4–6^. Despite its iconic status and wide-ranging ecological and socio-economic benefits^7–12^, the chloroplast genome of *A. digitata* remains incompletely characterized, with gaps in detailed structural organization, sequence diversity patterns, and evolutionary and functional interpretation. These gaps are likely to constrain efforts to explore its chloroplast genome for biotechnological applications and applied genomic studies^13,14^. As a long-lived, large-flowering tree with increasing demand for its nutrient-rich products ^11,15^, *A. digitata* represents a good model species for exploring how chloroplast genome structure and function relate to longevity, stress tolerance, and environmental resilience in woody angiosperms.

Understanding the significance of these gaps requires anchoring them within the broader context of chloroplast genome biology. The chloroplast genome is a key component of plant cells and has co-evolved with plants for millions of years following the original endosymbiotic event^16,17^. In most angiosperms, chloroplast genomes exhibit a conserved quadripartite structure consisting of large and small single-copy regions separated by a pair of inverted repeats^18^. The plastome is typically maternally inherited and evolves at a moderate rate, making it a reliable marker for phylogenetic reconstruction and comparative genomic studies ^19,20^. Although the overall structure of most chloroplast genomes is conserved, lineage-specific variation does occur, with certain genes being lost, pseudogenized, duplicated, inverted, or shifted within inverted repeat boundaries ^21^ - evolutionary signatures that are commonly used to reconstruct evolutionary relationships and infer divergence patterns among plant lineages ^22–24^.

Genes within the chloroplast genome, particularly those involved in photosynthesis, photoprotection, and stress response, also exhibit variation linked to environmental adaptation, providing insights into how species respond to diverse ecological conditions ^25,26^. Beyond evolutionary and adaptive studies, chloroplast genome sequences are valuable for species validation, population-level research, and conservation planning ^27–29^, and in applied biotechnology research, particularly in plastid-based genetic engineering, where chloroplast transformation enables targeted expression and containment of transgenes^19,30,31^. These features make chloroplast genome studies a suitable tool for gaining insight into the genetic basis of *A. digitata* ecological and functional diversity.

Although chloroplast genome sequences of *A. digitata* are available in public repositories such as the National Center for Biotechnology Information (NCBI), they have not been adequately examined through detailed comparative analyses. This study, therefore, sought to generate and comprehensively characterise the chloroplast genome of *A. digitata* through complete sequencing, assembly, and annotation. Specifically, the study aimed to describe its genome structure, gene content, and major genomic features, compare its chloroplast genome structure with those of other *Adansonia* species, and infer phylogenomic relationships with the selected chloroplast genomes of related members of the family Malvaceae. The goal was to produce a complete, quality chloroplast genome that strengthens existing genomic resources for *A. digitata* and supports continued applied genomic research.

## Material and Methods

### Plant material and whole genome sequencing

A mature fruit pod was collected from a naturally growing A. digitata tree in Gazi Bay, Kwale County, Kenya (−4.422680, 39.507275). The tree was taxonomically identified as A. digitata based on morphological features as described in ^32^. The sampled specimen was labeled and transported to the Pwani University Bioscience Research Centre (PUBReC), where it was processed to obtain viable seeds. The extracted seeds were surface-sterilized and germinated under controlled conditions. The extracted seeds were surface-sterilized and germinated under controlled conditions. The first pair of cotyledon leaves was harvested for genomic DNA extraction using a modified cetyltrimethylammonium bromide (CTAB) method (1.4 M NaCl, 3% (v/v) β-mercaptoethanol, 20 mM EDTA, 3% (w/v) CTAB, and 100 mM Tris-HCl, pH 8.0). DNA concentration and purity were assessed using a NanoDrop spectrophotometer, and integrity was verified by electrophoresis on a 1% agarose gel. High-quality, high-molecular-weight DNA was then shipped to Macrogen (Netherlands) for library preparation and sequencing. Paired-end sequencing was performed on the Illumina MiSeq platform to generate raw reads for downstream genome assembly.

### Genome assembly and annotation

Raw reads were quality assessed in Fastp^33^, with parameters set to default. Reads with low base quality (Phred score < 25), reads containing more than 20% low-quality bases, and reads shorter than 80 bp were removed. High-quality reads were processed and passed for *de novo* assembly in the GetOrganelle toolkit v1.7.7.1^34^, using SPAdes v3.15.4 ^35^ algorithm. Assembly parameters were set as described in ^36^. The complete assembled circular genome was visualized in Bandage v1.8.7^37^, identifying assembly coverage for each segment. Annotation of the genome was performed using two independent approaches. First, annotation was performed using Plastid Genome Annotator (PGA)^38^ in PlastidHub^39^, using *Arabidopsis thaliana* (NC_000932) as the reference genome with parameters set to default. Second, a *de novo* annotation was conducted using the GeSeq web application^40^, with third-party annotating tools tRNAscan-SE v2.0.7^41^, ARAGORN v1.2.38^42^, and Chloë v0.1.0 (https://github.com/ian-small/Chloe.jl) all activated. *BLAT* search for protein identity was set at 85%, while tRNA and rRNA search identities were set at 95%. The resulting GenBank files from PGA and GeSeq platforms were processed in GB2sequin^43^ to produce NCBI-compliant submission files using the NCBI table2asn^44^ program. Annotated genes from the two platforms were manually screened to identify potential mis-annotations and ensure consistency. The final curated GenBank file was loaded to OGDRAW v1.3.1^45^ to generate a circular graphical representation of the assembled *A. digitata* chloroplast genome.

### Repetitive sequence, codon usage analysis, and prediction of RNA editing sites

Simple sequence repeats (SSR) were identified using CPSTools^46^ with parameters set to -k 10,5,5,3,3,3 (≥C:10 repeats for mononucleotide,C:≥C:5 for dinucleotide, ≥C:4 for trinucleotide, and ≥C:3 for tetranucleotide, pentanucleotide, and hexanucleotide motifs). Long sequence repeats (LSRs) were identified using REPuter^47^ (repfind -c -f -p -r -l 30 -h 3 -best 50), with a minimum repeat length of 30 bp, a Hamming distance of 3, and repeat types specified as forward (F), reverse (R), palindromic (P), and complement (C). Relative synonymous codon usage (RSCU) was calculated using CPStools v3.0^46^, and the codon usage patterns in protein-coding genes were analysed by extracting all annotated coding DNA sequences from the chloroplast genome GenBank file using the RSCU function in CPStools^46^. The extracted sequences were converted to FASTA format, and only complete CDSs with valid start and stop codons and no internal stop codons were retained for codon usage analysis. The resulting CDS dataset was used as input for CodonW v1.4.2^48^ to compute codon usage bias and nucleotide composition indices. The computed indices included base frequencies at synonymous third codon positions (T3s, C3s, A3s, G3s), codon adaptation index (CAI), codon bias index (CBI), frequency of optimal codons (Fop), effective number of codons (Nc and eNC), GC content at synonymous third positions (GC3s), overall GC content (GC), number of synonymous codons (L_sym), number of amino acids (L_aa), hydropathicity (Gravy), aromaticity (Aromo), nucleotide bias indices (A3_bias, G3_bias, GC12, AT_bias, GC_bias), and ENC deviation metrics (ENC_diff_ratio). RNA editing site prediction in protein-coding genes was carried out using PREPACT 3.0. The chloroplast genome of *A. thaliana* (NC_000932) was used as the reference sequence for BLASTx-based prediction. Both forward (C→U) and reverse (U→C) editing directions were selected, and all remaining parameters were retained at their default settings.

### Comparative, collinearity and average nucleotide identity analyses

The assembled chloroplast genome was queried against the NCBI database (accessed on October 26, 2025) using BLASTn v2.17.0^50^ to identify homologous genome sequences. Complete chloroplast genomes from nine Adansonia species showing >99% sequence similarity were retrieved for comparative analysis. These included *A. za* (MT053006.1), *A. grandidieri* (MT052999.1), *A. gregorii* (MT053000.1), *A. madagascariensis* (MT053002.1), *A. rubrostipa* (MT053004.1), *A. kilima* (MT053001.1), *A. suarezensis* (MT053005.1), and *A. perrieri* (MT053003.1). All sequences were downloaded in GenBank format. Genome structural features and nucleotide composition were computed for each chloroplast genome using in-customized bash and R scripts. Contraction and expansion of inverted repeat (IR) regions and the junctions among LSC, IRb, SSC, and IRa were visualized using IRscope^51^. Multiple sequence alignment of the chloroplast genomes was performed in MAFFTv7.525^52^ with default parameters. Nucleotide diversity (Pi) across aligned genomes was estimated using CPStools^46^ with default settings. Collinearity analysis was performed using pgDRAW (https://gbdraw.app/) with default settings, enabling runLOSAT and BLASTN options. Average Nucleotide Identity was calculated using the JSpeciesWS web server (https://jspecies.ribohost.com/jspeciesws/) based on BLAST-based ANI comparisons among the chloroplast genome sequences.

### Analysis of selection pressure on protein-coding genes

Protein-coding sequences (CDSs) were extracted from GenBank files of *Adansonia* species using CPStools^46^. The sequences were then screened to identify incomplete genes, sequences containing internal stop codons, and spurious sequences, all of which were removed using customized Python scripts. The validated CDSs were aligned using MAFFT v7.525^52^ and back-translated with PAL2NAL. Final alignments were refined using MACSE^53^ to ensure accurate codon-based alignment. Substitution rates were calculated using the codeML module of EasyCodeML v1.41^54^, with other settings kept at default. The resulting dN/dS values were processed using a customized Python script and exported as a CSV file for downstream analyses in R v4.5.0^55^, using a suite of tidyverse packages v2.0.0^56^.

### Technical validation of the assembled genome through phylogenomic analysis

Complete chloroplast genomes of other species in the family *Malvaceae* were retrieved from the National Center for Biotechnology Information (NCBI; https://www.ncbi.nlm.nih.gov/, accessed 11 November, 2025) using the search query: (“Malvaceae”[Organism] OR malvaceae[All Fields]) AND chloroplast[All Fields] AND genomes[All Fields]. For species with multiple chloroplast genome versions, only the most recent, complete, and fully annotated genome was selected for download. The sequences were downloaded in FASTA format and merged with the assembled *A. digitata* genome into a single file. Sequence alignment was performed in MAFFT v7.525^52^ with parameters set to default. Maximum likelihood phylogenetic trees were reconstructed using IQ-TREE^57^, and Bayesian phylogenetic analyses were conducted with MrBayes v3.2^58^. The trees were annotated in iTOL v6^59^.

## Results

### Features of the assembled genome

The assembled chloroplast genome of *A. digitata* exhibited the canonical quadripartite circular structure characteristic of angiosperm plants, with a total length of 160,061 bp (Fig. 1). The genome comprised a large single-copy (LSC) region of 88,961 bp, a small single-copy (SSC) region of 25,553 bp, and two inverted repeat regions (IRa and IRb), each 19,994 bp in length. Sequencing coverage varied within the genomic regions, with the LSC, SSC, and IR regions having average assembly depths of 128×, 231×, and 132×, respectively (Supplementary Fig. S1). The overall GC content of the chloroplast genome was 36.87%, with noticeable variation among genomic regions and between coding and non-coding sequences. Within the structural regions, the IR regions showed the highest GC content (42.92%), whereas the SSC and LSC regions showed comparatively lower GC levels of 34.68% and 31.17%, respectively. Within the coding regions, tRNA and rRNA genes showed the highest GC contents, while protein-coding genes and intronic regions exhibited relatively lower GC contents (Supplementary Table S1).

**Figure 1:**
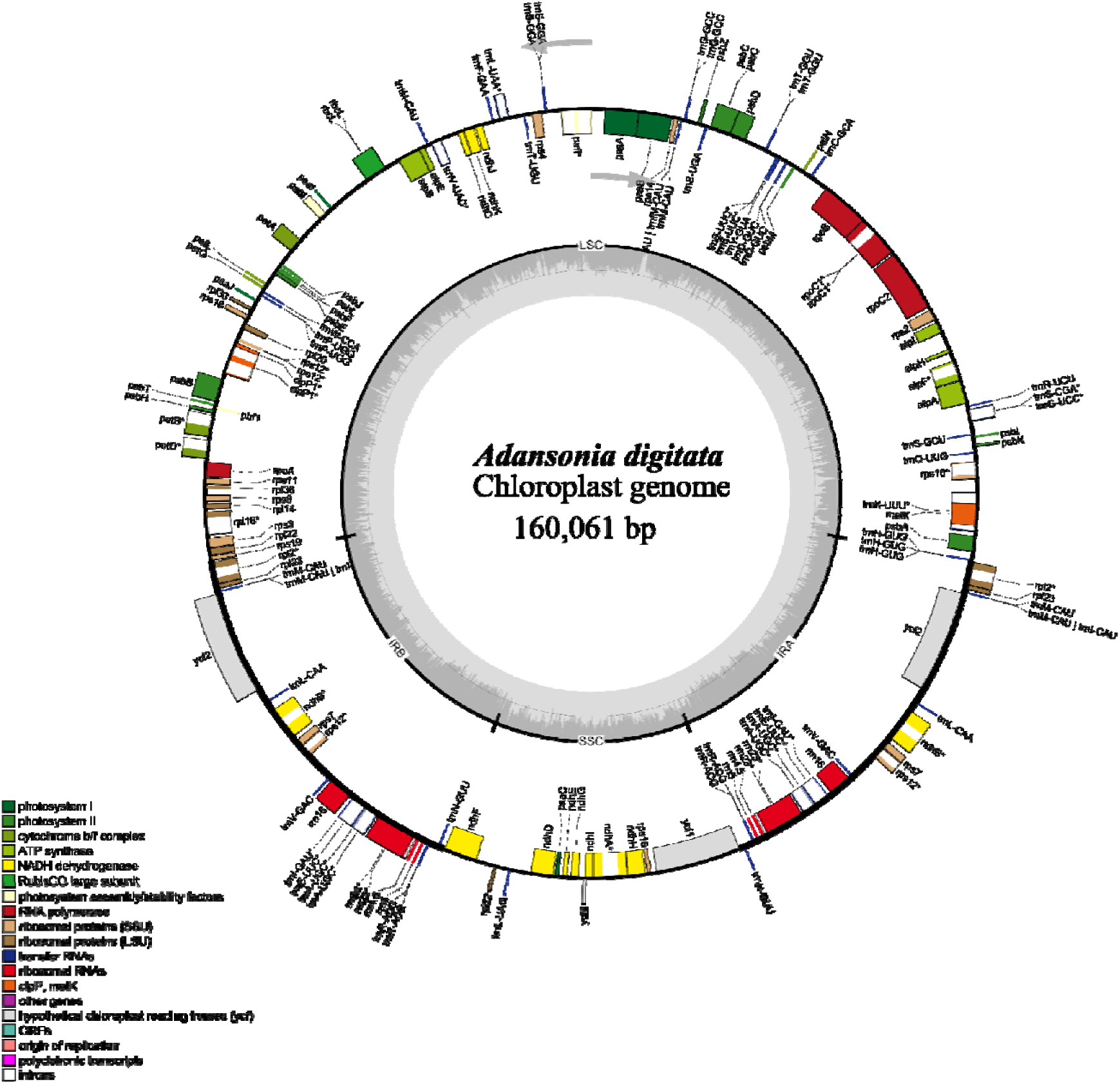
Circular map of assembled *A. digitata* chloroplast genome. The dark grey region within the circle represents the distribution of GC content across the genome, while the light grey region indicates AT content. Genes located on the inside of the circle are transcribed in the clockwise direction, whereas genes on the outside are transcribed in the anticlockwise direction, as indicated by the arrows. Gene features are colour-coded according to their functional categories.

A total of 115 genes were predicted in the *A. digitata* chloroplast genome. These comprised 79 protein-coding genes, 32 transfer RNA (tRNA) genes, and 4 ribosomal RNA (rRNA) genes (Supplementary Table S2). The genes were distributed across the four structural regions of the genome: 85 in the large single-copy (LSC) region, 13 in the small single-copy (SSC) region, and 18 and 19 in the inverted repeat regions IRa and IRb, respectively. Notably, the IRb region contained an additional copy of the *ycf1* gene. The remaining 30 genes were duplicated and located within the LSC, IRa, and IRb regions (Table 1). Among the annotated genes, nine protein-coding genes and nine tRNA genes each contained a single intron, producing two exons per gene. Three genes, *clpP1* and *pafI* (protein-coding), and rps12 (ribosomal protein), contained two introns (Supplementary Table S2). In addition, two genes, *infA* and *ndhD,* were identified to initiate translation with non-canonical start codons, ATT and ACG, respectively (Table 1).

**Table 1:**
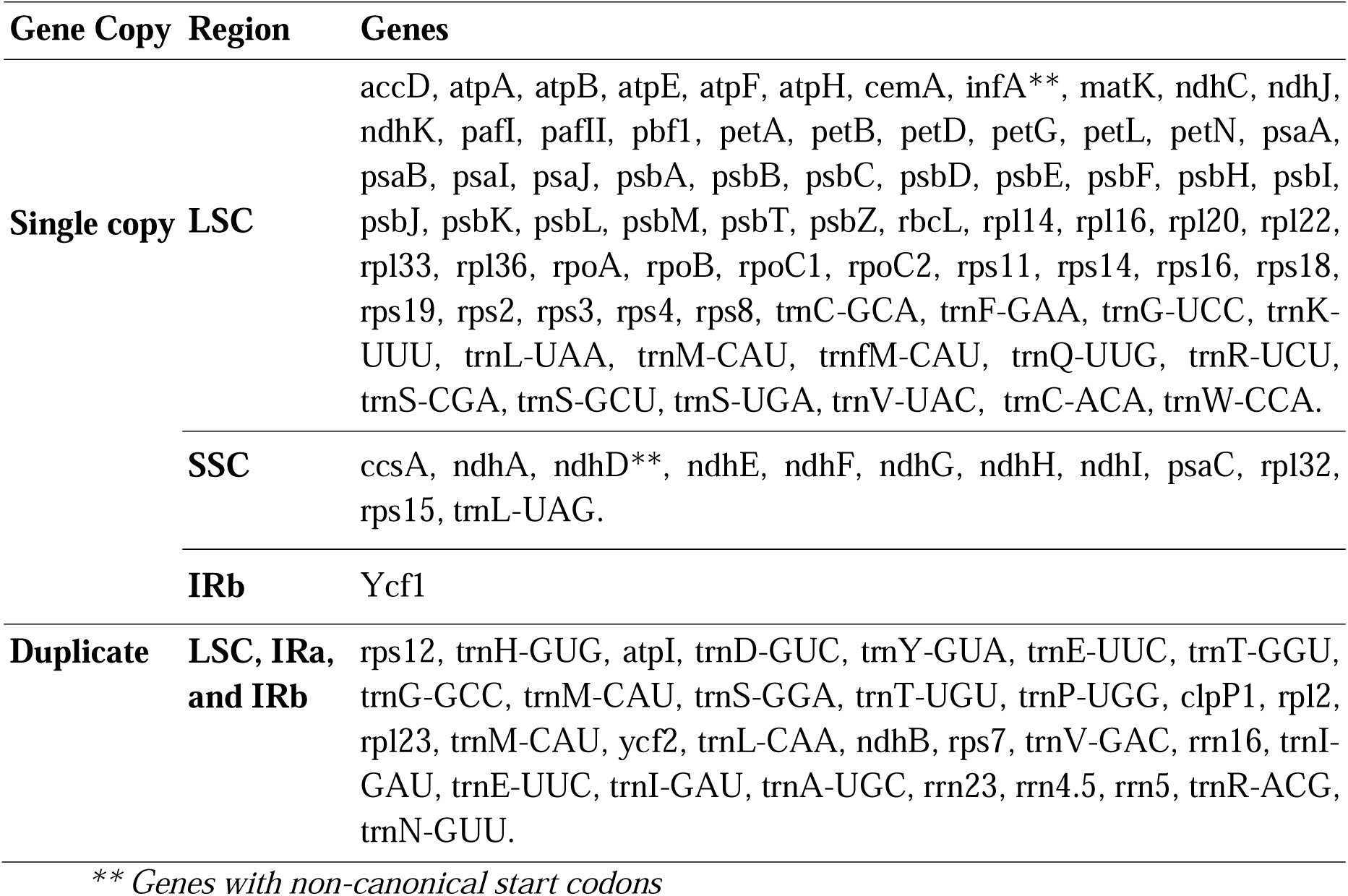
Genes identified in each structural region of the *A. digitata* chloroplast genome.

### Comparative structural and compositional analysis

Comparative analysis of the assembled *A. digitata* chloroplast genome with those of eight recognized *Adansonia* species and the putative *A. kilima* revealed high structural similarity across all analytical frameworks (Supplementary Table S1). All genomes exhibited the typical quadripartite structure comprising an LSC region, an SSC region, and a pair of IR regions. Genome sizes ranged from 159,710 to 160,621 bp, with the *A. digitata* having a genome of 160,061 bp long. The LSC region of *A. digitata* (88,961 bp) was comparable to those of *A. grandidieri, A. kilima*, and *A. gregorii.* Its SSC region (20,016 bp) was within the size range for most species, except *A. perrieri*, which exhibited a shorter SSC. IR lengths were highly conserved across species; *A. digitata* had a combined IR length of 51,106 bp, consistent with the other *Adansonia* chloroplasts. The protein-coding portion of the *A. digitata* genome spanned 84,624 bp, closely matching those of *A. suarezensis* and *A. kilima* and was slightly shorter than those of *A. grandidieri* and *A. gregorii*. The lengths of rRNA and tRNA coding regions were similar across the species. Total intron length in *A. digitata* was 31,012 bp, similar to most *Adansonia* species, with the exception of *A. gregorii*, which showed a reduced intron length. Overall GC contents across *Adansonia* chloroplast genomes ranged from 36.78% to 36.90%. *A. digitata* had a GC content of 36.87%, closely similar to values reported for *A. grandidieri*, *A. kilima,* and *A. za*. GC distribution followed a consistent regional pattern across all species, with the IR regions having the highest GC content, followed by the LSC region, whereas the SSC had the lowest GC content observed (Supplementary Table S1). Gene number and composition were highly conserved throughout the genus. All genomes contained 79 unique protein-coding genes, with total copy numbers varying between 95 and 97 protein-coding gene copies. *A. digitata* contained 79 protein-coding genes (95 total copies), 32 tRNA genes (66 copies), and 4 rRNA genes (10 copies), consistent across the genus. A conserved set of 22 genes contained introns, and the total intron count was stable at 43 across species, except for *A. gregorii,* which contained 41 introns (Supplementary Table S1). The assembled *A. digitata* genome showed concordance with the structural, compositional, and gene content characteristics observed across the genus.

### Collinearity and Average Nucleotide Identity analysis

Collinearity analysis revealed a high degree of similarity in chloroplast genome structure among the *Adansonia* species, with no notable large structural variation in genome organization or gene order (Supplementary Fig. S2a and S2b). Average nucleotide identity values between *Adansonia* species were relatively high, ranging from 99.28% to 99.96% among the species (Fig. 2). The highest pairwise ANI was observed between *A. digitata* and *A. kilima* (99.96%). High similarity was also observed among the Malagasy species, particularly between *A. madagascariensis* and *A. perrieri* (99.80%), and between *A. rubrostipa* and *A. za* (99.79%). *A. gregorii* showed consistently low ANI values relative to other species (approximately 99.28 - 99.40%). *A. suarezensis* showed relatively high similarity across multiple pairwise comparisons, up to 99.62% with *A. grandidieri* and 99.61% with *A. digitata*.

**Figure 2:**
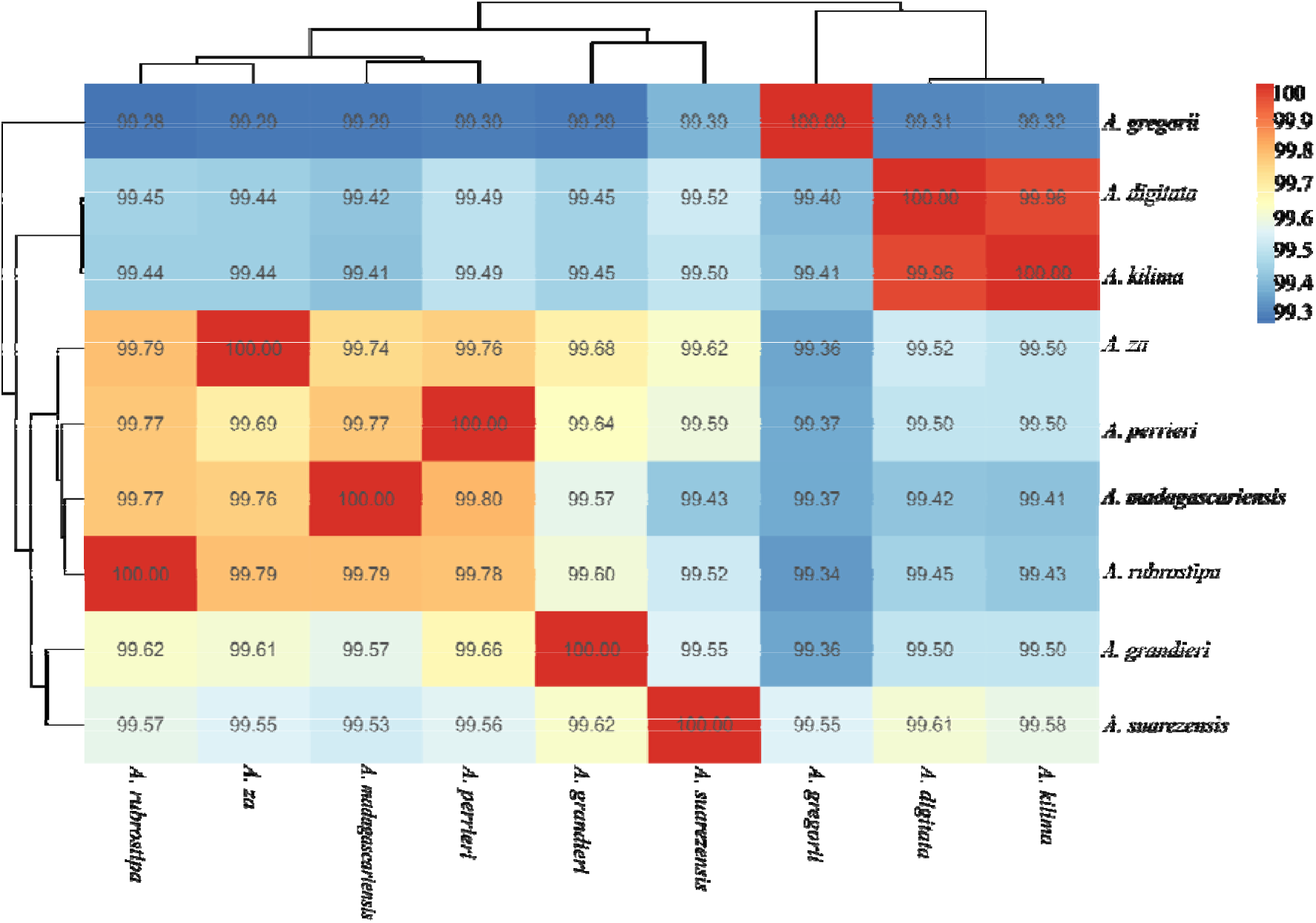
Comparative average nucleotide identity (ANI) among the eight known *Adansonia* species and putative *A. kilima*.

### Comparative analysis of IR boundaries

The positions of the LSC, SSC, and IR boundaries showed minor variation among the *Adansonia* chloroplast genomes, with most differences involving contraction and expansion of the *ycf1* and *ndhF* genes. The JLB (LSC/IRb) and JLA (LSC/IRa) junctions were generally consistent across species. At the JLB boundary, the flanking genes were *rpl22*, *rpl2*, and *rps19*, with *rps19* extending 5 bp into IRb in all species (Fig. 3). At the JLA boundary, the flanking genes consisted of *rpl2* in IRa and *psbA* in the LSC, with *trnH* located at the border, extending 2 bp into IRa in all *Adansonia* species. At the JSA boundary, the nine genomes showed no variation in gene arrangement or position; however, variation was observed in the length of the *ycf1* gene. The *ycf1* gene length was consistent in most *Adansonia* species, except in *A. gregorii*, where it was 5,681 bp long, with 5,647 bp located in the SSC region and 34 bp in the IRa region. In the other species, *ycf1* extended 34 bp into IRa and 5,672 bp into the SSC region. Differences were also observed at the JSB (IRb/SSC) boundary. In *A. perrieri* and *A. grandidieri*, *ycf1* was duplicated across the JSB boundary, with a shared 34 bp segment in IRb and shorter SSC-side fragments of 71 bp and 57 bp, respectively. The remaining portion of *ycf1* in both species was located in the SSC region, with a conserved length of 5,672 bp, and a corresponding 34 bp fragment extending into IRa at the JSA boundary. Notably, the *ndhF* gene also varied in its relative position from the JSB boundary, ranging from 50 to 113 bp among species (Fig. 3).

**Figure 3:**
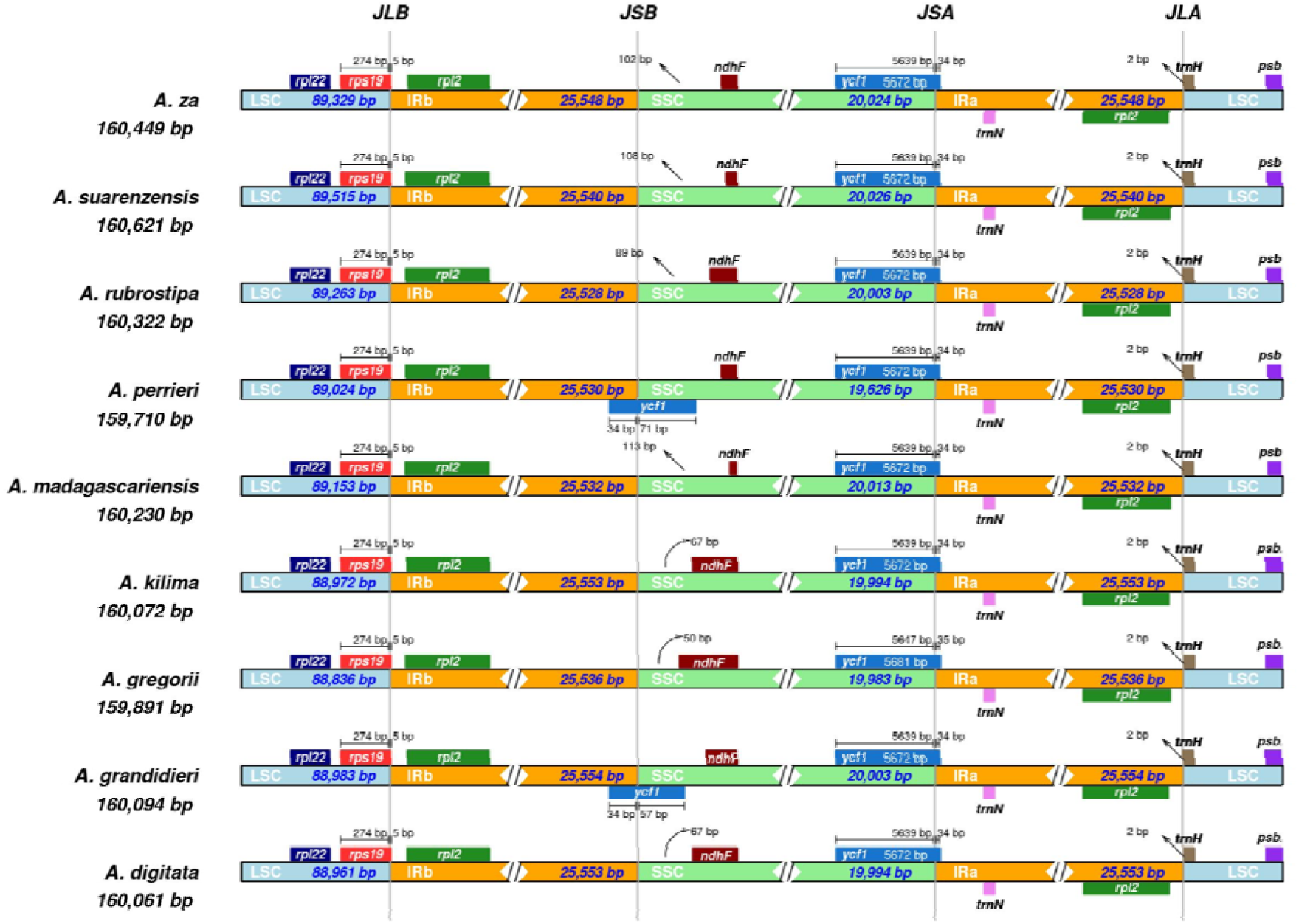
Contraction and expansion patterns at the JLB (LSC/IRb), JLA (LSC/IRa), JSB (IRb/SSC) and JSA (SSC/IRa) junctions among chloroplast genomes of *Adansonia* species, showing variation in boundary positions and gene arrangements across taxa.

### Long and Simple sequence repeats analysis

A total of 50 repeats (≥30 bp) were detected, majority of which were forward (26) and palindromic (17), whereas reverse and complementary repeats occurred less frequently (Supplementary Table S3). Repeat lengths ranged from 30 to 49 bp, with the majority (70%) having lengths between 30 and 36 bp. The longest non-IR dispersed repeats were 49 bp, followed by repeats of 46-43 bp, all of which showed low divergence and strong statistical support. Mismatch values across all repeats ranged from 0 to 3, with most palindromes and forward repeats (54%) showing zero or one mismatch. The statistical significance of detected repeats was high, with E-values ranging from 0.00 to 5.23 × 10[[. The repeats were distributed across the large single-copy (LSC), small single-copy (SSC), and inverted repeat (IR) regions, with a number of repeats located near IR boundaries (Supplementary Table S3). Analysis of simple sequence repeats (SSRs) across the *A. digitata* chloroplast genome identified a total of 100 sets of microsatellite motifs distributed in the coding, intronic, and intergenic regions (Supplementary Table S4). The detected SSRs comprised mono-, di-, tri-, tetra-, and pentanucleotide motifs, with the dominance of mononucleotide repeats (76%), most of which consisted of A/T bases. Dinucleotide repeats accounted for 12.12%, whereas only a single trinucleotide motif, 10 AT-rich tetranucleotide repeats, and one pentanucleotide repeat were identified. SSR motif lengths ranged from 3 to 17 bp, with the majority of the repeats (74%) having lengths between 10 and 14 bp. The longest repeat detected was a 17 bp poly-A motif, located within the pafI-trnS-GGA intergenic spacer, while several long poly-T motifs (≥14 bp) were also identified in non-coding regions, particularly within intergenic spacers associated with photosynthesis- and transcription-related genes. SSRs were unevenly distributed within the genome, with the intergenic spacers having the highest proportion of repeats (73%). Comparatively fewer (11%) SSRs were detected within the coding genes (Supplementary Table S4). Gene-associated SSRs were identified in *matK*, *atpI*, *rpoC2*, *rpoC1*, *rpoB*, *psaA*, *rbcL*, *cemA*, *ycf1*, and *ndhF*. Among intronic regions, SSRs were most frequently detected in genes containing large or multiple introns, such as *clpP1*, *rpl16*, and *ndhA*. Specific intergenic regions exhibited high clusters of SSRs, particularly those flanked by *trnK-rps16*, *atpF-atpH*, *trnE-trnT*, *trnT-trnL*, *psaJ-rpl33*, and *ycf1* (Supplementary Table S4).

### Prediction of RNA editing sites

Prediction of RNA editing sites identified 578 sites across 63 protein-coding genes (Supplementary Table S6). The edits involved conversion of C to U, and vice versa, with the C to U conversion constituting the highest proportion (55.02%). All predicted RNA editing events resulted in non-synonymous codon changes. The substitutions were limited to the first and second codon positions, with edits at the second position accounting for 54.17%, while none were observed at the third codon position. In addition to single-site edits, 31 cases were identified in which two RNA editing events occurred within the same codon, producing combined codon changes. These were detected in *accD, ndhE, ysfl1, rpl12,* and *ccsA*. The number of predicted editing sites varied among genes. The highest proportions of editing sites were recorded in *ycf1* (20.09%), *rpoC2* (11.51%), *ycf2* (8.12%), and *matK* (6.99%), as well as across multiple *ndh* genes, with additional edits detected in several other genes (Supplementary Table S6). No RNA editing sites were predicted in 21 protein-coding genes, namely *psaC*, *rps7*, *rpl2*, *rps19*, *rps3*, *rpl16*, *rpl36*, *clpP*, *rps12*, *rpl33*, *psbE*, *psbL*, *atpE*, *ycf3*, *psbC*, *psbD*, *psbM*, *petN*, *atpH*, *psbI*, and *psbA*. The amino acid change profile showed a concentration of substitutions among hydrophobic and polar residues, with a number of rare and stop-related conversions. Predicted amino acid substitutions comprised 26 substitution classes. The most frequent substitutions were phenylalanine to leucine (F→L) and serine to leucine (S→L), each recorded 55 times, followed by leucine to phenylalanine (L→F, 40) and threonine to isoleucine (T→I, 34). Other common substitutions, along with the reverse conversions of arginine to cysteine (R→C, 2) and arginine to tryptophan (R→W, 2), were detected less frequently (Supplementary Table S6). Predictions for stop codon editing were also detected, with stop to arginine (*→R, 2) in *rps16* and *rpl22*, and glutamine (Q→*, 1) in *rpoC2*.

### Codon usage analysis

Relative synonymous codon usage (RSCU) analysis of the protein-coding genes identified heterogeneous patterns of codon usage bias across the genome. RSCU values ranged from 0.34 to 2.09 (Fig. 4), indicating non-random codon selection. Tryptophan (Trp) and methionine (Met) were each encoded by a single codon (TGG and ATG, respectively), whereas all other amino acids were represented by two to six synonymous codons. A total of 26 codons showed preferential usage (RSCU > 1.3), with the highest values observed for leucine (TTA), arginine (AGA), and alanine (GCT). Fifteen codons displayed approximately neutral usage (RSCU 0.68–1.28), while the remainder were underrepresented (Fig. 4).

**Figure 4:**
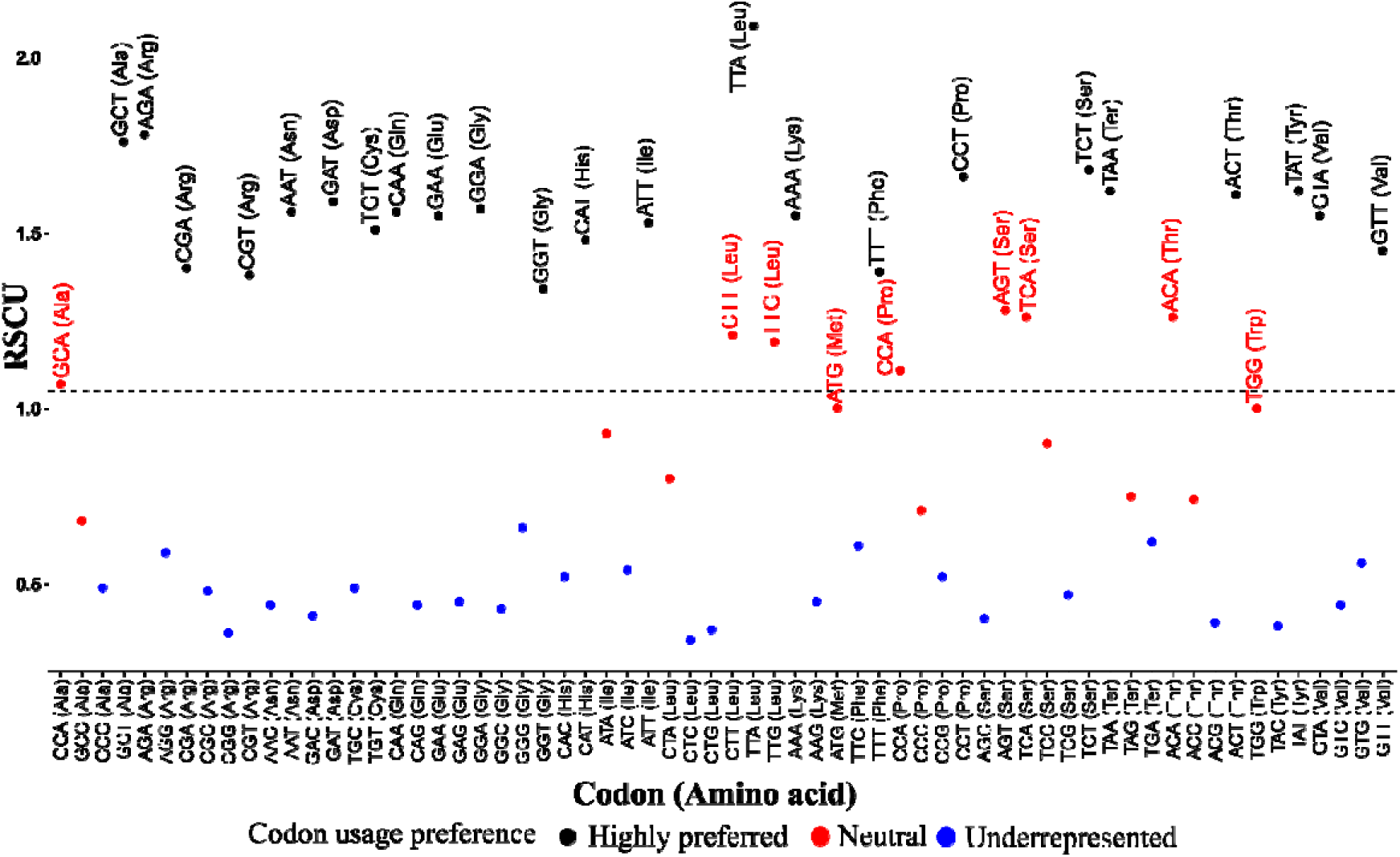
Relative synonymous codon usage (RSCU) of protein-coding genes in the *A. digitata* chloroplast genome. Codons are shown on the x-axis, with their corresponding amino acids indicated in parentheses. Codons with usage bias are labeled with the corresponding amino acid abbreviation.

### Codon usage bias in protein-coding sequences

The effective number of codons (Nc) ranged from 36.33 to 56.65, while GC3 values varied between 0.18 and 0.32. The Nc-GC3 analysis showed that codon usage bias across the protein-coding genes was generally low to moderate, with most genes having GC[ values ranging from 0.18 to 0.32, and Nc values of approximately 45-52 (Fig. 5a). Most genes clustered near the expected neutrality curve. This was supported by the Kolmogorov-Smirnov test, which revealed no significant difference between the distributions of Nc and eNC (D = 0.224, p = 0.165), with the number of codon cumulative distribution frequency being slightly higher than the eNC CDF (Fig. 5b). Distinct gene-level deviations were identified in *psbA*, *rps14*, *rps8*, and *rps18*, which showed strong codon preference, characterized by low Nc values (36.33 - 39.40) and high positive Nc deviation ratios (0.2266-0.3166), placing them well below the neutrality curve (Fig. 5a). Other genes, including *atpF*, *cemA*, *psbD*, *petD*, and *rpl16*, also showed moderate deviation ratios. Genes such as *clpP1*, *pafI*, and *ndhJ* had high Nc values (55.51 - 56.65) and negative deviation ratios (−0.0836 to −0.1200), positioning them above the neutrality curve, indicating weak codon preference (Fig. 5a). The frequency distribution of eNC deviation ratios showed concentration around zero, with a limited number of genes forming upper and lower tails corresponding to elevated and reduced codon bias, respectively (Fig. 5c).

**Figure 5.**
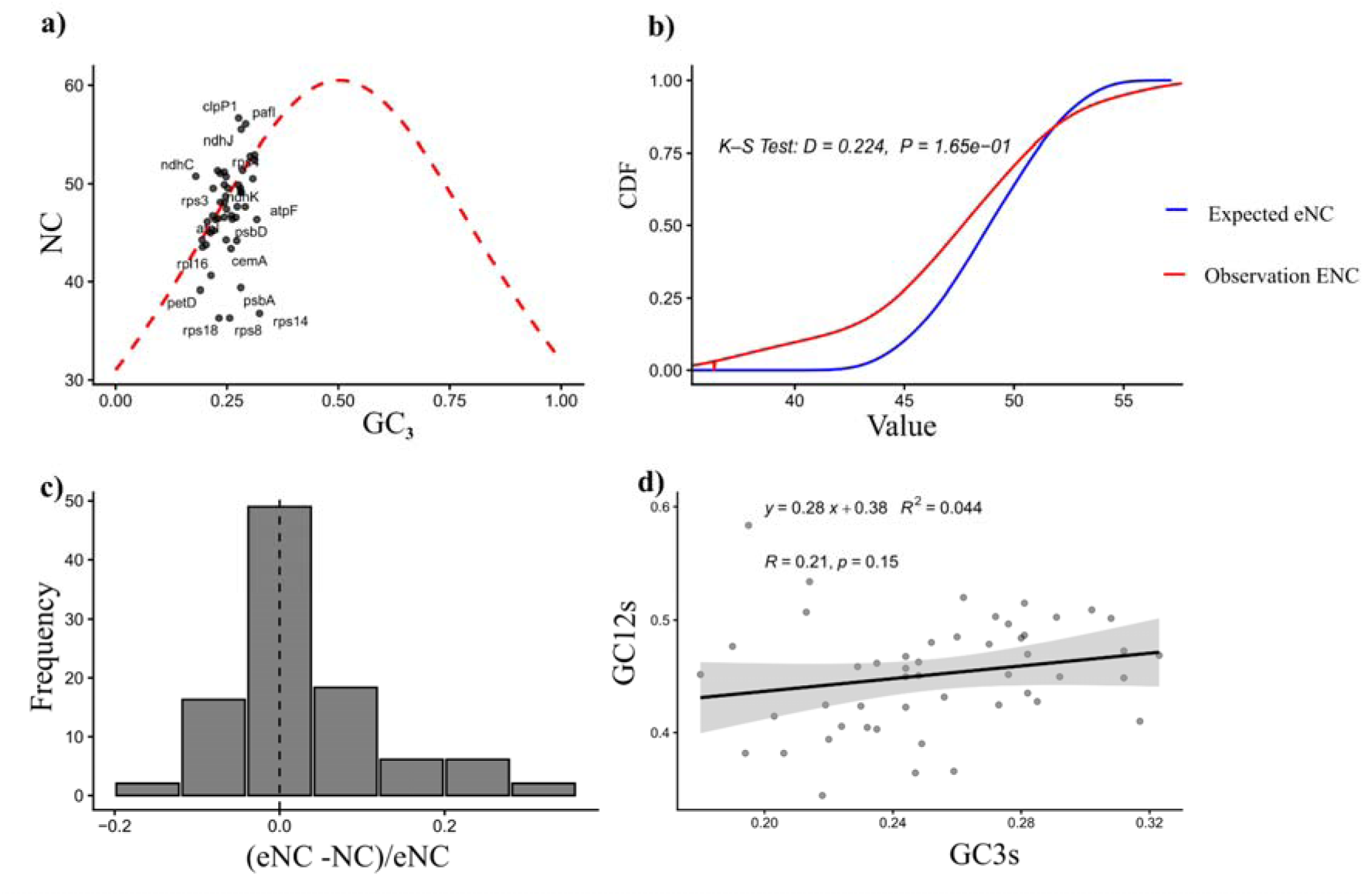
Codon usage bias in protein-coding sequences. (**a**) Nc–GC3 plot illustrating codon usage bias across genes. (**b**) eNc plot comparing expected and observed codon bias. (**c**) Frequency distribution of eNc values. (**d**) Relationship between GC content at the third codon position and the first and second positions across all genes.

### Frequency of optimal codons and neutrality analysis

Evaluation of the frequency of optimal codons (Fop) revealed variation among protein-coding genes. The highest Fop values were observed in psbA, rbcL, and psbD (Supplementary Table S5). Moderate Fop values were detected in atpB, petB, cemA, atpE, and rps7. A number of NADH dehydrogenase genes, ndhG, ndhF, ndhE, ndhA, and matK, had lower Fop values, indicating reduced optimal codon usage relative to other genes in the genome (Supplementary Table S5). Neutrality analysis based on GC content across coding regions showed variation in GC values at the first and second codon positions (GC12) and at the third codon position (GC3s) among the examined genes. GC12 values ranged from 0.344 in ycf1 to 0.5835 in rps11, with most genes showing GC12 values between 0.40 and 0.52. GC3s values were consistently lower, ranging from 0.18 in ndhC to 0.323 in rps14. Genes such as rps11, rpl16, and psbB showed the highest GC12 values, whereas ycf1, accD, and cemA had the lowest. For GC3s, the highest values were recorded in rps14, atpF, and atpE/petA, while the lowest GC3s values were observed in ndhC, ndhF, and petD. Overall, GC content was consistently higher at GC12 than at GC3s across all genes analyzed (Fig. 5d). Linear regression analysis indicated a weak positive correlation between GC3s and GC12 (R = 0.21, R² = 0.044, P = 0.15).

### Parity Rule 2 and correspondence analysis of codon bias in protein-coding sequences

Based on Parity Rule 2 analysis, the average A3/(A3+T3) ratio was 0.481 ± 0.05, lower than the expected value of 0.5, indicating a moderate preference for T over A. Similarly, the average G3/(G3+C3) ratio was 0.505 ± 0.09, indicating a moderate bias toward G over C (Fig. 6a). Correspondence analysis (COA) of codon usage generated 48 axes, of which the first two accounted for a major portion of the variation (27.99%). Axis 1 explained 17.48% of the total variation, while Axis 2 explained 10.51% (Fig. 6b). Codons with high positive scores on Axis 1 were predominantly A- and G-ending codons (AAA, AAG, AGA, AGG, CGA, and CGG), whereas strongly negative Axis 1 scores were mainly associated with U-and C-ending codons (GCU, GGU, GGC, GCC, and UUC). Axis 2 structured codon distribution by separating several G- and C-ending codons with high positive loadings (e.g., CGC, CGG, AGG, CGU) from U- and A-ending codons with negative loadings (e.g., UUU, UCU, UCA, UUA).

**Figure 6:**
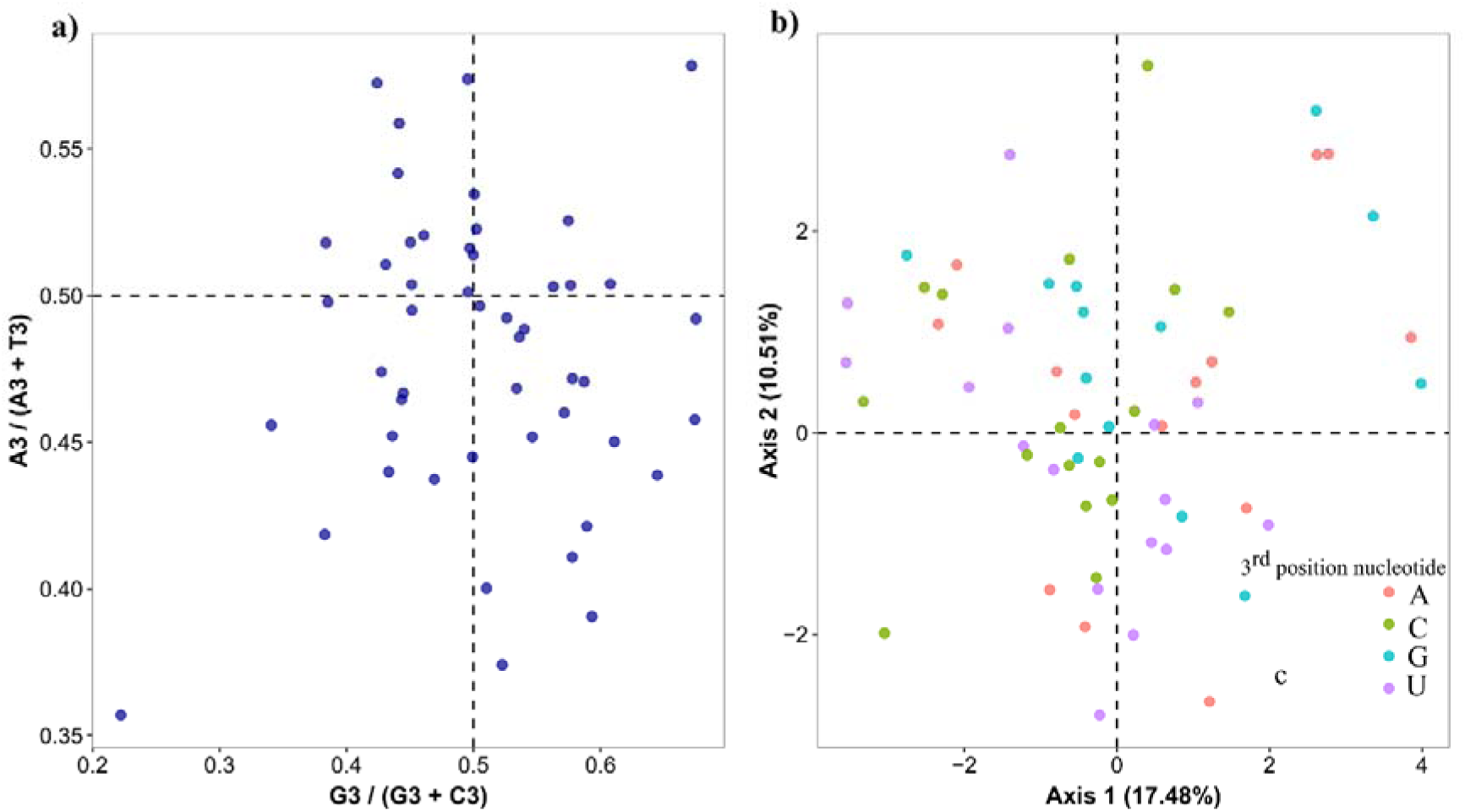
Codon usage bias in protein-coding sequences of *A. digitata*. (a) Parity Rule 2 analysis showing the average ratios of A/(A+T) and G/(G+C) at the third codon position. (b) Correspondence Analysis of relative synonymous codon usage (RSCU) illustrating the distribution of codons across the first two principal axes.

### Nucleotide diversity (π) analysis

Nucleotide diversity (π) varied across the coding and intergenic regions of the genome. Coding sequences were largely conserved (0.00167±0.00199), with most genes showing π values below 0.001 or complete invariance. Five genes, *infA, rps16, rpl33, ycf1*, and *ndhI*, were identified as polymorphic hotspots, while moderate levels of variation were observed in rbcL, petL, ndhD, and ndhH (Fig. 7a). Intergenic spacers showed greater variability (0.00490±0.00574), with several loci having relatively higher π values (Fig. 7c). Highly variable regions included trnM-CAU in the IR, trnG-UCC[-trnR-UCU, petD-rpoA, trnH-GUG-psbA, and rpl2[-trnH-GUG. Several spacer regions were completely conserved (π = 0). Nucleotide diversity was unevenly distributed across the genome, with most variation concentrated in intergenic regions of the LSC, whereas the IR regions and the majority of coding genes remained comparatively conserved (Fig. 7b).

**Figure 7:**
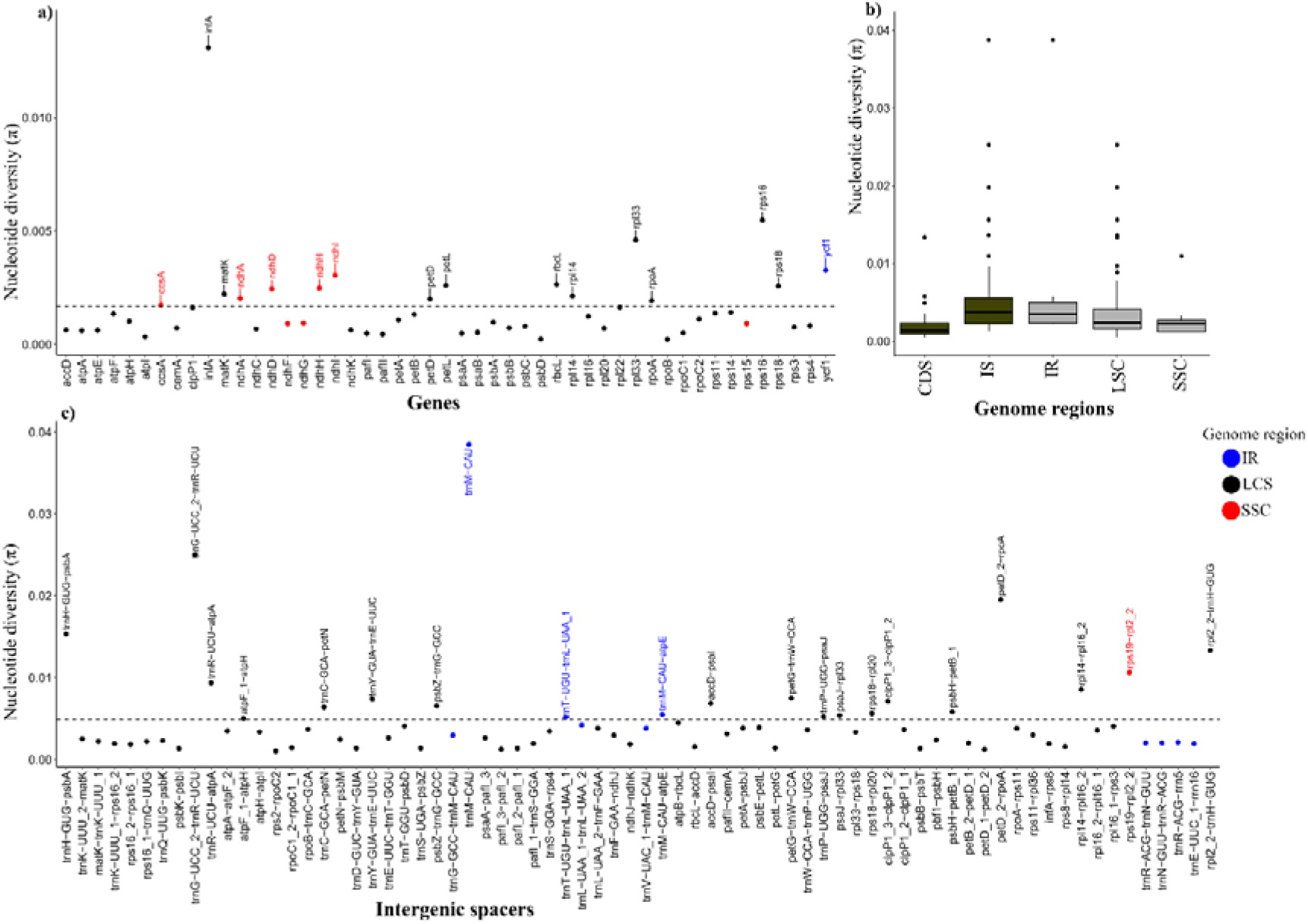
Nucleotide diversity (π) of the *A. digitata* chloroplast genome. (a) across protein-coding genes, (b) average diversity in coding (CDS), intergenic spacers (IS), and genome regions (IR, LSC, SSC), (c) in intergenic and intronic regions. Genes and intergenic regions with nucleotide diversity of less than 0.001 were not represented in graphs a and c, but were included as part of graph b.

### Substitution rates and selection pressure

Estimates of the substitution rate ratio, Kappa (κ), and the nonsynonymous to synonymous substitution ratio (ω = dN/dS) were obtained for all protein-coding genes in *A. digitata* chloroplast genome. The κ values showed wide variation among genes, ranging from 0.0001 in cemA, rps18, rpl20, ndhG, rps11, ndhK, psaB, rpl14, rpl22, pafII, petB, psbD, psaA, atpH, atpE, petA, rps15, ndhC, rps14, and petL, to a maximum of 7.64777 in rps19. Higher κ values were also observed in psaC and rps16 (Fig. 8). The ω values ranged from 0.0001 in multiple genes (rps19, psbB, rpl16, psbA, clpP1, psbC, rpoA, rps18, rpl20, ndhG, rps11, ndhK, psaB, rpl14, rpl22, pafII, petB, psbD, psaA, atpH, atpE, petA, rps15, ndhC, rps14, and petL) to 0.89043 in matK (Fig. 8). Relatively higher ω values were also observed in ycf1, accD, and rpoB. Most genes, particularly those associated with photosystem I and II, ribosomal structure, and NADH dehydrogenase functions, had ω values below 0.5.

**Figure 8:**
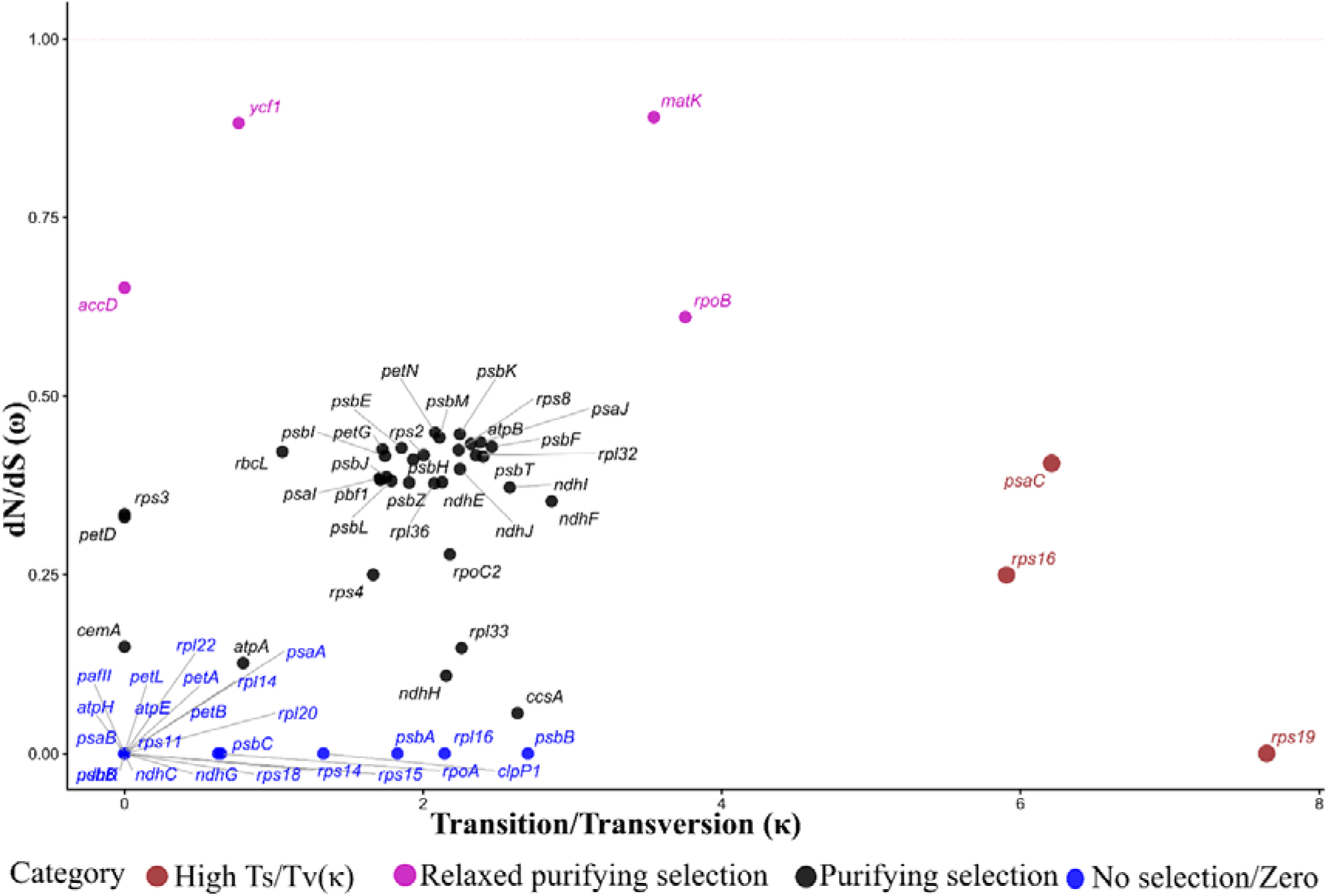
Scatter plot of ω (dN/dS) versus κ (Transition/Transversion) for protein-coding genes in the *A. digitata* chloroplast genome. Genes shown in pink exhibit high dN/dS(ω) values, suggesting relaxed or positive selection, while genes in red have high Transition/Transversion (κ) values, indicating increased transition rates relative to transversions.

### Phylogenetic analysis

A total of 900 chloroplast genomes from the family *Malvaceae* were downloaded from NCBI. Following the removal of duplicate entries and outdated assemblies, 248 high-quality, contiguous genomes were retained for phylogenomic reconstruction. Maximum likelihood (ML) and Bayesian inference (BI) analyses produced congruent topologies, consistently placing the assembled *A. digitata* plastome within the *Adansonia* lineage (Fig. 9). In all the analyses, A. digitata formed a monophyletic clade with other *Adansonia* species, having full posterior probability support (PP = 1.0) in the Bayesian tree and maximal bootstrap support (BS = 100) in the ML tree for the broader *Adansonia* clade and the subclade containing A. digitata. At the family level, *Adansonia* resolved as a well-supported, distinct clade within Malvaceae, clearly separated from other genera such as *Fremontodendron, Ceiba, Ochroma,* and *Firmiana*. Within the genus, the assembled *A. digitata* plastome was most closely related to the putative *A. kilima* and *A. gregorii*, a placement that was consistent across the two analytical frameworks.

**Figure 9.**
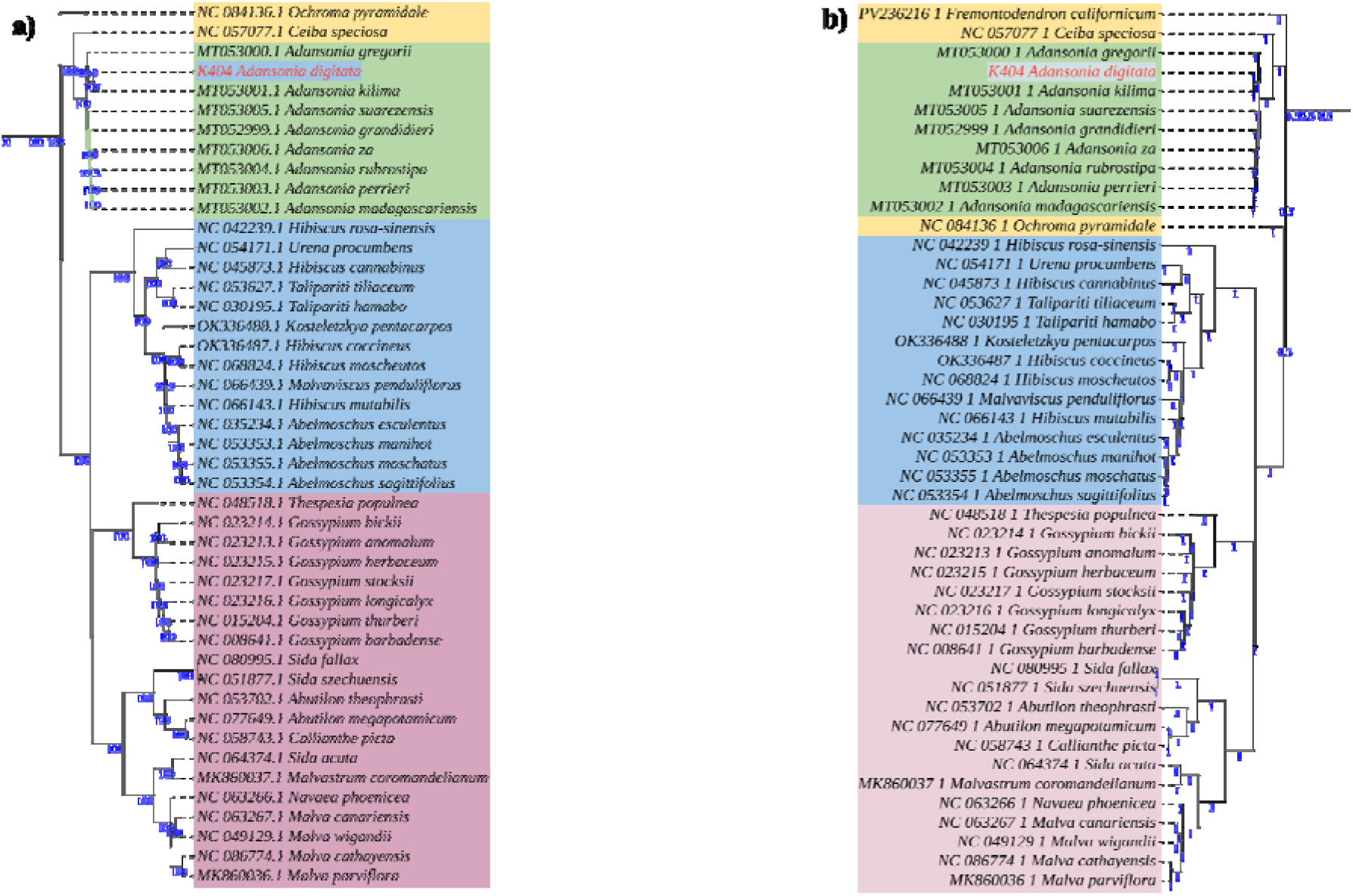
Phylogenomic placement of A. digitata within the family Malvaceae based on the assembled chloroplast genome. (a) Maximum likelihood (ML) subtree showing the phylogenetic position of the assembled *A. digitata* chloroplast genome in the selected clades within the family. (b) Bayesian inference (BI) subtree showing the placement of A. digitata within the family. Complete, fully resolved ML and BI trees are provided in Supplementary Figure S2.

## Discussion

Woody angiosperm fruit trees are central to global food and nutritional security ^60–63^, making the characterization of their genomic resources a priority for fundamental research and biotechnological advancement^64–66^. In this study, the chloroplast genome of *A. digitata* was assembled and comprehensively characterized, generating a genome of 160,061 bp that exhibited the typical quadripartite plastome architecture (Fig. 1), with gene content and structural features consistent with those reported for most angiosperm chloroplast genomes^18,67,68^. A total of 115 genes were annotated, comprising 79 protein-coding genes, 32 tRNAs, and 4 rRNAs. Compositional features such as nucleotide proportions were consistent with known angiosperms. The genome was AT-rich and relatively conserved in structure and gene order, suggesting a broadly conserved genome evolution and organization. Although chloroplast genomes are generally stable, they can undergo lineage-specific modifications involving gene loss, pseudogenization, or the transfer of ancestral plastid genes to the nuclear genome. In this regard, two genes, infA and ndhD, were found to initiate translation with non-canonical start codons, with infA starting at ATT and ndhD at ACG. Such deviations from the canonical ATG start codon are common phenomena in plastomes and are often accommodated by the chloroplast translation machinery. In the case of ndhD, RNA editing is a well-documented post-transcriptional mechanism, observed in plants such as tobacco, that converts the ACG genomic codon into a functional AUG start codon in the mRNA transcript, thereby restoring correct translation^69^. Consistent with this mechanism, RNA editing predictions in the present study also supported this codon conversion (Supplementary Table S6). For infA, non-standard start codons such as UUG, GUG, or AUU have been reported in some lineages^70^, although ATT is rarely observed as a functional initiator. In many angiosperms, infA is either pseudogenized, transferred to the nucleus, or retains a standard ATG start codon^71^, and its non-canonical start codon in *A. digitata* likely indicates functional modification or ongoing degeneration.

Comparative analysis with other *Adansonia* species showed that *A. digitata* chloroplast genome falls within the expected range for genome size, structural organization, gene content, and nucleotide composition (Supplementary Table S1). Collinearity analysis further indicated structural conservation, with no major genome rearrangements, large duplications, or deletions detected among the compared plastomes - a pattern consistent with previous reports describing the broadly conserved architecture of angiosperm chloroplast genomes, where plastome sizes typically range from approximately 150,000 to 170,000 bp and maintain a stable quadripartite structure^21^. Structural differences, when present, tend to be minor and are most commonly associated with shifts at inverted repeat boundaries that accumulate through lineage divergence and speciation^21,22^. Average Nucleotide Identity (ANI) analysis revealed high sequence similarity between the A. digitata plastome and those of other Adansonia species, with values exceeding 99% (Fig. 2). The putative A. kilima plastome showed the highest similarity to A. digitata (ANI ≈ 99.96%), indicating a high degree of plastome sequence similarity. This level of similarity is consistent with the characteristically slow evolutionary rate of plastid genomes relative to nuclear genomes and is commonly observed among closely related plant species^72,73^. The limited sequence divergence across *Adansonia* plastomes indicates the conservative nature of chloroplast genome evolution and points to relatively recent divergence within the lineage. While *Adansonia* species are notably long-lived trees, generation time alone does not fully determine plastome evolutionary rate, which is also shaped by mutation rate, selection, and demographic history^17,67,74^. The near identity between the *A. digitata* and putative *A. kilima* plastomes supports a very close evolutionary relationship; however, this finding warrants careful interpretation. Given that chloroplast genomes evolve slowly and are uniparentally inherited^68,75,76^, they may lack sufficient phylogenetic resolution to distinguish recently diverged or cryptic species, and clarifying species boundaries will require complementary evidence from nuclear genomic data and broader population-level sampling.

In the genus *Adansonia*, the junction boundaries are largely conserved, indicating similar structural organization across the genus. A notable exception is observed in *A. grandidieri* and *A. perrieri*, where a duplication of ycf1 at the JSB (IRa/SSC) boundary is apparent (Fig. 2). This localized duplication is most parsimoniously explained by independent, lineage-specific IR/SSC boundary shifts rather than inheritance from a shared ancestral duplication event, an interpretation supported by the phylogenomic topology (Fig. 8) and consistent with findings from prior studies in which *A. grandidieri* and *A. perrieri* resolve into distinct clades^2^ - an arrangement that contradicts a shared ancestral origin for the observed boundary variation. Similar junction dynamics have been reported across diverse plant lineages, where small IR expansions and contractions arise repeatedly and independently among closely related plastomes, suggesting ongoing independent fine-scale structural evolution at repeat boundaries^21,77^.

The repeat landscape of the plastome of *A. digitata* was generally consistent with patterns reported across angiosperm chloroplast genomes^30,68,78,79^, suggesting that the repeat dynamics observed are general features of plastid genome organization rather than something unique to this species. A total of 50 long repeats, most of which were forward and palindromic in orientation and had a size of 30-36 bp were identified. The predominantly short length and low sequence divergence of these repeats point to a structurally stable genome with little evidence of pervasive repeat-driven rearrangement. Particularly noteworthy was the concentration of repeats near the inverted repeat boundaries, an aspect widely associated with the maintenance of the quadripartite plastome architecture and hypothesised to contribute to genome stabilization through recombination-based repair^80^. The SSR profile was similarly typical of angiosperm plastomes^78,79^, dominated by A/T-rich mononucleotide motifs consistent with the well-documented AT compositional bias of plastid genomes^81^. Complex repeat units, dinucleotide through pentanucleotide motifs, were rarer, mirroring trends observed across land plants. The uneven distribution of SSRs across genomic compartments was particularly of interest: intergenic spacers had the highest proportion, followed by intronic regions, while coding sequences were comparatively repeat-poor. This pattern is consistent with the functional constrain^82^, given that coding regions are under purifying selection and are therefore less tolerant of the insertions and deletions that SSRs can introduce. Among protein-coding genes, SSRs were identified in a number of functionally important loci. These included *matK, atpI, rpoC2, rpoC1, rpoB, psaA, rbcL, cemA, ycf1*, and *ndhF*. While intron-rich genes such as *clpP1, rpl16*, and *ndhA* showed comparatively higher repeat densities, likely indicating the greater structural flexibility permitted in non-coding intronic sequences. Beyond their evolutionary significance as potential mutational hotspots and drivers of local recombination, the SSRs identified in *A. digitata* represent a promising resource for developing molecular markers for population genetic and phylogeographic studies within *Adansonia*. Such tools remain relatively limited for the genus *Adansonia*.

From the structural features, the prediction of RNA editing sites identified post-transcriptional modification in the *A. digitata* plastome, with 578 editing sites identified across 63 protein-coding genes. The majority of these were C→U conversions occurring mostly at the second codon position, a pattern consistent with the well-established role of RNA editing in restoring conserved amino acid identities and maintaining protein function in chloroplast genomes^69^. The complete absence of edits at the third codon position is noteworthy and indicates functional constraint on synonymous sites. Genes with the highest editing loads, *ycf1, rpoC2, ycf2*, and *matK*, are among the largest and most functionally complex in the plastome^83–85^, so their high editing frequencies are perhaps expected. At the other extreme, 21 genes, including *psaC, rps12*, and *psbA*, showed no predicted editing sites, suggesting that their transcripts are either already correctly encoded at the DNA level or subject to different post-transcriptional regulatory mechanisms. The dominance of amino acid changes involving hydrophobic and polar residues, particularly F→L and S→L substitutions, is consistent with patterns reported across plastosomes^69^ and likely indicates selection to restore specific physicochemical properties in functionally critical proteins. The detection of rare stop codon modifications adds an intriguing dimension (Supplementary Table S6), hinting that RNA editing may occasionally influence translation termination - a finding that requires experimental validation.

The codon usage landscape of the *A. digitata* chloroplast genome detected a structured and biologically meaningful pattern shaped by the interplay between mutational pressure and gene-specific selective forces. Rather than indicating a single pervasive mechanism, results from multiple complementary analyses, RSCU, codon bias indices, ENC-GC3 neutrality plots, PR2 bias, and correspondence analysis came to a consistent conclusion: mutational pressure establishes the baseline compositional framework, while natural selection drives deviations in specific functional gene groups. RSCU analysis confirmed clear non-random synonymous codon preferences, with codons ending in A or T favored across most amino acids (Fig. 4) - an expected consequence of the overall AT-rich composition of *A. digitata* plastomes (Fig. 1). Highly preferred codons such as TTA (Leu), AGA (Arg), and GCT (Ala) showed high RSCU values, while those with low RSCU values likely indicate compositional constraints or reduced availability of corresponding tRNAs. The co-existence of highly preferred, neutral, and underrepresented codons suggests that codon choice is influenced by genome-wide nucleotide bias and functional translational considerations. Overall, codon bias was low to moderate but heterogeneous across genes. The effective number of codons (Nc) ranged from 36.33 to 56.65 with a mean of approximately 47.8, supporting moderate but variable bias across the proteome (Supplementary Table S5). GC3s values were confined to a narrow range (0.18-0.32), supporting the AT bias at synonymous sites. In the Nc-GC3s neutrality plot (Fig. 5a), most genes clustered near the expected curve, and this pattern was statistically supported by an insignificant Kolmogorov-Smirnov test, indicating that mutational pressure is the dominant genome-wide driver of codon usage. At the gene-level, deviations were clearly evident. A number of genes, psbA, rps14, rps8, rps18, atpF, cemA, psbD, petD, and rpl16, had values below the neutrality curve and showed low Nc values and positive eNC deviation ratios - aspects that only stronger codon bias than composition alone can explain. Many of these are highly expressed genes involved in photosynthesis or core chloroplast functions (Supplementary Table S2), supporting the interpretation that translational selection contributes to codon optimization in these loci. In contrast, genes such as clpP1, pafI, and ndhJ showed high Nc values and negative deviation ratios, pointing to weaker codon preference and possible relaxation of translational selection.

Neutrality analysis of GC12 versus GC3 provided additional resolution on this balance. The regression slope of 0.28 indicates that mutational pressure accounts for approximately 28% of codon usage variation, with the remaining ∼72% attributable to natural selection and other constraints. The consistently higher and more stable GC12 values relative to GC3 indicate functional constraint at first and second codon positions, which directly determine amino acid identity, compared with the greater flexibility at the degenerate third position^86^. PR2 bias analysis supported this analogy, with most genes deviating from the central (0.5, 0.5) point and showing a general preference for T over A and C over G at the third codon position - a phenomenon that cannot be attributed to mutation alone and instead implicates additional selective or repair-related biases. Correspondence analysis also supported the conclusion that codon usage in *A. digitata* is non-random and biologically structured, with most explained variation concentrated in the first two axes and primary separation driven by nucleotide preference at the third codon position. These analytical frameworks indicate a plastome governed primarily by AT-rich mutational bias, with superimposed but detectable gene-specific translational selection - a dual influence that indicates the evolutionary stability typical of plastid genomes^87^, combined with localized codon optimization in functionally important and highly expressed loci.

In broader sequence variation, nucleotide diversity analysis across the plastome identified a biologically coherent structure that supports the well-established functional architecture of plastosomes^67^. Protein-coding genes were conserved, exhibiting low diversity consistent with the action of purifying selection on genes essential for photosynthesis, transcription, and core chloroplast metabolism^88^. A small number of loci emerged as relative diversity hotspots, that is, *infA, rps16, rpl33, ycf1*, and *ndhI*, consistent with previous studies identifying these genes as carriers of useful phylogenetic and population-level signals^89^. As expected, sequence variation was concentrated mostly in the intergenic regions of the LSC, while the IR regions remained highly conserved (Fig. 7c, Fig. 7b), most likely due to copy-correction mechanisms and the structural constraints inherent to inverted repeats. The dominance of low dN/dS (ω) ratios across the genome provided evidence for widespread purifying selection, particularly in genes encoding photosystem components, ribosomal proteins, and NADH dehydrogenase subunits (Fig. 7a). A few genes, *matK, ycf1, accD*, and *rpoB*, showed comparatively high ω values, suggesting either relaxed functional constraint or limited adaptive divergence in these loci. The heterogeneity in transition-transversion bias across genes also points to locus-specific mutational dynamics. These results indicate a plastome shaped primarily by functional constraint, with a limited subset of faster-evolving regions that are particularly informative for evolutionary and comparative plastosome analyses.

The evolutionary relics in the assembled *A. digitata* plastosomes showed a clear expression in the phylogenetic analyses, which were based on 248 Malvaceae chloroplast genomes and produced fully congruent Maximum Likelihood and Bayesian topologies. In the two analytical frameworks, the assembled plastosome clustered within the *Adansonia* species, forming a monophyletic clade with *A. kilima* and *A. gregorii* (Fig. 9). This arrangement was recovered with maximal statistical support (BS = 100; PP = 1.0). The clear separation of *Adansonia* from related genera such as *Ceiba*, *Ochroma*, and *Firmiana*, additionally confirms plastome divergence across *Malvaceae* lineages and supports the taxonomic coherence of the genus. The positioning of the putative *A. kilima* with *A. digitata* is particularly noteworthy. Initially described on the basis of morphological distinctions, *A. kilima* showed insignificant genetic differentiation from *A. digitata* at the plastome level, consistent with observations from prior molecular studies^6^. This raises caution on the resolving power of chloroplast data for species delimitation: while plastome sequences provide good phylogenetic signal at the genus level, their slow evolutionary rate and uniparental inheritance may render them insufficient for distinguishing recently diverged or morphologically defined species^68^. The taxonomic status of *A. kilima* therefore remains uncertain, and a definitive resolution will likely require the integration of nuclear genomic data and broader population-level sampling to determine whether the observed similarity reflects true taxonomic overlap, very recent divergence, or a history of introgression between the two lineages.

## Conclusion

This study presents a comprehensive characterization of the *A. digitata* plastosome, providing insights into its structure, gene content, codon usage patterns, repeat composition, and evolutionary relationships within Malvaceae. The plastome exhibits a canonical quadripartite architecture, moderate codon usage bias shaped primarily by mutational pressure, and a conserved arrangement of coding, non-coding regions, and SSR and long repeats. RNA editing analyses predicted extensive post-transcriptional modifications, highlighting the functional mechanisms that maintain protein integrity and plastome stability. Phylogenomic analyses positioned *A. digitata* within a well-supported Adansonia clade, closely related to *A. gregorii* and the putative *A. kilima*, although the latter’s status as a distinct species remains unresolved due to the highly conserved nature of chloroplast genomes. These findings support the evolutionary stability of the *A. digitata* plastome while revealing subtle signals of functional optimization in highly expressed genes, particularly those associated with photosynthesis. The SSRs and repeat elements identified in this study provide a valuable resource for future population genetic, phylogeographic, and conservation studies. Overally, this work establishes a genomic foundation for *A. digitata*, facilitating continued research into its evolutionary history, taxonomy, and potential adaptive mechanisms. Future studies integrating nuclear genomic data, broader population-level sampling, and functional analyses will be essential to resolving remaining taxonomic ambiguities and fully understanding the ecological and evolutionary significance of this iconic species.

## Data availability

The raw sequencing data and assembled African baobab (*Adansonia digitata)* chloroplast genome generated in this study will be deposited in the NCBI GenBank database prior to publication. The accession number will be provided upon submission.

## Acknowledgements

The authors wish to express their gratitude to all those who assisted during the project period.

## Contributions

O.F.O.: Data curation, visualization, and writing (original draft). Z.M.: Visualization and writing (original draft). E.O.O.: Writing (original draft and editing). M.W.: Writing (review and editing). O.M.: Writing (review and editing). R.K.: Funding acquisition, project administration, supervision, and writing (review and editing).

## Reference

1. Baum, D. A., Small, R. L. & Wendel, J. F. Biogeography and Floral Evolution of Baobabs Adansonia, Bombacaceae as Inferred From Multiple Data Sets. Syst. Biol. 47, 181–207 (1998).

2. Wan, J.-N. et al. The rise of baobab trees in Madagascar. Nature 629, 1091–1099 (2024).

3. Baum, D. A. A Systematic Revision of Adansonia (Bombacaceae). Ann. Mo. Bot. Gard. 82, 440–471 (1995).

4. Cron, G. V. et al. One African baobab species or two? Synonymy of Adansonia kilima and A. digitata. TAXON 65, 1037–1049 (2016).

5. Douie, C., Whitaker, J. & Grundy, I. Verifying the presence of the newly discovered African baobab, *Adansonia kilima*, in Zimbabwe through morphological analysis. South Afr. J. Bot. 100, 164–168 (2015).

6. Pettigrew FRS, J. D. et al. Morphology, ploidy and molecular phylogenetics reveal a new diploid species from Africa in the baobab genus Adansonia (Malvaceae: Bombacoideae). TAXON 61, 1240–1250 (2012).

7. Lisao, K., Geldenhuys, C. J. & Chirwa, P. W. Traditional uses and local perspectives on baobab (*Adansonia digitata*) population structure by selected ethnic groups in northern Namibia. South Afr. J. Bot. 113, 449–456 (2017).

8. Meinhold, K. & Darr, D. Using a multi-stakeholder approach to increase value for traditional agroforestry systems: the case of baobab (Adansonia digitata L.) in Kilifi, Kenya. Agrofor. Syst. 95, (2021).

9. Assogbadjo, A. E. et al. Folk Classification, Perception, and Preferences of Baobab Products in West Africa: Consequences for Species Conservation and Improvement. Econ. Bot. 62, 74–84 (2008).

10. Chapotin, S. M., Razanameharizaka, J. H. & Holbrook, N. M. Water relations of baobab trees (Adansonia spp. L.) during the rainy season: does stem water buffer daily water deficits? Plant Cell Environ. 29, 1021–1032 (2006).

11. Meinhold, K. & Darr, D. Keeping Up With Rising (Quality) Demands? The Transition of a Wild Food Resource to Mass Market, Using the Example of Baobab in Malawi. Front. Sustain. Food Syst. 6, (2022).

12. Hyacinthe, T. et al. Variability of vitamins B1, B2 and minerals content in baobab (Adansonia digitata) leaves in East and West Africa. Food Sci. Nutr. 3, 17–24 (2015).

13. Olejniczak, S. A., Łojewska, E., Kowalczyk, T. & Sakowicz, T. Chloroplasts: state of research and practical applications of plastome sequencing. Planta 244, 517–527 (2016).

14. Mathuria, A. et al. Chloroplast Genomics and Their Uses in Crop Improvement. in Advances in Genomics: Methods and Applications 331–356 (Springer, 2024).

15. Venter, S. M. & Witkowski, E. T. Baobabs as symbols of resilience. Nat. Plants 10, 732–735 (2024).

16. Stiller, J. W. Plastid endosymbiosis, genome evolution and the origin of green plants. Trends Plant Sci. 12, 391–396 (2007).

17. Stadnichuk, I. & Kusnetsov, V. Endosymbiotic origin of chloroplasts in plant cells’ evolution. Russ. J. Plant Physiol. 68, 1–16 (2021).

18. Dobrogojski, J., Adamiec, M. & Luciński, R. The chloroplast genome: a review. Acta Physiol. Plant. 42, 98 (2020).

19. Schneider, A. Organelle inheritance: understanding the basis of plastid transmission for transgenic engineering. J. Mitochondria Plast. Endosymbiosis 1, 2261790 (2023).

20. Gitzendanner, M. A., Soltis, P. S., Yi, T.-S., Li, D.-Z. & Soltis, D. E. Plastome phylogenetics: 30 years of inferences into plant evolution. in Advances in botanical research vol. 85 293–313 (Elsevier, 2018).

21. Cauz-Santos, L. A. Beyond conservation: the landscape of chloroplast genome rearrangements in angiosperms. New Phytol. (2025).

22. Magadum, S., Banerjee, U., Murugan, P., Gangapur, D. & Ravikesavan, R. Gene duplication as a major force in evolution. J. Genet. 92, 155–161 (2013).

23. Rensing, S. A. Gene duplication as a driver of plant morphogenetic evolution. Curr. Opin. Plant Biol. 17, 43–48 (2014).

24. Zhou, L., Chen, T., Qiu, X., Liu, J. & Guo, S. Evolutionary differences in gene loss and pseudogenization. Struct. Var. Chloroplast Genome Relat. Bioinforma. Tools (2024).

25. Xu, Z., Jiang, Y. & Zhou, G. Response and adaptation of photosynthesis, respiration, and antioxidant systems to elevated CO2 with environmental stress in plants. Front. Plant Sci. 6, 701 (2015).

26. Yamori, W. Photosynthetic response to fluctuating environments and photoprotective strategies under abiotic stress. J. Plant Res. 129, 379–395 (2016).

27. Nguyen, V. B. et al. Authentication markers for five major Panax species developed via comparative analysis of complete chloroplast genome sequences. J. Agric. Food Chem. 65, 6298–6306 (2017).

28. Song, Y. et al. The population genomic analyses of chloroplast genomes shed new insights on the complicated ploidy and evolutionary history in Fragaria. Front. Plant Sci. 13, 1065218 (2023).

29. Li, C. et al. Chloroplast genome, an effective strategy for identifying hybrid species, using Dendrobium’Black Gold’as an example. BMC Plant Biol. 25, 1257 (2025).

30. Daniell, H., Lin, C.-S., Yu, M. & Chang, W.-J. Chloroplast genomes: diversity, evolution, and applications in genetic engineering. Genome Biol. 17, 134 (2016).

31. Bock, R. Engineering Plastid Genomes: Methods, Tools, and Applications in Basic Research and Biotechnology. Annu. Rev. Plant Biol. 66, 211–241 (2015).

32. Kehlenbeck, K., Padulosi, S. & Alercia, A. Descriptors for baobab (Adansonia digitata L.). Bioversity Int. Rome Italy World Agrofor. Cent. Nairobi Kenya ISBN*-*13 978–92 (2015).

33. Chen, S., Zhou, Y., Chen, Y. & Gu, J. fastp: an ultra-fast all-in-one FASTQ preprocessor. Bioinformatics 34, i884–i890 (2018).

34. Jin, J.-J. et al. GetOrganelle: a fast and versatile toolkit for accurate de novo assembly of organelle genomes. Genome Biol. 21, 241 (2020).

35. Bankevich, A. et al. SPAdes: a new genome assembly algorithm and its applications to single-cell sequencing. J. Comput. Biol. 19, 455–477 (2012).

36. Zou, T., Li, D., Zhao, C.-Y. & Chen, M.-L. Chloroplast whole genome assembly and phylogenetic analysis of Persicaria criopolitana reveals its new taxonomic status. Sci. Rep. 15, 19890 (2025).

37. Wick, R. R., Schultz, M. B., Zobel, J. & Holt, K. E. Bandage: interactive visualization of de novo genome assemblies. Bioinformatics 31, 3350–3352 (2015).

38. Qu, X.-J., Moore, M. J., Li, D.-Z. & Yi, T.-S. PGA: a software package for rapid, accurate, and flexible batch annotation of plastomes. Plant Methods 15, 50 (2019).

39. Zhang, N.-N., et al. PlastidHub: an integrated analysis platform for plastid phylogenomics and comparative genomics. Plant Divers. (2025).

40. Tillich, M. et al. GeSeq–versatile and accurate annotation of organelle genomes. Nucleic Acids Res. 45, W6–W11 (2017).

41. Chan, P. P., Lin, B. Y., Mak, A. J. & Lowe, T. M. tRNAscan-SE 2.0: improved detection and functional classification of transfer RNA genes. Nucleic Acids Res. 49, 9077–9096 (2021).

42. Laslett, D. & Canback, B. ARAGORN, a program to detect tRNA genes and tmRNA genes in nucleotide sequences. Nucleic Acids Res. 32, 11–16 (2004).

43. Lehwark, P. & Greiner, S. GB2sequin-A file converter preparing custom GenBank files for database submission. Genomics 111, 759–761 (2019).

44. Sayers, E. W. et al. GenBank 2024 Update. Nucleic Acids Res. 52, D134–D137 (2024).

45. Greiner, S., Lehwark, P. & Bock, R. OrganellarGenomeDRAW (OGDRAW) version 1.3. 1: expanded toolkit for the graphical visualization of organellar genomes. Nucleic Acids Res. 47, W59–W64 (2019).

46. Huang, L., Yu, H., Wang, Z. & Xu, W. CPStools: A package for analyzing chloroplast genome sequences. iMetaOmics 1, e25 (2024).

47. Kurtz, S. et al. REPuter: the manifold applications of repeat analysis on a genomic scale. Nucleic Acids Res. 29, 4633–4642 (2001).

48. Peden, J. F. Analysis of codon usage. PhD thesis. U. K. Univ. Nottm. (1999).

49. Lenz, H. et al. Introducing the plant RNA editing prediction and analysis computer tool PREPACT and an update on RNA editing site nomenclature. Curr. Genet. 56, 189–201 (2010).

50. Chen, Y., Ye, W., Zhang, Y. & Xu, Y. High speed BLASTN: an accelerated MegaBLAST search tool. Nucleic Acids Res. 43, 7762–7768 (2015).

51. Amiryousefi, A., Hyvönen, J. & Poczai, P. IRscope: an online program to visualize the junction sites of chloroplast genomes. Bioinformatics 34, 3030–3031 (2018).

52. Katoh, K., Misawa, K., Kuma, K. & Miyata, T. MAFFT: a novel method for rapid multiple sequence alignment based on fast Fourier transform. Nucleic Acids Res. 30, 3059–3066 (2002).

53. Ranwez, V., Douzery, E. J., Cambon, C., Chantret, N. & Delsuc, F. MACSE v2: toolkit for the alignment of coding sequences accounting for frameshifts and stop codons. Mol. Biol. Evol. 35, 2582–2584 (2018).

54. Gao, F. et al. EasyCodeML: A visual tool for analysis of selection using CodeML *Ecol*. Evol. 9, 3891–3898 (2019).

55. Team, R. C. R language definition. Vienna Austria R Found. Stat. Comput. 3, 116 (2000).

56. Wickham, H. et al. Welcome to the Tidyverse. J. Open Source Softw. 4, 1686 (2019).

57. Minh, B. Q. et al. IQ-TREE 2: new models and efficient methods for phylogenetic inference in the genomic era. Mol. Biol. Evol. 37, 1530–1534 (2020).

58. Ronquist, F. et al. MrBayes 3.2: efficient Bayesian phylogenetic inference and model choice across a large model space. Syst. Biol. 61, 539–542 (2012).

59. Letunic, I. & Bork, P. Interactive Tree of Life (iTOL) v6: recent updates to the phylogenetic tree display and annotation tool. Nucleic Acids Res. 52, W78–W82 (2024).

60. Bundela, A. K., Abhilash, P. C. & Peñuelas, J. Securing Wild Edible Plants for Planetary Healthy Diet. Anthr. Sci. 1–3 (2023) doi:10.1007/s44177-023-00054-4.

61. Tadesse, D., Masresha, G., Lulekal, E. & Wondafrash, M. A systematic review exploring the diversity and food security potential of wild edible plants in Ethiopia. Sci. Rep. 14, 17821 (2024).

62. Kumar, B. M., Bhavya, G., De Britto, S. & Jogaiah, S. Wild edible plants for food security, dietary diversity, and nutraceuticals: a global overview of emerging research. Front. Sustain. Food Syst. 9, (2025).

63. Vázquez-Martin, Á. E. & Aguilar-Rivera, N. Edible Flora as a Sustainable Resource for World Food. in Handbook of Climate Change Across the Food Supply Chain (eds Leal Filho, W., Djekic, I., Smetana, S. & Kovaleva, M.) 145–161 (Springer International Publishing, Cham, 2022). doi:10.1007/978-3-030-87934-1_8.

64. Shi, J., Tian, Z., Lai, J. & Huang, X. Plant pan-genomics and its applications. Mol. Plant 16, 168–186 (2023).

65. Kumar, R. et al. Advances in genomic tools for plant breeding: harnessing DNA molecular markers, genomic selection, and genome editing. Biol. Res. 57, 80 (2024).

66. Wambugu, P. W., Ndjiondjop, M.-N. & Henry, R. J. Role of genomics in promoting the utilization of plant genetic resources in genebanks. Brief. Funct. Genomics 17, 198–206 (2018).

67. Jansen, R. K. & Ruhlman, T. A. Plastid Genomes of Seed Plants. in Genomics of Chloroplasts and Mitochondria (eds Bock, R. & Knoop, V.) 103–126 (Springer Netherlands, Dordrecht, 2012). doi:10.1007/978-94-007-2920-9_5.

68. Wicke, S., Schneeweiss, G. M., dePamphilis, C. W., Müller, K. F. & Quandt, D. The evolution of the plastid chromosome in land plants: gene content, gene order, gene function. Plant Mol. Biol. 76, 273–297 (2011).

69. Hirose, T. & Sugiura, M. Both RNA editing and RNA cleavage are required for translation of tobacco chloroplast ndhD mRNA: a possible regulatory mechanism for the expression of a chloroplast operon consisting of functionally unrelated genes. EMBO J. 16, 6804–6811 (1997).

70. Hirose, T., Ideue, T., Wakasugi, T. & Sugiura, M. The chloroplast infA gene with a functional UUG initiation codon. FEBS Lett. 445, 169–172 (1999).

71. Millen, R. S. et al. Many parallel losses of infA from chloroplast DNA during angiosperm evolution with multiple independent transfers to the nucleus. Plant Cell 13, 645–658 (2001).

72. Smith, D. R. Mutation rates in plastid genomes: they are lower than you might think. Genome Biol. Evol. 7, 1227–1234 (2015).

73. Zhong, Y. et al. Comparative genomics and phylogenetic analysis of six Malvaceae species based on chloroplast genomes. BMC Plant Biol. 24, 1245 (2024).

74. Guo, Y.-Y., Yang, J.-X., Bai, M.-Z., Zhang, G.-Q. & Liu, Z.-J. The chloroplast genome evolution of Venus slipper (Paphiopedilum): IR expansion, SSC contraction, and highly rearranged SSC regions. BMC Plant Biol. 21, 248 (2021).

75. Jr, C. W. B. The Inheritance of Genes in Mitochondria and Chloroplasts: Laws, Mechanisms, and Models. Annu. Rev. Genet. 35, 125–148 (2001).

76. Wolfe, K. H., Li, W.-H. & Sharp, P. M. Rates of nucleotide substitution vary greatly among plant mitochondrial, chloroplast, and nuclear DNAs. Proc. Natl. Acad. Sci. 84, 9054–9058 (1987).

77. Wang, R.-J. et al. Dynamics and evolution of the inverted repeat-large single copy junctions in the chloroplast genomes of monocots. BMC Evol. Biol. 8, 36 (2008).

78. Provan, J., Powell, W. & Hollingsworth, P. M. Chloroplast microsatellites: new tools for studies in plant ecology and evolution. Trends Ecol. Evol. 16, 142–147 (2001).

79. Ebert, D. & Peakall, R. Chloroplast simple sequence repeats (cpSSRs): technical resources and recommendations for expanding cpSSR discovery and applications to a wide array of plant species. Mol. Ecol. Resour. 9, 673–690 (2009).

80. Ruhlman, T. A., Zhang, J., Blazier, J. C., Sabir, J. S. & Jansen, R. K. Recombination-dependent replication and gene conversion homogenize repeat sequences and diversify plastid genome structure. Am. J. Bot. 104, 559–572 (2017).

81. Long, H. et al. Evolutionary determinants of genome-wide nucleotide composition. *Nat*. Ecol. Evol. 2, 237–240 (2018).

82. Zhao, M., Shu, G., Hu, Y., Cao, G. & Wang, Y. Pattern and variation in simple sequence repeat (SSR) at different genomic regions and its implications to maize evolution and breeding. BMC Genomics 24, 136 (2023).

83. Tillich, M., Lehwark, P., Morton, B. R. & Maier, U. G. The evolution of chloroplast RNA editing. Mol. Biol. Evol. 23, 1912–1921 (2006).

84. Chateigner-Boutin, A.-L. & Small, I. Plant RNA editing. RNA Biol. 7, 213–219 (2010).

85. Drescher, A., Ruf, S., Calsa Jr, T., Carrer, H. & Bock, R. The two largest chloroplast genome-encoded open reading frames of higher plants are essential genes. Plant J. 22, 97–104 (2000).

86. Mondal, S. K., Kundu, S., Das, R. & Roy, S. Analysis of phylogeny and codon usage bias and relationship of GC content, amino acid composition with expression of the structural nif genes. J. Biomol. Struct. Dyn. 34, 1649–1666 (2016).

87. Suzuki, H. & Morton, B. R. Codon adaptation of plastid genes. PLoS One 11, e0154306 (2016).

88. Bauer, J., Hiltbrunner, A. & Kessler, F. Molecular biology of chloroplast biogenesis: gene expression, protein import and intraorganellar sorting. Cell. Mol. Life Sci. CMLS 58, 420–433 (2001).

89. Shaw, J., Lickey, E. B., Schilling, E. E. & Small, R. L. Comparison of whole chloroplast genome sequences to choose noncoding regions for phylogenetic studies in angiosperms: the tortoise and the hare III. Am. J. Bot. 94, 275–288 (2007).

