## Supplementary Table S1 for "Complete chloroplast genome of African Baobab (*Adansonia digitata* L.): structural characterization, comparative genomics, and phylogenetic placement within Malvaceae"

**Table S1**: Genome attributes across the *Adansonia* genus, comparing nucleotide composition, and genic features across the plastomes.

| **Genome attributes** | ***A. grandidieri*** | ***A. madagascariensis*** | ***A. perrieri*** | ***A. rubrostipa*** | ***A. gregorii*** | ***A. suarezensis*** | ***A. za*** | ***A. kilima*** | ***A. digitata*** |
| --- | --- | --- | --- | --- | --- | --- | --- | --- | --- |
| Total genome length (bp) | 160094 | 160230 | 159710 | 160322 | 159891 | 160621 | 160449 | 160072 | 160061 |
| LSC (bp) | 88983 | 89153 | 89024 | 89263 | 88836 | 89515 | 89329 | 88972 | 88961 |
| SSC (bp) | 20003 | 20013 | 19626 | 20003 | 19983 | 20026 | 20024 | 19994 | 19994 |
| IR (bp) | 51108 | 51064 | 51060 | 51056 | 51072 | 51080 | 51096 | 51106 | 51106 |
| Protein-coding region | 85116 | 85028 | 84792 | 84792 | 84994 | 84789 | 84792 | 84624 | 84624 |
| rRNA coding region (bp) | 14272 | 14272 | 14272 | 14272 | 14272 | 14272 | 14272 | 14272 | 14272 |
| tRNA coding region (bp) | 5115 | 5115 | 5115 | 5115 | 5115 | 5115 | 5115 | 5115 | 5115 |
| Intron (bp) | 30985 | 30937 | 30948 | 30959 | 29442 | 31001 | 30964 | 31012 | 31012 |
| Total GC (%) | 36.86 | 36.84 | 36.9 | 36.83 | 36.88 | 36.78 | 36.79 | 36.88 | 36.88 |
| LSC GC (%) | 34.66 | 34.6 | 34.66 | 34.59 | 34.68 | 34.52 | 34.54 | 34.68 | 34.68 |
| SSC GC(%) | 31.2 | 31.17 | 31.31 | 31.19 | 31.18 | 31.12 | 31.17 | 31.21 | 31.17 |
| IR GC(%) | 42.92 | 42.96 | 42.97 | 42.95 | 42.95 | 42.95 | 42.92 | 42.92 | 42.92 |
| Protein coding regions GC(%) | 38.22 | 38.24 | 38.26 | 38.27 | 38.22 | 38.24 | 38.25 | 38.24 | 38.25 |
| Intron GC(%) | 39.99 | 40.03 | 40.03 | 40.02 | 40.33 | 39.98 | 39.99 | 39.98 | 39.98 |
| rRNA GC(%) | 53.77 | 53.77 | 53.77 | 53.77 | 53.77 | 53.77 | 53.77 | 53.77 | 53.77 |
| tRNA GC(%) | 53.58 | 53.58 | 53.58 | 53.58 | 53.58 | 53.58 | 53.58 | 53.58 | 53.58 |
| Number of protein-coding genes | 79 (97) | 79 (97) | 79 (96) | 79 (96) | 79 (97) | 79 (96) | 79 (96) | 79 (95) | 79 (95) |
| Number of tRNA coding genes | 32 (66) | 32 (66) | 32 (66) | 32 (66) | 32 (66) | 32 (66) | 32 (66) | 32 (66) | 32 (66) |
| Number of rRNA coding genes | 4 (10) | 4 (10) | 4 (10) | 4 (10) | 4 (10) | 4 (10) | 4 (10) | 4 (10) | 4 (10) |
| Number of introns | 43 | 43 | 43 | 43 | 41 | 43 | 43 | 43 | 43 |
| Number of genes with introns | 22 | 22 | 22 | 22 | 22 | 22 | 22 | 22 | 22 |
