## Supplementary Table S2 for "Complete chloroplast genome of African Baobab (*Adansonia digitata* L.): structural characterization, comparative genomics, and phylogenetic placement within Malvaceae"

**Table S2**. Annotated genes of the *A. digitata* chloroplast genome, classified by function, gene category, and structural features.

| **Gene Function** | **Gene category** | **Genes** | **Gene features** | **Group of genes** |
| --- | --- | --- | --- | --- |
| Splicing of group II introns | Metabolism / RNA processing | matK | CDS, gene | Maturase |
| Assembly and biogenesis of c-type cytochromes | Metabolism / RNA processing | ccsA | CDS, gene | C-type cytochrome synthase gene |
| Envelope membrane protein | Metabolism / RNA processing | cemA | CDS, gene | Envelope membrane protein |
| ATP-dependent serine protease | Metabolism / RNA processing | clpP1** | CDS, exon, gene, intron | Protease |
| NADH dehydrogenase | Electron transport | ndhA* | CDS, exon, gene, intron | Subunit of NADH dehydrogenase |
| NADH dehydrogenase | Electron transport | ndhB* | CDS, exon, gene, intron | Subunit of NADH dehydrogenase |
| NADH dehydrogenase | Electron transport | ndhC | CDS, gene | Subunit of NADH dehydrogenase |
| NADH dehydrogenase | Electron transport | ndhD | CDS, gene | Subunit of NADH dehydrogenase |
| NADH dehydrogenase | Electron transport | ndhE | CDS, gene | Subunit of NADH dehydrogenase |
| NADH dehydrogenase | Electron transport | ndhF | CDS, gene | Subunit of NADH dehydrogenase |
| NADH dehydrogenase | Electron transport | ndhG | CDS, gene | Subunit of NADH dehydrogenase |
| NADH dehydrogenase | Electron transport | ndhH | CDS, gene | Subunit of NADH dehydrogenase |
| NADH dehydrogenase | Electron transport | ndhI | CDS, gene | Subunit of NADH dehydrogenase |
| NADH dehydrogenase | Electron transport | ndhJ | CDS, gene | Subunit of NADH dehydrogenase |
| NADH dehydrogenase | Electron transport | ndhK | CDS, gene | Subunit of NADH dehydrogenase |
| Translational machinery | Translation | infA | CDS, gene | Translation initiation factor |
| Photosystem I | Photosynthesis / Light reactions | pafI* | CDS, exon, gene, intron | Photosystem I assembly factor I |
| Photosystem I | Photosynthesis / Light reactions | pafII** | CDS, gene | Photosystem I assembly factor II |
| Photosystem I | Photosynthesis / Light reactions | pbf1 | CDS, gene | Photosystem biogenesis factor 1 |
| Fatty acid biosynthesis | Lipid metabolism | accD | CDS, gene | Acetyl-CoA carboxylase subunit beta |
| ATP synthase | ATP synthase | atpA | CDS, gene | CF1 subunit alpha |
| ATP synthase | ATP synthase | atpB | CDS, gene | CF1 subunit beta |
| ATP synthase | ATP synthase | atpE | CDS, gene | CF1 subunit epsilon |
| ATP synthase | ATP synthase | atpF* | CDS, exon, gene, intron | CF0 subunit I |
| ATP synthase | ATP synthase | atpH | CDS, gene | CF0 subunit III |
| ATP synthase | ATP synthase | atpI | CDS, gene | CF0 subunit IV |
| Cytochrome b6/f complex | Electron transport | petA | CDS, gene | Cytochrome f |
| Cytochrome b6/f complex | Electron transport | petB* | CDS, exon, gene, intron | Cytochrome b6 |
| Cytochrome b6/f complex | Electron transport | petD* | CDS, exon, gene, intron | Cytochrome b6/f subunit IV |
| Cytochrome b6/f complex | Electron transport | petG | CDS, gene | Cytochrome b6/f subunit G |
| Cytochrome b6/f complex | Electron transport | petL | CDS, gene | Cytochrome b6/f subunit L |
| Cytochrome b6/f complex | Electron transport | petN | CDS, gene | Cytochrome b6/f subunit N |
| Photosystem I | Photosynthesis / Light reactions | psaA | CDS, gene | Photosystem I P700 apoprotein A1 |
| Photosystem I | Photosynthesis / Light reactions | psaB | CDS, gene | Photosystem I P700 apoprotein A2 |
| Photosystem I | Photosynthesis / Light reactions | psaC | CDS, gene | Photosystem I subunit C |
| Photosystem I | Photosynthesis / Light reactions | psaI | CDS, gene | Photosystem I subunit I |
| Photosystem I | Photosynthesis / Light reactions | psaJ | CDS, gene | Photosystem I subunit J |
| Photosystem II | Photosynthesis / Light reactions | psbA | CDS, gene | Photosystem II protein D1 |
| Photosystem II | Photosynthesis / Light reactions | psbB | CDS, gene | Photosystem II 47 kDa protein |
| Photosystem II | Photosynthesis / Light reactions | psbC | CDS, gene | Photosystem II 43 kDa protein |
| Photosystem II | Photosynthesis / Light reactions | psbD | CDS, gene | Photosystem II protein D2 |
| Photosystem II | Photosynthesis / Light reactions | psbE | CDS, gene | Cytochrome b559 subunit alpha |
| Photosystem II | Photosynthesis / Light reactions | psbF | CDS, gene | Cytochrome b559 subunit beta |
| Photosystem II | Photosynthesis / Light reactions | psbH | CDS, gene | Photosystem II subunit H |
| Photosystem II | Photosynthesis / Light reactions | psbI | CDS, gene | Photosystem II subunit I |
| Photosystem II | Photosynthesis / Light reactions | psbJ | CDS, gene | Photosystem II subunit J |
| Photosystem II | Photosynthesis / Light reactions | psbK | CDS, gene | Photosystem II subunit K |
| Photosystem II | Photosynthesis / Light reactions | psbL | CDS, gene | Photosystem II subunit L |
| Photosystem II | Photosynthesis / Light reactions | psbM | CDS, gene | Photosystem II subunit M |
| Photosystem II | Photosynthesis / Light reactions | psbT | CDS, gene | Photosystem II subunit T |
| Photosystem II | Photosynthesis / Light reactions | psbZ | CDS, gene | Photosystem II subunit Z |
| Carbon fixation | Rubisco | rbcL | CDS, gene | RuBisCO large subunit |
| Ribosomal | Translational machinery | rps11 | CDS, gene | Ribosomal protein S11 |
| Ribosomal | Translational machinery | rps12** | CDS, exon, gene, intron | Ribosomal protein S12 |
| Ribosomal | Translational machinery | rps14 | CDS, gene | Ribosomal protein S14 |
| Ribosomal | Translational machinery | rps15 | CDS, gene | Ribosomal protein S15 |
| Ribosomal | Translational machinery | rps16* | CDS, exon, gene, intron | Ribosomal protein S16 |
| Ribosomal | Translational machinery | rps18 | CDS, gene | Ribosomal protein S18 |
| Ribosomal | Translational machinery | rps19 | CDS, gene | Ribosomal protein S19 |
| Ribosomal | Translational machinery | rps2 | CDS, gene | Ribosomal protein S2 |
| Ribosomal | Translational machinery | rps3 | CDS, gene | Ribosomal protein S3 |
| Ribosomal | Translational machinery | rps4 | CDS, gene | Ribosomal protein S4 |
| Ribosomal | Translational machinery | rps7 | CDS, gene | Ribosomal protein S7 |
| Ribosomal | Translational machinery | rps8 | CDS, gene | Ribosomal protein S8 |
| Ribosomal | Translational machinery | rrn16 | gene, rRNA | Ribosomal RNA |
| Ribosomal | Translational machinery | rrn23* | exon, gene, intron, rRNA | Ribosomal RNA |
| Ribosomal | Translational machinery | rrn4.5 | gene, rRNA | Ribosomal RNA |
| Ribosomal | Translational machinery | rrn5 | gene, rRNA | Ribosomal RNA |
| Ribosomal | Translational machinery | rpl14 | CDS, gene | Large subunit of ribosomal protein |
| Ribosomal | Translational machinery | rpl16* | CDS, exon, gene, intron | Large subunit of ribosomal protein |
| Ribosomal | Translational machinery | rpl2* | CDS, exon, gene, intron | Large subunit of ribosomal protein |
| Ribosomal | Translational machinery | rpl20 | CDS, gene | Large subunit of ribosomal protein |
| Ribosomal | Translational machinery | rpl22 | CDS, gene | Large subunit of ribosomal protein |
| Ribosomal | Translational machinery | rpl23 | CDS, gene | Large subunit of ribosomal protein |
| Ribosomal | Translational machinery | rpl32 | CDS, gene | Large subunit of ribosomal protein |
| Ribosomal | Translational machinery | rpl33 | CDS, gene | Large subunit of ribosomal protein |
| Ribosomal | Translational machinery | rpl36 | CDS, gene | Large subunit of ribosomal protein |
| RNA polymerase | Transcription | rpoA | CDS, gene | DNA-dependent RNA polymerase |
| RNA polymerase | Transcription | rpoB | CDS, gene | DNA-dependent RNA polymerase |
| RNA polymerase | Transcription | rpoC1* | CDS, exon, gene, intron | DNA-dependent RNA polymerase |
| RNA polymerase | Transcription | rpoC2 | CDS, gene | DNA-dependent RNA polymerase |
| tRNA | Translation | trnA-UGC* | exon, gene, intron, tRNA | tRNA-Ala |
| tRNA | Translation | trnC-ACA* | exon, intron, tRNA | tRNA-Cys |
| tRNA | Translation | trnC-GCA | gene, tRNA | tRNA-Cys |
| tRNA | Translation | trnD-GUC | gene, tRNA | tRNA-Asp |
| tRNA | Translation | trnE-UUC* | exon, gene, intron, tRNA | tRNA-Glu |
| tRNA | Translation | trnF-GAA | gene, tRNA | tRNA-Phe |
| tRNA | Translation | trnG-GCC | gene, tRNA | tRNA-Gly |
| tRNA | Translation | trnG-UCC* | exon, gene, intron, tRNA | tRNA-Gly |
| tRNA | Translation | trnH-GUG | gene, tRNA | tRNA-His |
| tRNA | Translation | trnI-GAU \| trnE-UUC | Gene | tRNA-Ile / tRNA-Glu |
| tRNA | Translation | trnI-GAU* | exon, gene, intron, tRNA | tRNA-Ile |
| tRNA | Translation | trnK-UUU* | exon, gene, intron, tRNA | tRNA-Lys |
| tRNA | Translation | trnL-CAA | gene, tRNA | tRNA-Leu |
| tRNA | Translation | trnL-UAA* | exon, gene, intron, tRNA | tRNA-Leu |
| tRNA | Translation | trnL-UAG | gene, tRNA | tRNA-Leu |
| tRNA | Translation | trnM-CAU | gene, tRNA | tRNA-Met |
| tRNA | Translation | trnN-GUU | gene, tRNA | tRNA-Asn |
| tRNA | Translation | trnP-UGG | gene, tRNA | tRNA-Pro |
| tRNA | Translation | trnQ-UUG | gene, tRNA | tRNA-Gln |
| tRNA | Translation | trnR-ACG | gene, tRNA | tRNA-Arg |
| tRNA | Translation | trnR-UCU | gene, tRNA | tRNA-Arg |
| tRNA | Translation | trnS-CGA* | exon, gene, intron, tRNA | tRNA-Ser |
| tRNA | Translation | trnS-GCU | gene, tRNA | tRNA-Ser |
| tRNA | Translation | trnS-GGA | gene, tRNA | tRNA-Ser |
| tRNA | Translation | trnS-UGA | gene, tRNA | tRNA-Ser |
| tRNA | Translation | trnT-GGU | gene, tRNA | tRNA-Thr |
| tRNA | Translation | trnT-UGU | gene, tRNA | tRNA-Thr |
| tRNA | Translation | trnV-GAC | gene, tRNA | tRNA-Val |
| tRNA | Translation | trnV-UAC \| trnC-ACA | exon, gene | tRNA-Val / tRNA-Cys |
| tRNA | Translation | trnV-UAC* | exon, intron, tRNA | tRNA-Val |
| tRNA | Translation | trnW-CCA | gene, tRNA | tRNA-Trp |
| tRNA | Translation | trnY-GUA | gene, tRNA | tRNA-Tyr |
| Protein import | Unknown / Putative function | ycf1 | CDS, gene | Conserved hypothetical chloroplast ORF |
| Protein import | Unknown / Putative function | ycf2 | CDS, gene | Conserved hypothetical chloroplast ORF |

**** Genes with one intron, ** genes with two introns***
