## Supplementary Table S3 for "Complete chloroplast genome of African Baobab (*Adansonia digitata* L.): structural characterization, comparative genomics, and phylogenetic placement within Malvaceae"

**Table S3.** Long sequence repeats identified in the *A. digitata* chloroplast genome.

| **Repeat length** | **Position 1** | **Type** | **Repeat length** | **Position 2** | **Mismatch** | **E-value** |
| --- | --- | --- | --- | --- | --- | --- |
| 25553 | 88961 | P | 25553 | 134508 | 0 | 0.00E+00 |
| 48 | 78650 | P | 48 | 78650 | 0 | 9.09E-20 |
| 49 | 40463 | F | 49 | 42687 | -2 | 2.41E-16 |
| 40 | 53719 | P | 40 | 53719 | 0 | 5.96E-15 |
| 46 | 37720 | F | 46 | 37747 | -2 | 1.36E-14 |
| 38 | 103669 | F | 38 | 125288 | 0 | 9.54E-14 |
| 38 | 125288 | P | 38 | 145315 | 0 | 9.54E-14 |
| 41 | 40471 | F | 41 | 42695 | -1 | 1.83E-13 |
| 41 | 45635 | F | 41 | 125283 | -1 | 1.83E-13 |
| 43 | 37722 | P | 43 | 37722 | -3 | 3.10E-11 |
| 43 | 99300 | R | 43 | 99310 | -3 | 3.10E-11 |
| 43 | 99300 | C | 43 | 149669 | -3 | 3.10E-11 |
| 43 | 99310 | C | 43 | 149679 | -3 | 3.10E-11 |
| 43 | 149669 | R | 43 | 149679 | -3 | 3.10E-11 |
| 33 | 86704 | F | 33 | 86733 | 0 | 9.77E-11 |
| 36 | 45640 | F | 36 | 103669 | -1 | 1.65E-10 |
| 36 | 45640 | P | 36 | 145317 | -1 | 1.65E-10 |
| 36 | 49857 | F | 36 | 49875 | -1 | 1.65E-10 |
| 37 | 49825 | R | 37 | 49825 | -2 | 2.29E-09 |
| 34 | 37847 | F | 34 | 37867 | -1 | 2.49E-09 |
| 30 | 6017 | F | 30 | 28519 | 0 | 6.25E-09 |
| 30 | 64850 | P | 30 | 86456 | 0 | 6.25E-09 |
| 35 | 49828 | R | 35 | 49828 | -2 | 3.27E-08 |
| 35 | 99318 | C | 35 | 149687 | -2 | 3.27E-08 |
| 32 | 60218 | F | 32 | 60256 | -1 | 3.75E-08 |
| 34 | 112813 | F | 34 | 112845 | -2 | 1.23E-07 |
| 34 | 112813 | P | 34 | 136143 | -2 | 1.23E-07 |
| 34 | 112845 | P | 34 | 136175 | -2 | 1.23E-07 |
| 34 | 136143 | F | 34 | 136175 | -2 | 1.23E-07 |
| 31 | 93813 | F | 31 | 93834 | -1 | 1.45E-07 |
| 31 | 93813 | P | 31 | 155157 | -1 | 1.45E-07 |
| 31 | 93834 | P | 31 | 155178 | -1 | 1.45E-07 |
| 31 | 104604 | F | 31 | 104631 | -1 | 1.45E-07 |
| 31 | 104604 | P | 31 | 144360 | -1 | 1.45E-07 |
| 31 | 104631 | P | 31 | 144387 | -1 | 1.45E-07 |
| 31 | 144360 | F | 31 | 144387 | -1 | 1.45E-07 |
| 31 | 155157 | F | 31 | 155178 | -1 | 1.45E-07 |
| 33 | 53984 | R | 33 | 53984 | -2 | 4.64E-07 |
| 30 | 8356 | P | 30 | 47495 | -1 | 5.62E-07 |
| 30 | 37886 | F | 30 | 37910 | -1 | 5.62E-07 |
| 30 | 45649 | F | 30 | 103678 | -1 | 5.62E-07 |
| 30 | 45649 | P | 30 | 145314 | -1 | 5.62E-07 |
| 32 | 74796 | P | 32 | 74796 | -2 | 1.74E-06 |
| 31 | 99296 | F | 31 | 99322 | -2 | 6.54E-06 |
| 31 | 99296 | P | 31 | 149669 | -2 | 6.54E-06 |
| 31 | 99322 | P | 31 | 149695 | -2 | 6.54E-06 |
| 31 | 149669 | F | 31 | 149695 | -2 | 6.54E-06 |
| 30 | 37816 | R | 30 | 37816 | -2 | 2.45E-05 |
| 30 | 45649 | F | 30 | 125297 | -2 | 2.45E-05 |
| 32 | 8354 | F | 32 | 36739 | -3 | 5.23E-05 |

*****Dispersed forward (F), palindromic (P), reverse (R), and complementary ©***
