## Supplementary Table S4 for "Complete chloroplast genome of African Baobab (*Adansonia digitata* L.): structural characterization, comparative genomics, and phylogenetic placement within Malvaceae"

**Table S4.** Simple sequence repeats identified in *A. digitata* plastosome

| **Type** | **Length of SSR repeat** | **Start** | **End** | **Loci** | **Loci type** |
| --- | --- | --- | --- | --- | --- |
| AAT | 4 | 62916 | 62927 | accD-psaI | IGS |
| TAGT | 3 | 62489 | 62500 | accD-psaI | IGS |
| T | 11 | 57800 | 57810 | atpB-rbcL | IGS |
| TA | 5 | 58394 | 58403 | atpB-rbcL | IGS |
| AT | 5 | 58409 | 58418 | atpB-rbcL | IGS |
| A | 12 | 13479 | 13490 | atpF_1-atpH | IGS |
| T | 11 | 13588 | 13598 | atpF_1-atpH | IGS |
| T | 13 | 81208 | 81220 | petD_1-petD_2 | Intron |
| T | 10 | 14866 | 14875 | atpH-atpI | IGS |
| A | 11 | 16255 | 16265 | atpI-rps2 | IGS |
| T | 13 | 133585 | 133597 | ycf1 | Gene |
| T | 11 | 120416 | 120426 | ccsA-ndhD | IGS |
| T | 10 | 120478 | 120487 | ccsA-ndhD | IGS |
| T | 12 | 27158 | 27169 | rpoB | Gene |
| T | 10 | 66070 | 66079 | cemA-petA | IGS |
| T | 10 | 76666 | 76675 | clpP1_1-psbB | IGS |
| A | 10 | 54417 | 54426 | ndhC-trnV-UAC_2 | IGS |
| T | 12 | 74801 | 74812 | clpP1_3-clpP1_2 | Intron |
| A | 12 | 75008 | 75019 | clpP1_3-clpP1_2 | Intron |
| TATT | 3 | 53911 | 53922 | ndhC-trnV-UAC_2 | IGS |
| T | 12 | 117358 | 117369 | ndhF-rpl32 | IGS |
| T | 10 | 117754 | 117763 | ndhF-rpl32 | IGS |
| A | 17 | 46594 | 46610 | pafI_1-trnS-GGA | IGS |
| A | 12 | 46656 | 46667 | pafI_1-trnS-GGA | IGS |
| T | 11 | 19431 | 19441 | rpoC2 | Gene |
| T | 11 | 21860 | 21870 | rpoC1_2 | Gene |
| A | 11 | 47262 | 47272 | pafI_1-trnS-GGA | IGS |
| A | 11 | 65126 | 65136 | pafII-cemA | IGS |
| A | 10 | 64416 | 64425 | pafII-cemA | IGS |
| A | 11 | 80843 | 80853 | petB_2-petD_1 | IGS |
| T | 13 | 82330 | 82342 | petD_2-rpoA | IGS |
| A | 10 | 82350 | 82359 | petD_2-rpoA | IGS |
| T | 12 | 70314 | 70325 | petG-trnW-CCA | IGS |
| AT | 6 | 44076 | 44087 | psaA-pafI_3 | IGS |
| T | 11 | 74711 | 74721 | clpP1_3-clpP1_2 | Intron |
| A | 11 | 74814 | 74824 | clpP1_3-clpP1_2 | Intron |
| ATCT | 3 | 43702 | 43713 | psaA-pafI_3 | IGS |
| T | 11 | 71624 | 71634 | psaJ-rpl33 | IGS |
| A | 11 | 86782 | 86792 | rpl16_2-rpl16_1 | Intron |
| T | 10 | 71297 | 71306 | psaJ-rpl33 | IGS |
| T | 11 | 131692 | 131702 | ycf1 | Gene |
| T | 11 | 132942 | 132952 | ycf1 | Gene |
| T | 10 | 4345 | 4354 | matK-trnK-UUU_1 | Intron |
| T | 11 | 1813 | 1823 | psbA-trnK-UUU_2 | IGS |
| A | 10 | 69540 | 69549 | psbE-petL | IGS |
| T | 11 | 30440 | 30450 | psbM-trnD-GUC | IGS |
| T | 10 | 12646 | 12655 | atpF_2-atpF_1 | Intron |
| TA | 5 | 30690 | 30699 | psbM-trnD-GUC | IGS |
| A | 12 | 37488 | 37499 | psbZ-trnG-GCC | IGS |
| A | 11 | 37605 | 37615 | psbZ-trnG-GCC | IGS |
| ATAA | 3 | 37673 | 37684 | psbZ-trnG-GCC | IGS |
| T | 11 | 60677 | 60687 | rbcL-accD | IGS |
| A | 13 | 160045 | 160057 | rpl2_2-trnH-GUG | IGS |
| T | 10 | 118247 | 118256 | rpl32-trnL-UAG | IGS |
| AT | 6 | 118417 | 118428 | rpl32-trnL-UAG | IGS |
| TA | 5 | 71992 | 72001 | rpl33-rps18 | IGS |
| AATA | 3 | 72085 | 72096 | rpl33-rps18 | IGS |
| A | 14 | 28094 | 28107 | rpoB-trnC-GCA | IGS |
| A | 10 | 28734 | 28743 | rpoB-trnC-GCA | IGS |
| T | 10 | 104066 | 104075 | rps12_2-trnV-GAC | IGS |
| A | 10 | 7094 | 7103 | rps16_1-trnQ-UUG | IGS |
| T | 10 | 74842 | 74851 | clpP1_3-clpP1_2 | Intron |
| A | 10 | 76034 | 76043 | clpP1_2-clpP1_1 | Intron |
| T | 10 | 72684 | 72693 | rps18-rpl20 | IGS |
| AAAT | 3 | 72643 | 72654 | rps18-rpl20 | IGS |
| A | 10 | 86174 | 86183 | rpl16_2-rpl16_1 | Intron |
| A | 10 | 87091 | 87100 | rpl16_2-rpl16_1 | Intron |
| T | 13 | 88966 | 88978 | rps19-rpl2_2 | IGS |
| T | 14 | 17175 | 17188 | rps2-rpoC2 | IGS |
| AT | 6 | 48581 | 48592 | rps4-trnT-UGU | IGS |
| T | 11 | 85067 | 85077 | rps8-rpl14 | IGS |
| A | 10 | 125469 | 125478 | ndhA_2-ndhA_1 | Intron |
| A | 10 | 125675 | 125684 | ndhA_2-ndhA_1 | Intron |
| T | 10 | 130548 | 130557 | ycf1 | Gene |
| T | 10 | 133466 | 133475 | ycf1 | Gene |
| A | 14 | 113185 | 113198 | rrn5-trnR-ACG | IGS |
| T | 10 | 32166 | 32175 | trnE-UUC-trnT-GGU | IGS |
| TA | 6 | 31904 | 31915 | trnE-UUC-trnT-GGU | IGS |
| TA | 6 | 32463 | 32474 | trnE-UUC-trnT-GGU | IGS |
| AT | 6 | 46319 | 46330 | pafI_2-pafI_1 | Intron |
| T | 11 | 9908 | 9918 | trnG-UCC_1-trnS-CGA_2 | IGS |
| T | 16 | 5019 | 5034 | trnK-UUU_1-rps16_2 | IGS |
| A | 14 | 4588 | 4601 | trnK-UUU_1-rps16_2 | IGS |
| T | 10 | 4748 | 4757 | trnK-UUU_1-rps16_2 | IGS |
| T | 10 | 50908 | 50917 | trnL-UAA_2-trnF-GAA | IGS |
| AATA | 3 | 70836 | 70847 | trnP-UGG-psaJ | IGS |
| TC | 5 | 65324 | 65333 | cemA | Gene |
| T | 14 | 135825 | 135838 | trnR-ACG-rrn5 | IGS |
| AT | 5 | 86321 | 86330 | rpl16_2-rpl16_1 | Intron |
| A | 10 | 8699 | 8708 | trnS-GCU-trnS-CGA_1 | IGS |
| T | 10 | 32600 | 32609 | trnT-GGU-psbD | IGS |
| A | 10 | 32683 | 32692 | trnT-GGU-psbD | IGS |
| TGAAA | 3 | 33491 | 33505 | trnT-GGU-psbD | IGS |
| T | 11 | 49307 | 49317 | trnT-UGU-trnL-UAA_1 | IGS |
| T | 10 | 49388 | 49397 | trnT-UGU-trnL-UAA_1 | IGS |
| A | 10 | 49563 | 49572 | trnT-UGU-trnL-UAA_1 | IGS |
| AAAT | 5 | 49835 | 49854 | trnT-UGU-trnL-UAA_1 | IGS |
| A | 10 | 144948 | 144957 | trnV-GAC-rps12_2 | IGS |
| TTGA | 3 | 122645 | 122656 | ndhE | Gene |
| CATT | 3 | 130722 | 130733 | ycf1 | Gene |
