## Supplementary Table S5 for "Complete chloroplast genome of African Baobab (*Adansonia digitata* L.): structural characterization, comparative genomics, and phylogenetic placement within Malvaceae"

**Table S5.** Codon usage metrics for protein-coding genes of *A. digitata* chloroplast genome.

| **Genes** | **Function** | **T3s** | **C3s** | **A3s** | **G3s** | **CAI** | **CBI** | **Fop** | **Nc** | **GC3s** | **GC** | **GC12** | **AT_bias** | **GC_ bias** | **eNC** |
| --- | --- | --- | --- | --- | --- | --- | --- | --- | --- | --- | --- | --- | --- | --- | --- |
| rps14 | Ribosomal protein | 0.347 | 0.133 | 0.476 | 0.273 | 0.127 | -0.159 | 0.323 | 36.810 | 0.323 | 0.420 | 0.469 | 0.578 | 0.672 | 53.864 |
| atpF | ATP synthase | 0.407 | 0.179 | 0.451 | 0.242 | 0.146 | -0.129 | 0.339 | 46.350 | 0.317 | 0.379 | 0.410 | 0.526 | 0.575 | 53.465 |
| atpE | ATP synthase | 0.417 | 0.185 | 0.398 | 0.217 | 0.161 | -0.030 | 0.391 | 52.950 | 0.312 | 0.419 | 0.473 | 0.489 | 0.540 | 53.128 |
| petA | Cytochrome b6f complex | 0.490 | 0.171 | 0.357 | 0.246 | 0.186 | -0.103 | 0.350 | 52.460 | 0.312 | 0.403 | 0.449 | 0.421 | 0.589 | 53.128 |
| psaA | Photosystem I | 0.438 | 0.211 | 0.383 | 0.169 | 0.194 | -0.087 | 0.364 | 50.520 | 0.308 | 0.437 | 0.502 | 0.467 | 0.445 | 52.855 |
| rpl2 | Ribosomal protein | 0.402 | 0.188 | 0.425 | 0.188 | 0.139 | -0.091 | 0.362 | 52.810 | 0.302 | 0.440 | 0.509 | 0.514 | 0.500 | 52.440 |
| pafI | Photosystem assembly | 0.451 | 0.173 | 0.458 | 0.235 | 0.155 | -0.134 | 0.360 | 56.060 | 0.292 | 0.397 | 0.450 | 0.504 | 0.576 | 51.736 |
| atpB | ATP synthase | 0.425 | 0.197 | 0.417 | 0.162 | 0.196 | 0.006 | 0.413 | 47.640 | 0.291 | 0.432 | 0.503 | 0.495 | 0.451 | 51.664 |
| rpoC2 | RNA polymerase | 0.441 | 0.167 | 0.446 | 0.215 | 0.154 | -0.155 | 0.326 | 51.380 | 0.285 | 0.380 | 0.428 | 0.503 | 0.563 | 51.234 |
| ndhJ | NAD(P)H dehydrogenase | 0.476 | 0.175 | 0.420 | 0.200 | 0.176 | -0.098 | 0.349 | 55.510 | 0.282 | 0.407 | 0.470 | 0.468 | 0.534 | 51.018 |
| pafII | Photosystem II | 0.510 | 0.159 | 0.356 | 0.217 | 0.169 | -0.063 | 0.374 | 49.140 | 0.282 | 0.384 | 0.435 | 0.411 | 0.578 | 51.018 |
| rbcL | Rubisco | 0.496 | 0.202 | 0.390 | 0.154 | 0.255 | 0.070 | 0.464 | 49.200 | 0.281 | 0.437 | 0.515 | 0.440 | 0.433 | 50.945 |
| psbA | Photosystem II | 0.536 | 0.253 | 0.298 | 0.072 | 0.321 | 0.226 | 0.550 | 39.400 | 0.281 | 0.418 | 0.487 | 0.357 | 0.222 | 50.945 |
| psaB | Photosystem I | 0.479 | 0.177 | 0.384 | 0.177 | 0.183 | -0.082 | 0.370 | 49.560 | 0.280 | 0.416 | 0.484 | 0.445 | 0.499 | 50.872 |
| clpP1 | Protease subunit | 0.442 | 0.173 | 0.436 | 0.177 | 0.178 | -0.106 | 0.341 | 56.650 | 0.276 | 0.423 | 0.497 | 0.497 | 0.505 | 50.581 |
| rpoB | RNA polymerase | 0.443 | 0.143 | 0.450 | 0.222 | 0.155 | -0.117 | 0.344 | 49.880 | 0.276 | 0.393 | 0.452 | 0.504 | 0.608 | 50.581 |
| ndhB | NAD(P)H dehydrogenase | 0.433 | 0.206 | 0.429 | 0.129 | 0.166 | -0.074 | 0.362 | 47.650 | 0.273 | 0.374 | 0.425 | 0.498 | 0.385 | 50.361 |
| psbD | Photosystem II | 0.507 | 0.203 | 0.365 | 0.126 | 0.249 | 0.042 | 0.441 | 44.180 | 0.272 | 0.426 | 0.503 | 0.419 | 0.383 | 50.288 |
| petB | Cytochrome b6f complex | 0.506 | 0.161 | 0.337 | 0.168 | 0.222 | 0.010 | 0.405 | 46.550 | 0.270 | 0.409 | 0.479 | 0.400 | 0.510 | 50.141 |
| psbB | PSII CP47 antenna protein | 0.491 | 0.171 | 0.382 | 0.151 | 0.182 | -0.087 | 0.364 | 46.380 | 0.262 | 0.434 | 0.520 | 0.437 | 0.469 | 49.548 |
| atpA | ATP synthase | 0.488 | 0.150 | 0.402 | 0.181 | 0.197 | -0.059 | 0.383 | 46.740 | 0.260 | 0.410 | 0.485 | 0.452 | 0.546 | 49.399 |
| cemA | Membrane protein | 0.514 | 0.199 | 0.424 | 0.154 | 0.205 | -0.027 | 0.394 | 43.370 | 0.259 | 0.330 | 0.366 | 0.452 | 0.436 | 49.325 |
| rps8 | Ribosomal protein | 0.417 | 0.165 | 0.457 | 0.167 | 0.103 | -0.053 | 0.372 | 36.330 | 0.256 | 0.373 | 0.432 | 0.523 | 0.502 | 49.100 |
| rpl14 | Ribosomal protein | 0.431 | 0.177 | 0.464 | 0.145 | 0.167 | -0.056 | 0.370 | 49.560 | 0.252 | 0.404 | 0.480 | 0.518 | 0.450 | 48.800 |
| accD | Acetyl-CoA carboxylase | 0.595 | 0.165 | 0.356 | 0.181 | 0.205 | -0.154 | 0.349 | 47.390 | 0.249 | 0.343 | 0.390 | 0.374 | 0.523 | 48.575 |
| rps2 | Ribosomal protein | 0.492 | 0.108 | 0.416 | 0.223 | 0.165 | -0.146 | 0.327 | 50.730 | 0.248 | 0.391 | 0.463 | 0.458 | 0.674 | 48.499 |
| atpI | ATP synthase | 0.481 | 0.198 | 0.403 | 0.102 | 0.180 | -0.050 | 0.370 | 44.270 | 0.248 | 0.383 | 0.451 | 0.456 | 0.341 | 48.499 |
| matK | Maturase | 0.510 | 0.144 | 0.453 | 0.204 | 0.160 | -0.186 | 0.305 | 48.730 | 0.247 | 0.325 | 0.364 | 0.471 | 0.587 | 48.424 |
| rps4 | Ribosomal protein | 0.453 | 0.176 | 0.472 | 0.133 | 0.150 | -0.020 | 0.386 | 51.200 | 0.244 | 0.381 | 0.450 | 0.511 | 0.431 | 48.198 |
| ndhK | NAD(P)H dehydrogenase | 0.476 | 0.173 | 0.429 | 0.129 | 0.148 | -0.169 | 0.313 | 49.910 | 0.244 | 0.393 | 0.468 | 0.474 | 0.427 | 48.198 |
| rpoC1 | RNA polymerase | 0.483 | 0.150 | 0.456 | 0.173 | 0.150 | -0.138 | 0.326 | 48.070 | 0.244 | 0.386 | 0.457 | 0.486 | 0.536 | 48.198 |
| ndhI | NAD(P)H dehydrogenase | 0.515 | 0.181 | 0.446 | 0.144 | 0.212 | -0.095 | 0.363 | 46.570 | 0.244 | 0.363 | 0.423 | 0.465 | 0.443 | 48.198 |
| ndhH | NAD(P)H dehydrogenase | 0.456 | 0.167 | 0.495 | 0.143 | 0.160 | -0.075 | 0.363 | 51.070 | 0.235 | 0.386 | 0.462 | 0.521 | 0.461 | 47.516 |
| rpoA | RNA polymerase | 0.471 | 0.161 | 0.502 | 0.159 | 0.158 | -0.120 | 0.342 | 48.120 | 0.235 | 0.347 | 0.403 | 0.516 | 0.497 | 47.516 |
| rps18 | Ribosomal protein | 0.488 | 0.125 | 0.435 | 0.171 | 0.106 | -0.156 | 0.313 | 36.330 | 0.232 | 0.347 | 0.405 | 0.472 | 0.578 | 47.288 |
| rpl20 | Ribosomal protein | 0.460 | 0.100 | 0.446 | 0.208 | 0.095 | -0.193 | 0.292 | 46.440 | 0.230 | 0.359 | 0.424 | 0.492 | 0.675 | 47.136 |
| rpl22 | Ribosomal protein | 0.403 | 0.151 | 0.543 | 0.149 | 0.143 | -0.077 | 0.368 | 51.340 | 0.229 | 0.382 | 0.459 | 0.574 | 0.495 | 47.059 |
| ndhG | NAD(P)H dehydrogenase | 0.500 | 0.106 | 0.391 | 0.193 | 0.130 | -0.226 | 0.253 | 46.350 | 0.224 | 0.345 | 0.406 | 0.439 | 0.645 | 46.679 |
| ccsA | Cytochrome c synthesis | 0.451 | 0.166 | 0.485 | 0.103 | 0.149 | -0.199 | 0.292 | 45.220 | 0.220 | 0.336 | 0.394 | 0.518 | 0.384 | 46.373 |
| rps3 | Ribosomal protein | 0.450 | 0.169 | 0.532 | 0.133 | 0.145 | -0.143 | 0.333 | 49.530 | 0.219 | 0.356 | 0.425 | 0.542 | 0.441 | 46.297 |
| ycf1 | TIC complex | 0.484 | 0.155 | 0.556 | 0.155 | 0.161 | -0.135 | 0.345 | 46.720 | 0.218 | 0.302 | 0.344 | 0.535 | 0.500 | 46.221 |
| rpl16 | Ribosomal protein | 0.387 | 0.151 | 0.518 | 0.111 | 0.128 | -0.072 | 0.373 | 40.630 | 0.214 | 0.427 | 0.534 | 0.572 | 0.424 | 45.916 |
| rps7 | Ribosomal protein | 0.413 | 0.149 | 0.523 | 0.118 | 0.186 | -0.068 | 0.380 | 45.000 | 0.213 | 0.409 | 0.507 | 0.559 | 0.441 | 45.839 |
| ndhE | NAD(P)H dehydrogenase | 0.562 | 0.112 | 0.360 | 0.164 | 0.145 | -0.186 | 0.299 | 46.130 | 0.206 | 0.323 | 0.382 | 0.391 | 0.593 | 45.305 |
| ndhA | NAD(P)H dehydrogenase | 0.462 | 0.128 | 0.464 | 0.126 | 0.132 | -0.146 | 0.301 | 43.760 | 0.203 | 0.344 | 0.415 | 0.501 | 0.496 | 45.076 |
| rps11 | Ribosomal protein | 0.450 | 0.125 | 0.457 | 0.103 | 0.153 | -0.113 | 0.346 | 43.540 | 0.195 | 0.454 | 0.584 | 0.504 | 0.451 | 44.466 |
| ndhF | NAD(P)H dehydrogenase | 0.533 | 0.103 | 0.437 | 0.161 | 0.133 | -0.242 | 0.264 | 44.260 | 0.194 | 0.319 | 0.382 | 0.450 | 0.611 | 44.390 |
| petD | Cytochrome b6f complex | 0.478 | 0.109 | 0.464 | 0.121 | 0.175 | -0.092 | 0.320 | 39.160 | 0.190 | 0.381 | 0.477 | 0.492 | 0.526 | 44.085 |
| ndhC | NAD(P)H dehydrogenase | 0.520 | 0.100 | 0.443 | 0.133 | 0.186 | -0.076 | 0.342 | 50.770 | 0.180 | 0.361 | 0.452 | 0.460 | 0.571 | 43.326 |
