## Supplementary Table S6 for "Complete chloroplast genome of African Baobab (*Adansonia digitata* L.): structural characterization, comparative genomics, and phylogenetic placement within Malvaceae"

**Table S6:** Predicted RNA editing sites identified across protein-coding genes of the assembled *A. digitata* chloroplast genome, consisting of affected codon position, nucleotide conversion, and resulting amino acid change.

| **Gene** | **No.** | **Step** | **Base** | **Aa** | **Triplet pos.** | **Bases** | **Codon** | **Aa change** | **Label** | **Silenet** |
| --- | --- | --- | --- | --- | --- | --- | --- | --- | --- | --- |
| **matK** | 1 | 1/1 | 89 | 30 | 2 | C→U | GCA→GUA | A→V | matKeU89AV |  |
|  | 2 | 1/1 | 97 | 33 | 1 | C→U | CAU→UAU | H→Y | matKeU97HY |  |
|  | 3 | 1/1 | 232 | 78 | 1 | U→C | UCU→CCU | S→P | matKeC232SP |  |
|  | 4 | 1/1 | 256 | 86 | 1 | C→U | CCA→UCA | P→S | matKeU256PS |  |
|  | 5 | 1/1 | 262 | 88 | 1 | U→C | UUU→CUU | F→L | matKeC262FL |  |
|  | 6 | 1/1 | 302 | 101 | 2 | C→U | GCG→GUG | A→V | matKeU302AV |  |
|  | 7 | 1/1 | 382 | 128 | 1 | C→U | CAU→UAU | H→Y | matKeU382HY |  |
|  | 8 | 1/1 | 430 | 144 | 1 | U→C | UUC→CUC | F→L | matKeC430FL |  |
|  | 9 | 1/1 | 469 | 157 | 1 | C→U | CAC→UAC | H→Y | matKeU469HY |  |
|  | 10 | 1/1 | 511 | 171 | 1 | U→C | UGG→CGG | W→R | matKeC511WR |  |
|  | 11 | 1/1 | 587 | 196 | 2 | C→U | ACU→AUU | T→I | matKeU587TI |  |
|  | 12 | 1/1 | 604 | 202 | 1 | U→C | UUU→CUU | F→L | matKeC604FL |  |
|  | 13 | 1/1 | 622 | 208 | 1 | U→C | UUC→CUC | F→L | matKeC622FL |  |
|  | 14 | 1/1 | 640 | 214 | 1 | C→U | CAU→UAU | H→Y | matKeU640HY |  |
|  | 15 | 1/1 | 644 | 215 | 2 | C→U | GCA→GUA | A→V | matKeU644AV |  |
|  | 16 | 1/1 | 667 | 223 | 1 | C→U | CUU→UUU | L→F | matKeU667LF |  |
|  | 17 | 1/1 | 718 | 240 | 1 | U→C | UUU→CUU | F→L | matKeC718FL |  |
|  | 18 | 1/1 | 721 | 241 | 1 | C→U | CUU→UUU | L→F | matKeU721LF |  |
|  | 19 | 1/1 | 754 | 252 | 1 | U→C | UAU→CAU | Y→H | matKeC754YH |  |
|  | 20 | 1/1 | 757 | 253 | 1 | C→U | CUU→UUU | L→F | matKeU757LF |  |
|  | 21 | 1/1 | 802 | 268 | 1 | U→C | UUC→CUC | F→L | matKeC802FL |  |
|  | 22 | 1/1 | 868 | 290 | 1 | U→C | UCU→CCU | S→P | matKeC868SP |  |
|  | 23 | 1/1 | 919 | 307 | 1 | C→U | CAU→UAU | H→Y | matKeU919HY |  |
|  | 24 | 1/1 | 935 | 312 | 2 | C→U | UCU→UUU | S→F | matKeU935SF |  |
|  | 25 | 1/1 | 1022 | 341 | 2 | C→U | UCA→UUA | S→L | matKeU1022SL |  |
|  | 26 | 1/1 | 1070 | 357 | 2 | C→U | GCU→GUU | A→V | matKeU1070AV |  |
|  | 27 | 1/1 | 1243 | 415 | 1 | C→U | CAC→UAC | H→Y | matKeU1243HY |  |
|  | 28 | 1/1 | 1297 | 433 | 1 | U→C | UUU→CUU | F→L | matKeC1297FL |  |
|  | 29 | 1/1 | 1372 | 458 | 1 | U→C | UUU→CUU | F→L | matKeC1372FL |  |
|  | 30 | 1/1 | 1387 | 463 | 1 | U→C | UUU→CUU | F→L | matKeC1387FL |  |
|  | 31 | 1/1 | 1408 | 470 | 1 | U→C | UUU→CUU | F→L | matKeC1408FL |  |
| **rps16** | 1 | 1/1 | 40 | 14 | 1 | U→C | UGA→CGA | *→R | rps16eC40*R |  |
|  | 2 | 1/1 | 128 | 43 | 2 | U→C | AUU→ACU | I→T | rps16eC128IT |  |
|  | 3 | 1/1 | 151 | 51 | 1 | C→U | CCC→UCC | P→S | rps16eU151PS |  |
|  | 4 | 1/1 | 200 | 67 | 2 | U→C | GUU→GCU | V→A | rps16eC200VA |  |
|  | 5 | 1/1 | 212 | 71 | 2 | U→C | UUA→UCA | L→S | rps16eC212LS |  |
| **psbK** | 2 | 1/1 | 142 | 48 | 1 | U→C | UUU→CUU | F→L | psbKeC142FL |  |
|  | 1 | 1/1 | 136 | 46 | 1 | C→U | CUC→UUC | L→F | psbKeU136LF |  |
| **atpA** | 1 | 1/1 | 127 | 43 | 1 | C→U | CAC→UAC | H→Y | atpAeU127HY |  |
|  | 2 | 1/1 | 637 | 213 | 1 | U→C | UUC→CUC | F→L | atpAeC637FL |  |
|  | 3 | 1/1 | 914 | 305 | 2 | C→U | UCA→UUA | S→L | atpAeU914SL |  |
|  | 4 | 1/1 | 1148 | 383 | 2 | C→U | UCA→UUA | S→L | atpAeU1148SL |  |
|  | 5 | 1/1 | 1444 | 482 | 1 | U→C | UUC→CUC | F→L | atpAeC1444FL |  |
|  | 6 | 1/1 | 1465 | 489 | 1 | C→U | CUU→UUU | L→F | atpAeU1465LF |  |
| **atpF** | 1 | 1/1 | 269 | 90 | 2 | U→C | AUG→ACG | M→T | atpFeC269MT |  |
|  | 2 | 1/1 | 350 | 117 | 2 | U→C | AUU→ACU | I→T | atpFeC350IT |  |
|  | 1 | 1/1 | 92 | 31 | 2 | C→U | CCA→CUA | P→L | atpFeU92PL |  |
|  | 2 | 1/1 | 112 | 38 | 1 | C→U | CUU→UUU | L→F | atpFeU112LF |  |
| **atpI** | 1 | 1/1 | 635 | 212 | 2 | C→U | UCA→UUA | S→L | atpIeU635SL |  |
| **rps2** | 1 | 1/1 | 134 | 45 | 2 | C→U | ACA→AUA | T→I | rps2eU134TI |  |
|  | 2 | 1/1 | 248 | 83 | 2 | C→U | UCG→UUG | S→L | rps2eU248SL |  |
|  | 3 | 1/1 | 325 | 109 | 1 | C→U | CCC→UCC | P→S | rps2eU325PS |  |
|  | 4 | 1/1 | 388 | 130 | 1 | C→U | CUC→UUC | L→F | rps2eU388LF |  |
|  | 5 | 1/1 | 706 | 236 | 1 | C→U | CCU→UCU | P→S | rps2eU706PS |  |
| **rpoC2** | 1 | 1/1 | 668 | 223 | 2 | C→U | ACC→AUC | T→I | rpoC2eU668TI |  |
|  | 2 | 1/1 | 707 | 236 | 2 | C→U | ACG→AUG | T→M | rpoC2eU707TM |  |
|  | 3 | 1/1 | 712 | 238 | 1 | C→U | CCG→UCG | P→S | rpoC2eU712PS |  |
|  | 4 | 1/1 | 775 | 259 | 1 | C→U | CCG→UCG | P→S | rpoC2eU775PS |  |
|  | 5 | 1/1 | 829 | 277 | 1 | U→C | UUC→CUC | F→L | rpoC2eC829FL |  |
|  | 6 | 1/1 | 850 | 284 | 1 | C→U | CCA→UCA | P→S | rpoC2eU850PS |  |
|  | 7 | 1/1 | 1321 | 441 | 1 | C→U | CAU→UAU | H→Y | rpoC2eU1321HY |  |
|  | 8 | 1/1 | 1457 | 486 | 2 | U→C | UUC→UCC | F→S | rpoC2eC1457FS |  |
|  | 9 | 1/1 | 1543 | 515 | 1 | U→C | UUC→CUC | F→L | rpoC2eC1543FL |  |
|  | 10 | 1/1 | 1633 | 545 | 1 | U→C | UUU→CUU | F→L | rpoC2eC1633FL |  |
|  | 11 | 1/1 | 1666 | 556 | 1 | U→C | UUC→CUC | F→L | rpoC2eC1666FL |  |
|  | 12 | 1/1 | 1684 | 562 | 1 | C→U | CUC→UUC | L→F | rpoC2eU1684LF |  |
|  | 13 | 1/1 | 1748 | 583 | 2 | U→C | UUG→UCG | L→S | rpoC2eC1748LS |  |
|  | 14 | 1/1 | 1777 | 593 | 1 | U→C | UCA→CCA | S→P | rpoC2eC1777SP |  |
|  | 15 | 1/1 | 1837 | 613 | 1 | C→U | CUU→UUU | L→F | rpoC2eU1837LF |  |
|  | 16 | 1/1 | 2026 | 676 | 1 | U→C | UCC→CCC | S→P | rpoC2eC2026SP |  |
|  | 17 | 1/1 | 2096 | 699 | 2 | C→U | ACA→AUA | T→I | rpoC2eU2096TI |  |
|  | 18 | 1/1 | 2396 | 799 | 2 | C→U | CCG→CUG | P→L | rpoC2eU2396PL |  |
|  | 19 | 1/1 | 2551 | 851 | 1 | U→C | UGU→CGU | C→R | rpoC2eC2551CR |  |
|  | 20 | 1/1 | 2558 | 853 | 2 | C→U | UCC→UUC | S→F | rpoC2eU2558SF |  |
|  | 21 | 1/1 | 2630 | 877 | 2 | U→C | UUU→UCU | F→S | rpoC2eC2630FS |  |
|  | 22 | 1/1 | 2653 | 885 | 1 | C→U | CCG→UCG | P→S | rpoC2eU2653PS |  |
|  | 23 | 1/1 | 2656 | 886 | 1 | U→C | UCA→CCA | S→P | rpoC2eC2656SP |  |
|  | 24 | 1/1 | 2710 | 904 | 1 | U→C | UCC→CCC | S→P | rpoC2eC2710SP |  |
|  | 25 | 1/1 | 2737 | 913 | 1 | U→C | UUU→CUU | F→L | rpoC2eC2737FL |  |
|  | 26 | 1/1 | 2765 | 922 | 2 | C→U | ACG→AUG | T→M | rpoC2eU2765TM |  |
|  | 27 | 1/1 | 2767 | 923 | 1 | C→U | CUC→UUC | L→F | rpoC2eU2767LF |  |
|  | 28 | 1/1 | 2854 | 952 | 1 | U→C | UAU→CAU | Y→H | rpoC2eC2854YH |  |
|  | 29 | 1/1 | 2912 | 971 | 2 | U→C | UUA→UCA | L→S | rpoC2eC2912LS |  |
|  | 30 | 1/1 | 2915 | 972 | 2 | U→C | UUG→UCG | L→S | rpoC2eC2915LS |  |
|  | 31 | 1/1 | 2977 | 993 | 1 | C→U | CAU→UAU | H→Y | rpoC2eU2977HY |  |
|  | 32 | 1/1 | 2987 | 996 | 2 | C→U | ACC→AUC | T→I | rpoC2eU2987TI |  |
|  | 33 | 1/1 | 3029 | 1010 | 2 | C→U | ACU→AUU | T→I | rpoC2eU3029TI |  |
|  | 34 | 1/1 | 3082 | 1028 | 1 | U→C | UUC→CUC | F→L | rpoC2eC3082FL |  |
|  | 35 | 1/1 | 3137 | 1046 | 2 | C→U | CCC→CUC | P→L | rpoC2eU3137PL |  |
|  | 36 | 1/1 | 3259 | 1087 | 1 | U→C | UUU→CUU | F→L | rpoC2eC3259FL |  |
|  | 37 | 1/1 | 3281 | 1094 | 2 | U→C | GUA→GCA | V→A | rpoC2eC3281VA |  |
|  | 38 | 1/1 | 4042 | 1348 | 1 | U→C | UUC→CUC | F→L | rpoC2eC4042FL |  |
|  | 39 | 1/1 | 4069 | 1357 | 1 | U→C | UUC→CUC | F→L | rpoC2eC4069FL |  |
|  | 40 | 1/1 | 4075 | 1359 | 1 | C→U | CAC→UAC | H→Y | rpoC2eU4075H |  |
| **rpoC2** | 1 | 1/1 | 488 | 163 | 2 | C→U | UCA→UUA | S→L | rpoC1eU488SL |  |
|  | 2 | 1/1 | 557 | 186 | 2 | C→U | ACA→AUA | T→I | rpoC1eU557TI |  |
|  | 3 | 1/1 | 680 | 227 | 2 | U→C | CUC→CCC | L→P | rpoC1eC680LP |  |
|  | 4 | 1/1 | 1261 | 421 | 1 | C→U | CCG→UCG | P→S | rpoC1eU1261PS |  |
|  | 5 | 1/1 | 1831 | 611 | 1 | U→C | UGC→CGC | C→R | rpoC1eC1831CR |  |
|  | 6 | 1/1 | 1853 | 618 | 2 | C→U | GCU→GUU | A→V | rpoC1eU1853AV |  |
|  | 7 | 1/1 | 1942 | 648 | 1 | C→U | CUU→UUU | L→F | rpoC1eU1942LF |  |
|  | 8 | 1/1 | 1981 | 661 | 1 | C→U | CUU→UUU | L→F | rpoC1eU1981LF |  |
|  | 9 | 1/1 | 2041 | 681 | 1 | C→U | CAA→UAA | Q→* | rpoC1eU2041Q* |  |
|  | 1 | 1/1 | 41 | 14 | 2 | C→U | UCA→UUA | S→L | rpoC1eU41SL |  |
|  | 2 | 1/1 | 98 | 33 | 2 | C→U | ACC→AUC | T→I | rpoC1eU98TI |  |
| **rpoB** | 1 | 1/1 | 26 | 9 | 2 | U→C | AUG→ACG | M→T | rpoBeC26MT |  |
|  | 2 | 1/1 | 95 | 32 | 2 | C→U | ACA→AUA | T→I | rpoBeU95TI |  |
|  | 3 | 1/1 | 131 | 44 | 2 | C→U | ACA→AUA | T→I | rpoBeU131TI |  |
|  | 4 | 1/1 | 338 | 113 | 2 | C→U | UCU→UUU | S→F | rpoBeU338SF |  |
|  | 5 | 1/1 | 551 | 184 | 2 | C→U | UCA→UUA | S→L | rpoBeU551SL |  |
|  | 6 | 1/1 | 566 | 189 | 2 | C→U | UCG→UUG | S→L | rpoBeU566SL |  |
|  | 7 | 1/1 | 836 | 279 | 2 | C→U | ACA→AUA | T→I | rpoBeU836TI |  |
|  | 8 | 1/1 | 973 | 325 | 1 | U→C | UUC→CUC | F→L | rpoBeC973FL |  |
|  | 9 | 1/1 | 1493 | 498 | 2 | U→C | GUU→GCU | V→A | rpoBeC1493VA |  |
|  | 10 | 1/1 | 2032 | 678 | 1 | C→U | CGU→UGU | R→C | rpoBeU2032RC |  |
|  | 11 | 1/1 | 2414 | 805 | 2 | C→U | ACA→AUA | T→I | rpoBeU2414TI |  |
|  | 12 | 1/1 | 2432 | 811 | 2 | C→U | UCA→UUA | S→L | rpoBeU2432SL |  |
|  | 13 | 1/1 | 3215 | 1072 | 2 | C→U | GCU→GUU | A→V | rpoBeU3215AV |  |
| **psbZ** | 1 | 1/1 | 50 | 17 | 2 | C→U | UCA→UUA | S→L | psbZeU50SL |  |
|  | 2 | 1/1 | 101 | 34 | 2 | U→C | UUG→UCG | L→S | psbZeC101LS |  |
| **psaB** | 1 | 1/1 | 329 | 110 | 2 | C→U | CCU→CUU | P→L | psaBeU329PL |  |
|  | 2 | 1/1 | 673 | 225 | 1 | U→C | UUU→CUU | F→L | psaBeC673FL |  |
| **psaA** | 1 | 1/1 | 1481 | 494 | 2 | C→U | GCA→GUA | A→V | psaAeU1481AV |  |
|  | 2 | 1/1 | 2008 | 670 | 1 | C→U | CUU→UUU | L→F | psaAeU2008LF |  |
|  | 1 | 1/1 | 112 | 38 | 1 | C→U | CCC→UCC | P→S | rps4eU112PS |  |
|  | 2 | 1/1 | 305 | 102 | 2 | C→U | UCG→UUG | S→L | rps4eU305SL |  |
|  | 3 | 1/1 | 365 | 122 | 2 | C→U | ACA→AUA | T→I | rps4eU365TI |  |
|  | 4 | 1/1 | 413 | 138 | 2 | C→U | GCG→GUG | A→V | rps4eU413AV |  |
| **ndhJ** | 1 | 1/1 | 29 | 10 | 2 | U→C | GUC→GCC | V→A | ndhJeC29VA |  |
|  | 2 | 1/1 | 472 | 158 | 1 | C→U | CAU→UAU | H→Y | ndhJeU472HY |  |
| **mdhK** | 1 | 1/1 | 484 | 162 | 1 | C→U | CAU→UAU | H→Y | ndhKeU484HY |  |
|  | 2 | 1/1 | 502 | 168 | 1 | U→C | UCU→CCU | S→P | ndhKeC502SP |  |
|  | 3 | 1/1 | 566 | 189 | 2 | C→U | ACU→AUU | T→I | ndhKeU566TI |  |
|  | 4 | 1/1 | 601 | 201 | 1 | C→U | CCA→UCA | P→S | ndhKeU601PS |  |
|  | 5 | 1/1 | 619 | 207 | 1 | C→U | CCU→UCU | P→S | ndhKeU619PS |  |
|  | 6 | 1/1 | 649 | 217 | 1 | U→C | UCA→CCA | S→P | ndhKeC649SP |  |
| **ndhC** | 1 | 1/1 | 274 | 92 | 1 | C→U | CCC→UCC | P→S | ndhCeU274PS |  |
|  | 2 | 1/1 | 278 | 93 | 2 | U→C | GUA→GCA | V→A | ndhCeC278VA |  |
|  | 3 | 1/1 | 323 | 108 | 2 | C→U | UCA→UUA | S→L | ndhCeU323SL |  |
| **atpB** | 1 | 1/1 | 8 | 3 | 2 | U→C | AUA→ACA | I→T | atpBeC8IT |  |
|  | 2 | 1/1 | 281 | 94 | 2 | C→U | ACG→AUG | T→M | atpBeU281TM |  |
|  | 3 | 1/1 | 403 | 135 | 1 | C→U | CCU→UCU | P→S | atpBeU403PS |  |
| **rbcL** | 1 | 1/1 | 29 | 10 | 2 | U→C | UUU→UCU | F→S | rbcLeC29FS |  |
|  | 2 | 1/1 | 434 | 145 | 2 | U→C | AUU→ACU | I→T | rbcLeC434IT |  |
|  | 3 | 1/1 | 785 | 262 | 2 | C→U | GCU→GUU | A→V | rbcLeU785AV |  |
| **accD** | 1 | 1/1 | 82 | 28 | 1 | C→U | CUU→UUU | L→F | accDeU82LF |  |
|  | 2 | 1/1 | 134 | 45 | 2 | C→U | ACG→AUG | T→M | accDeU134TM |  |
|  | 3 | 1/1 | 148 | 50 | 1 | C→U | CAU→UAU | H→Y | accDeU148HY |  |
|  | 4 | 1/1 | 410 | 137 | 2 | C→U | CCC→CUC | P→L | accDeU410PL |  |
|  | 5 | 1/2 | 421 | 141 | 1 | U→C | UUU→CCU | F→P | accDeCC421FP |  |
|  | 5 | 2/2 | 422 | 141 | 2 | U→C | UUU→CCU | F→P | accDeCC421FP |  |
|  | 6 | 1/1 | 484 | 162 | 1 | C→U | CUU→UUU | L→F | accDeU484LF |  |
|  | 7 | 1/1 | 536 | 179 | 2 | C→U | UCC→UUC | S→F | accDeU536SF |  |
|  | 8 | 1/1 | 638 | 213 | 2 | C→U | ACU→AUU | T→I | accDeU638TI |  |
|  | 9 | 1/1 | 652 | 218 | 1 | C→U | CUC→UUC | L→F | accDeU652LF |  |
|  | 10 | 1/1 | 754 | 252 | 1 | U→C | UAU→CAU | Y→H | accDeC754YH |  |
|  | 11 | 1/1 | 757 | 253 | 1 | C→U | CAU→UAU | H→Y | accDeU757HY |  |
|  | 12 | 1/1 | 794 | 265 | 2 | C→U | UCG→UUG | S→L | accDeU794SL |  |
|  | 13 | 1/1 | 1154 | 385 | 2 | C→U | GCU→GUU | A→V | accDeU1154AV |  |
|  | 14 | 1/1 | 1363 | 455 | 1 | U→C | UUC→CUC | F→L | accDeC1363FL |  |
|  | 15 | 1/1 | 1378 | 460 | 1 | U→C | UUC→CUC | F→L | accDeC1378FL |  |
|  | 16 | 1/1 | 1403 | 468 | 2 | C→U | CCU→CUU | P→L | accDeU1403 |  |
| **psaI** | 1 | 1/1 | 19 | 7 | 1 | U→C | UUU→CUU | F→L | psaIeC19FL |  |
|  | 2 | 1/1 | 83 | 28 | 2 | C→U | UCU→UUU | S→F | psaIeU83SF |  |
|  | 3 | 1/1 | 88 | 30 | 1 | U→C | UAU→CAU | Y→H | psaIeC88YH |  |
| **ycf4** | 1 | 1/1 | 88 | 30 | 1 | U→C | UUU→CUU | F→L | ycf4eC88FL |  |
|  | 2 | 1/1 | 253 | 85 | 1 | U→C | UUU→CUU | F→L | ycf4eC253FL |  |
|  | 3 | 1/1 | 307 | 103 | 1 | U→C | UGU→CGU | C→R | ycf4eC307CR |  |
|  | 4 | 1/1 | 361 | 121 | 1 | C→U | CUU→UUU | L→F | ycf4eU361LF |  |
|  | 5 | 1/1 | 464 | 155 | 2 | C→U | ACU→AUU | T→I | ycf4eU464TI |  |
| **cemA** | 1 | 1/1 | 23 | 8 | 2 | C→U | ACU→AUU | T→I | cemAeU23TI |  |
|  | 2 | 1/1 | 28 | 10 | 1 | C→U | CUU→UUU | L→F | cemAeU28LF |  |
|  | 3 | 1/1 | 37 | 13 | 1 | C→U | CUU→UUU | L→F | cemAeU37LF |  |
|  | 4 | 1/1 | 250 | 84 | 1 | C→U | CUU→UUU | L→F | cemAeU250LF |  |
| **petA** | 1 | 1/1 | 698 | 233 | 2 | U→C | UUG→UCG | L→S | petAeC698LS |  |
|  | 2 | 1/1 | 737 | 246 | 2 | C→U | CCA→CUA | P→L | petAeU737PL |  |
|  | 3 | 1/1 | 892 | 298 | 1 | U→C | UUU→CUU | F→L | petAeC892FL |  |
| **psbJ** | 1 | 1/1 | 59 | 20 | 2 | C→U | CCU→CUU | P→L | psbJeU59PL |  |
| **psbF** | 1 | 1/1 | 77 | 26 | 2 | C→U | UCU→UUU | S→F | psbFeU77SF |  |
| **petL** | 1 | 1/1 | 5 | 2 | 2 | C→U | CCU→CUU | P→L | petLeU5PL |  |
|  | 2 | 1/1 | 59 | 20 | 2 | C→U | GCU→GUU | A→V | petLeU59AV |  |
| **petG** | 1 | 1/1 | 109 | 37 | 1 | C→U | CUU→UUU | L→F | petGeU109LF |  |
| **psaJ** | 1 | 1/1 | 94 | 32 | 1 | U→C | UUC→CUC | F→L | psaJeC94FL |  |
| **rps18** | 1 | 1/1 | 221 | 74 | 2 | C→U | UCG→UUG | S→L | rps18eU221SL |  |
| **rpl20** | 1 | 1/1 | 224 | 75 | 2 | C→U | UCC→UUC | S→F | rpl20eU224SF |  |
|  | 2 | 1/1 | 308 | 103 | 2 | C→U | UCA→UUA | S→L | rpl20eU308SL |  |
|  | 3 | 1/1 | 329 | 110 | 2 | U→C | AUG→ACG | M→T | rpl20eC329MT |  |
| **rps12** | 1 | 1/1 | 221 | 74 | 2 | C→U | UCA→UUA | S→L | rps12eU221SL |  |
|  | 1 | 1/1 | 221 | 74 | 2 | C→U | UCA→UUA | S→L | rps12eU221SL |  |
| **clpP** | 1 | 1/1 | 97 | 33 | 1 | C→U | CUU→UUU | L→F | clpPeU97LF |  |
|  | 2 | 1/1 | 226 | 76 | 1 | C→U | CCU→UCU | P→S | clpPeU226PS |  |
|  | 3 | 1/1 | 323 | 108 | 2 | C→U | GCC→GUC | A→V | clpPeU323AV |  |
|  | 1 | 1/1 | 559 | 187 | 1 | C→U | CAU→UAU | H→Y | clpPeU559HY |  |
| **psbB** | 1 | 1/1 | 1520 | 507 | 2 | U→C | GUA→GCA | V→A | psbBeC1520VA |  |
| **psbT** | 1 | 1/1 | 88 | 30 | 1 | C→U | CCA→UCA | P→S | psbTeU88PS |  |
| **psbN** | 1 | 1/1 | 29 | 10 | 2 | C→U | UCU→UUU | S→F | psbNeU29SF |  |
| **psbH** | 1 | 1/1 | 64 | 22 | 1 | U→C | UUU→CUU | F→L | psbHeC64FL |  |
| **petD** | 1 | 1/1 | 416 | 139 | 2 | U→C | GUG→GCG | V→A | petDeC416VA |  |
| **rpoA** | 1 | 1/1 | 200 | 67 | 2 | C→U | UCU→UUU | S→F | rpoAeU200SF |  |
|  | 2 | 1/1 | 233 | 78 | 2 | U→C | GUA→GCA | V→A | rpoAeC233VA |  |
|  | 3 | 1/1 | 319 | 107 | 1 | U→C | UUU→CUU | F→L | rpoAeC319FL |  |
|  | 4 | 1/1 | 329 | 110 | 2 | C→U | GCC→GUC | A→V | rpoAeU329AV |  |
|  | 5 | 1/1 | 482 | 161 | 2 | C→U | ACG→AUG | T→M | rpoAeU482TM |  |
|  | 6 | 1/1 | 484 | 162 | 1 | C→U | CCA→UCA | P→S | rpoAeU484PS |  |
|  | 7 | 1/1 | 695 | 232 | 2 | C→U | GCG→GUG | A→V | rpoAeU695AV |  |
|  | 8 | 1/1 | 712 | 238 | 1 | C→U | CAU→UAU | H→Y | rpoAeU712HY |  |
|  | 9 | 1/1 | 773 | 258 | 2 | C→U | GCU→GUU | A→V | rpoAeU773AV |  |
|  | 10 | 1/1 | 836 | 279 | 2 | C→U | UCA→UUA | S→L | rpoAeU836SL |  |
|  | 11 | 1/1 | 893 | 298 | 2 | C→U | UCA→UUA | S→L | rpoAeU893SL |  |
| **rps11** | 1 | 1/1 | 17 | 6 | 2 | C→U | CCA→CUA | P→L | rps11eU17PL |  |
|  | 2 | 1/1 | 65 | 22 | 2 | C→U | GCA→GUA | A→V | rps11eU65AV |  |
| **rps8** | 1 | 1/1 | 214 | 72 | 1 | C→U | CCA→UCA | P→S | rps8eU214PS |  |
|  | 2 | 1/1 | 217 | 73 | 1 | C→U | CAU→UAU | H→Y | rps8eU217HY |  |
| **rpl1** | 1 | 1/1 | 19 | 7 | 1 | C→U | CAU→UAU | H→Y | rpl14eU19HY |  |
|  | 2 | 1/1 | 224 | 75 | 2 | U→C | AUG→ACG | M→T | rpl14eC224MT |  |
| **rpl22** | 1 | 1/1 | 25 | 9 | 1 | C→U | CCA→UCA | P→S | rpl22eU25PS |  |
|  | 2 | 1/1 | 55 | 19 | 1 | C→U | CAU→UAU | H→Y | rpl22eU55HY |  |
|  | 3 | 1/1 | 314 | 105 | 2 | U→C | CUG→CCG | L→P | rpl22eC314LP |  |
|  | 4 | 1/1 | 325 | 109 | 1 | C→U | CCC→UCC | P→S | rpl22eU325PS |  |
|  | 5 | 1/1 | 347 | 116 | 2 | C→U | GCA→GUA | A→V | rpl22eU347AV |  |
|  | 6 | 1/1 | 427 | 143 | 1 | U→C | UGA→CGA | *→R | rpl22eC427*R |  |
|  | 7 | 1/1 | 449 | 150 | 2 | U→C | AUA→ACA | I→T | rpl22eC449IT |  |
| **rpl12** | 1 | 1/1 | 301 | 101 | 1 | C→U | CAU→UAU | H→Y | rpl2eU301HY |  |
|  | 2 | 1/2 | 361 | 121 | 1 | U→C | UUG→CCG | L→P | rpl2eCC361LP | X |
|  | 2 | 2/2 | 362 | 121 | 2 | U→C | UUG→CCG | L→P | rpl2eCC361LP |  |
|  | 3 | 1/2 | 382 | 128 | 1 | U→C | UCU→CUU | S→L | rpl2eCU382SL |  |
|  | 3 | 2/2 | 383 | 128 | 2 | C→U | UCU→CUU | S→L | rpl2eCU382SL |  |
|  | 4 | 1/1 | 389 | 130 | 2 | C→U | UCA→UUA | S→L | rpl2eU389SL |  |
|  | 5 | 1/1 | 734 | 245 | 2 | C→U | GCA→GUA | A→V | rpl2eU734AV |  |
|  | 1 | 1/1 | 301 | 101 | 1 | C→U | CAU→UAU | H→Y | rpl2eU301HY |  |
|  | 2 | 1/2 | 361 | 121 | 1 | U→C | UUG→CCG | L→P | rpl2eCC361LP | X |
|  | 2 | 2/2 | 362 | 121 | 2 | U→C | UUG→CCG | L→P | rpl2eCC361LP |  |
|  | 3 | 1/2 | 382 | 128 | 1 | U→C | UCU→CUU | S→L | rpl2eCU382SL |  |
|  | 3 | 2/2 | 383 | 128 | 2 | C→U | UCU→CUU | S→L | rpl2eCU382SL |  |
|  | 4 | 1/1 | 389 | 130 | 2 | C→U | UCA→UUA | S→L | rpl2eU389SL |  |
|  | 5 | 1/1 | 734 | 245 | 2 | C→U | GCA→GUA | A→V | rpl2eU734AV |  |
| **rpl123** | 1 | 1/1 | 71 | 24 | 2 | C→U | UCU→UUU | S→F | rpl23eU71SF |  |
|  | 2 | 1/1 | 89 | 30 | 2 | C→U | UCA→UUA | S→L | rpl23eU89SL |  |
|  | 1 | 1/1 | 71 | 24 | 2 | C→U | UCU→UUU | S→F | rpl23eU71SF |  |
|  | 2 | 1/1 | 89 | 30 | 2 | C→U | UCA→UUA | S→L | rpl23eU89SL |  |
| **ycf2** | 1 | 1/1 | 188 | 63 | 2 | C→U | UCA→UUA | S→L | ycf2eU188SL |  |
|  | 2 | 1/1 | 260 | 87 | 2 | U→C | GUC→GCC | V→A | ycf2eC260VA |  |
|  | 3 | 1/1 | 368 | 123 | 2 | U→C | UUG→UCG | L→S | ycf2eC368LS |  |
|  | 4 | 1/1 | 526 | 176 | 1 | C→U | CGG→UGG | R→W | ycf2eU526RW |  |
|  | 5 | 1/1 | 572 | 191 | 2 | C→U | UCU→UUU | S→F | ycf2eU572SF |  |
|  | 6 | 1/1 | 676 | 226 | 1 | U→C | UCG→CCG | S→P | ycf2eC676SP |  |
|  | 7 | 1/1 | 1199 | 400 | 2 | C→U | UCC→UUC | S→F | ycf2eU1199SF |  |
|  | 8 | 1/1 | 1387 | 463 | 1 | U→C | UCC→CCC | S→P | ycf2eC1387SP |  |
|  | 9 | 1/1 | 1688 | 563 | 2 | C→U | CCG→CUG | P→L | ycf2eU1688PL |  |
|  | 10 | 1/1 | 1970 | 657 | 2 | C→U | UCC→UUC | S→F | ycf2eU1970SF |  |
|  | 11 | 1/1 | 2000 | 667 | 2 | C→U | UCG→UUG | S→L | ycf2eU2000SL |  |
|  | 12 | 1/1 | 2245 | 749 | 1 | U→C | UCC→CCC | S→P | ycf2eC2245SP |  |
|  | 13 | 1/1 | 2536 | 846 | 1 | C→U | CGU→UGU | R→C | ycf2eU2536RC |  |
|  | 14 | 1/1 | 2872 | 958 | 1 | C→U | CCG→UCG | P→S | ycf2eU2872PS |  |
|  | 15 | 1/1 | 3242 | 1081 | 2 | U→C | AUA→ACA | I→T | ycf2eC3242IT |  |
|  | 16 | 1/1 | 3251 | 1084 | 2 | U→C | UUG→UCG | L→S | ycf2eC3251LS |  |
|  | 17 | 1/1 | 3307 | 1103 | 1 | C→U | CCA→UCA | P→S | ycf2eU3307PS |  |
|  | 18 | 1/1 | 3470 | 1157 | 2 | C→U | ACA→AUA | T→I | ycf2eU3470TI |  |
|  | 19 | 1/1 | 3530 | 1177 | 2 | C→U | CCA→CUA | P→L | ycf2eU3530PL |  |
|  | 20 | 1/1 | 3721 | 1241 | 1 | C→U | CGG→UGG | R→W | ycf2eU3721RW |  |
|  | 21 | 1/1 | 3754 | 1252 | 1 | C→U | CUU→UUU | L→F | ycf2eU3754LF |  |
|  | 22 | 1/1 | 3875 | 1292 | 2 | C→U | UCA→UUA | S→L | ycf2eU3875SL |  |
|  | 23 | 1/1 | 3922 | 1308 | 1 | U→C | UGG→CGG | W→R | ycf2eC3922WR |  |
|  | 24 | 1/1 | 3964 | 1322 | 1 | U→C | UUU→CUU | F→L | ycf2eC3964FL |  |
|  | 25 | 1/1 | 4006 | 1336 | 1 | C→U | CCA→UCA | P→S | ycf2eU4006PS |  |
|  | 26 | 1/1 | 4126 | 1376 | 1 | C→U | CAU→UAU | H→Y | ycf2eU4126HY |  |
|  | 27 | 1/1 | 4390 | 1464 | 1 | C→U | CUC→UUC | L→F | ycf2eU4390LF |  |
|  | 28 | 1/1 | 4445 | 1482 | 2 | U→C | AUU→ACU | I→T | ycf2eC4445IT |  |
|  | 29 | 1/1 | 5045 | 1682 | 2 | U→C | CUU→CCU | L→P | ycf2eC5045LP |  |
|  | 30 | 1/1 | 5056 | 1686 | 1 | C→U | CUU→UUU | L→F | ycf2eU5056LF |  |
|  | 31 | 1/1 | 5180 | 1727 | 2 | C→U | UCG→UUG | S→L | ycf2eU5180SL |  |
|  | 32 | 1/1 | 5699 | 1900 | 2 | U→C | UUG→UCG | L→S | ycf2eC5699LS |  |
|  | 33 | 1/1 | 5747 | 1916 | 2 | U→C | GUA→GCA | V→A | ycf2eC5747VA |  |
|  | 34 | 1/1 | 6100 | 2034 | 1 | C→U | CUU→UUU | L→F | ycf2eU6100LF |  |
|  | 35 | 1/1 | 6292 | 2098 | 1 | U→C | UGG→CGG | W→R | ycf2eC6292WR |  |
|  | 36 | 1/1 | 6856 | 2286 | 1 | C→U | CCG→UCG | P→S | ycf2eU6856PS |  |
|  | 1 | 1/1 | 188 | 63 | 2 | C→U | UCA→UUA | S→L | ycf2eU188SL |  |
|  | 2 | 1/1 | 260 | 87 | 2 | U→C | GUC→GCC | V→A | ycf2eC260VA |  |
|  | 3 | 1/1 | 368 | 123 | 2 | U→C | UUG→UCG | L→S | ycf2eC368LS |  |
|  | 4 | 1/1 | 526 | 176 | 1 | C→U | CGG→UGG | R→W | ycf2eU526RW |  |
|  | 5 | 1/1 | 572 | 191 | 2 | C→U | UCU→UUU | S→F | ycf2eU572SF |  |
|  | 6 | 1/1 | 676 | 226 | 1 | U→C | UCG→CCG | S→P | ycf2eC676SP |  |
|  | 7 | 1/1 | 1199 | 400 | 2 | C→U | UCC→UUC | S→F | ycf2eU1199SF |  |
|  | 8 | 1/1 | 1387 | 463 | 1 | U→C | UCC→CCC | S→P | ycf2eC1387SP |  |
|  | 9 | 1/1 | 1688 | 563 | 2 | C→U | CCG→CUG | P→L | ycf2eU1688PL |  |
|  | 10 | 1/1 | 1970 | 657 | 2 | C→U | UCC→UUC | S→F | ycf2eU1970SF |  |
|  | 11 | 1/1 | 2000 | 667 | 2 | C→U | UCG→UUG | S→L | ycf2eU2000SL |  |
|  | 12 | 1/1 | 2245 | 749 | 1 | U→C | UCC→CCC | S→P | ycf2eC2245SP |  |
|  | 13 | 1/1 | 2536 | 846 | 1 | C→U | CGU→UGU | R→C | ycf2eU2536RC |  |
|  | 14 | 1/1 | 2872 | 958 | 1 | C→U | CCG→UCG | P→S | ycf2eU2872PS |  |
|  | 15 | 1/1 | 3242 | 1081 | 2 | U→C | AUA→ACA | I→T | ycf2eC3242IT |  |
|  | 16 | 1/1 | 3251 | 1084 | 2 | U→C | UUG→UCG | L→S | ycf2eC3251LS |  |
|  | 17 | 1/1 | 3307 | 1103 | 1 | C→U | CCA→UCA | P→S | ycf2eU3307PS |  |
|  | 18 | 1/1 | 3470 | 1157 | 2 | C→U | ACA→AUA | T→I | ycf2eU3470TI |  |
|  | 19 | 1/1 | 3530 | 1177 | 2 | C→U | CCA→CUA | P→L | ycf2eU3530PL |  |
|  | 20 | 1/1 | 3721 | 1241 | 1 | C→U | CGG→UGG | R→W | ycf2eU3721RW |  |
|  | 21 | 1/1 | 3754 | 1252 | 1 | C→U | CUU→UUU | L→F | ycf2eU3754LF |  |
|  | 22 | 1/1 | 3875 | 1292 | 2 | C→U | UCA→UUA | S→L | ycf2eU3875SL |  |
|  | 23 | 1/1 | 3922 | 1308 | 1 | U→C | UGG→CGG | W→R | ycf2eC3922WR |  |
|  | 24 | 1/1 | 3964 | 1322 | 1 | U→C | UUU→CUU | F→L | ycf2eC3964FL |  |
|  | 25 | 1/1 | 4006 | 1336 | 1 | C→U | CCA→UCA | P→S | ycf2eU4006PS |  |
|  | 26 | 1/1 | 4126 | 1376 | 1 | C→U | CAU→UAU | H→Y | ycf2eU4126HY |  |
|  | 27 | 1/1 | 4390 | 1464 | 1 | C→U | CUC→UUC | L→F | ycf2eU4390LF |  |
|  | 28 | 1/1 | 4445 | 1482 | 2 | U→C | AUU→ACU | I→T | ycf2eC4445IT |  |
|  | 29 | 1/1 | 5045 | 1682 | 2 | U→C | CUU→CCU | L→P | ycf2eC5045LP |  |
|  | 30 | 1/1 | 5056 | 1686 | 1 | C→U | CUU→UUU | L→F | ycf2eU5056LF |  |
|  | 31 | 1/1 | 5180 | 1727 | 2 | C→U | UCG→UUG | S→L | ycf2eU5180SL |  |
|  | 32 | 1/1 | 5699 | 1900 | 2 | U→C | UUG→UCG | L→S | ycf2eC5699LS |  |
|  | 33 | 1/1 | 5747 | 1916 | 2 | U→C | GUA→GCA | V→A | ycf2eC5747VA |  |
|  | 34 | 1/1 | 6100 | 2034 | 1 | C→U | CUU→UUU | L→F | ycf2eU6100LF |  |
|  | 35 | 1/1 | 6292 | 2098 | 1 | U→C | UGG→CGG | W→R | ycf2eC6292WR |  |
|  | 36 | 1/1 | 6856 | 2286 | 1 | C→U | CCG→UCG | P→S | ycf2eU6856PS |  |
| **ndhB** | 1 | 1/1 | 149 | 50 | 2 | C→U | UCA→UUA | S→L | ndhBeU149SL |  |
|  | 2 | 1/1 | 467 | 156 | 2 | C→U | CCA→CUA | P→L | ndhBeU467PL |  |
|  | 3 | 1/1 | 542 | 181 | 2 | C→U | ACG→AUG | T→M | ndhBeU542TM |  |
|  | 4 | 1/1 | 586 | 196 | 1 | C→U | CAU→UAU | H→Y | ndhBeU586HY |  |
|  | 5 | 1/1 | 737 | 246 | 2 | C→U | CCA→CUA | P→L | ndhBeU737PL |  |
|  | 6 | 1/1 | 746 | 249 | 2 | C→U | UCU→UUU | S→F | ndhBeU746SF |  |
|  | 1 | 1/1 | 149 | 50 | 2 | C→U | UCA→UUA | S→L | ndhBeU149SL |  |
|  | 2 | 1/1 | 467 | 156 | 2 | C→U | CCA→CUA | P→L | ndhBeU467PL |  |
|  | 3 | 1/1 | 542 | 181 | 2 | C→U | ACG→AUG | T→M | ndhBeU542TM |  |
|  | 4 | 1/1 | 586 | 196 | 1 | C→U | CAU→UAU | H→Y | ndhBeU586HY |  |
|  | 5 | 1/1 | 737 | 246 | 2 | C→U | CCA→CUA | P→L | ndhBeU737PL |  |
|  | 6 | 1/1 | 746 | 249 | 2 | C→U | UCU→UUU | S→F | ndhBeU746SF |  |
|  | 1 | 1/1 | 830 | 277 | 2 | C→U | UCG→UUG | S→L | ndhBeU830SL |  |
|  | 2 | 1/1 | 836 | 279 | 2 | C→U | UCA→UUA | S→L | ndhBeU836SL |  |
|  | 3 | 1/1 | 872 | 291 | 2 | C→U | UCA→UUA | S→L | ndhBeU872SL |  |
|  | 4 | 1/1 | 1255 | 419 | 1 | C→U | CAU→UAU | H→Y | ndhBeU1255HY |  |
|  | 5 | 1/1 | 1481 | 494 | 2 | C→U | CCA→CUA | P→L | ndhBeU1481PL |  |
|  | 1 | 1/1 | 830 | 277 | 2 | C→U | UCG→UUG | S→L | ndhBeU830SL |  |
|  | 2 | 1/1 | 836 | 279 | 2 | C→U | UCA→UUA | S→L | ndhBeU836SL |  |
|  | 3 | 1/1 | 872 | 291 | 2 | C→U | UCA→UUA | S→L | ndhBeU872SL |  |
|  | 4 | 1/1 | 1255 | 419 | 1 | C→U | CAU→UAU | H→Y | ndhBeU1255HY |  |
|  | 5 | 1/1 | 1481 | 494 | 2 | C→U | CCA→CUA | P→L | ndhBeU1481PL |  |
| **ndhF** | 1 | 1/1 | 92 | 31 | 2 | U→C | AUG→ACG | M→T | ndhFeC92MT |  |
|  | 2 | 1/1 | 128 | 43 | 2 | C→U | CCC→CUC | P→L | ndhFeU128PL |  |
|  | 3 | 1/1 | 205 | 69 | 1 | U→C | UAU→CAU | Y→H | ndhFeC205YH |  |
|  | 4 | 1/1 | 271 | 91 | 1 | U→C | UCA→CCA | S→P | ndhFeC271SP |  |
|  | 5 | 1/1 | 290 | 97 | 2 | C→U | UCA→UUA | S→L | ndhFeU290SL |  |
|  | 6 | 1/1 | 316 | 106 | 1 | U→C | UUU→CUU | F→L | ndhFeC316FL |  |
|  | 7 | 1/1 | 586 | 196 | 1 | U→C | UUU→CUU | F→L | ndhFeC586FL |  |
|  | 8 | 1/1 | 667 | 223 | 1 | U→C | UUU→CUU | F→L | ndhFeC667FL |  |
|  | 9 | 1/1 | 692 | 231 | 2 | C→U | UCU→UUU | S→F | ndhFeU692SF |  |
|  | 10 | 1/1 | 704 | 235 | 2 | C→U | GCC→GUC | A→V | ndhFeU704AV |  |
|  | 11 | 1/1 | 1147 | 383 | 1 | U→C | UUU→CUU | F→L | ndhFeC1147FL |  |
|  | 12 | 1/1 | 1187 | 396 | 2 | U→C | AUU→ACU | I→T | ndhFeC1187IT |  |
|  | 13 | 1/1 | 1379 | 460 | 2 | U→C | AUU→ACU | I→T | ndhFeC1379IT |  |
|  | 14 | 1/1 | 1381 | 461 | 1 | C→U | CAU→UAU | H→Y | ndhFeU1381HY |  |
|  | 15 | 1/1 | 1570 | 524 | 1 | C→U | CUU→UUU | L→F | ndhFeU1570LF |  |
|  | 16 | 1/1 | 1841 | 614 | 2 | C→U | UCU→UUU | S→F | ndhFeU1841SF |  |
|  | 17 | 1/1 | 1856 | 619 | 2 | C→U | ACA→AUA | T→I | ndhFeU1856TI |  |
|  | 18 | 1/1 | 1933 | 645 | 1 | C→U | CCU→UCU | P→S | ndhFeU1933PS |  |
|  | 19 | 1/1 | 2018 | 673 | 2 | U→C | AUA→ACA | I→T | ndhFeC2018IT |  |
|  | 20 | 1/1 | 2027 | 676 | 2 | C→U | ACA→AUA | T→I | ndhFeU2027TI |  |
|  | 21 | 1/1 | 2135 | 712 | 2 | U→C | AUA→ACA | I→T | ndhFeC2135IT |  |
|  | 22 | 1/1 | 2165 | 722 | 2 | U→C | UUU→UCU | F→S | ndhFeC2165FS |  |
|  | 23 | 1/1 | 2186 | 729 | 2 | U→C | UUU→UCU | F→S | ndhFeC2186FS |  |
| **rpl3** | 1 | 1/1 | 29 | 10 | 2 | C→U | ACA→AUA | T→I | rpl32eU29TI |  |
| **ccsA** | 1 | 1/1 | 14 | 5 | 2 | C→U | ACU→AUU | T→I | ccsAeU14TI |  |
|  | 2 | 1/1 | 73 | 25 | 1 | C→U | CAU→UAU | H→Y | ccsAeU73HY |  |
|  | 3 | 1/1 | 88 | 30 | 1 | U→C | UUC→CUC | F→L | ccsAeC88FL |  |
|  | 4 | 1/1 | 326 | 109 | 2 | C→U | GCU→GUU | A→V | ccsAeU326AV |  |
|  | 5 | 1/1 | 335 | 112 | 2 | C→U | ACC→AUC | T→I | ccsAeU335TI |  |
|  | 6 | 1/1 | 392 | 131 | 2 | C→U | GCG→GUG | A→V | ccsAeU392AV |  |
|  | 7 | 1/1 | 512 | 171 | 2 | C→U | GCC→GUC | A→V | ccsAeU512AV |  |
|  | 8 | 1/1 | 521 | 174 | 2 | U→C | AUU→ACU | I→T | ccsAeC521IT |  |
|  | 9 | 1/1 | 566 | 189 | 2 | C→U | UCA→UUA | S→L | ccsAeU566SL |  |
|  | 10 | 1/1 | 629 | 210 | 2 | C→U | ACU→AUU | T→I | ccsAeU629TI |  |
|  | 11 | 1/2 | 637 | 213 | 1 | C→U | CUU→UCU | L→S | ccsAeUC637LS |  |
|  | 11 | 2/2 | 638 | 213 | 2 | U→C | CUU→UCU | L→S | ccsAeUC637LS |  |
|  | 12 | 1/1 | 641 | 214 | 2 | C→U | UCU→UUU | S→F | ccsAeU641SF |  |
|  | 13 | 1/1 | 671 | 224 | 2 | C→U | ACU→AUU | T→I | ccsAeU671TI |  |
|  | 14 | 1/1 | 848 | 283 | 2 | C→U | ACC→AUC | T→I | ccsAeU848TI |  |
|  | 15 | 1/2 | 895 | 299 | 1 | U→C | UCU→CUU | S→L | ccsAeCU895SL |  |
|  | 15 | 2/2 | 896 | 299 | 2 | C→U | UCU→CUU | S→L | ccsAeCU895SL |  |
| **ndhD** | 1 | 1/1 | 2 | 1 | 2 | C→U | ACG→AUG | T→M | ndhDeU2TM |  |
|  | 2 | 1/1 | 124 | 42 | 1 | U→C | UUC→CUC | F→L | ndhDeC124FL |  |
|  | 3 | 1/1 | 383 | 128 | 2 | C→U | UCA→UUA | S→L | ndhDeU383SL |  |
|  | 4 | 1/1 | 427 | 143 | 1 | U→C | UUC→CUC | F→L | ndhDeC427FL |  |
|  | 5 | 1/1 | 589 | 197 | 1 | U→C | UUU→CUU | F→L | ndhDeC589FL |  |
|  | 6 | 1/1 | 620 | 207 | 2 | C→U | GCC→GUC | A→V | ndhDeU620AV |  |
|  | 7 | 1/1 | 674 | 225 | 2 | C→U | UCG→UUG | S→L | ndhDeU674SL |  |
|  | 8 | 1/1 | 878 | 293 | 2 | C→U | UCA→UUA | S→L | ndhDeU878SL |  |
|  | 9 | 1/1 | 1240 | 414 | 1 | C→U | CCA→UCA | P→S | ndhDeU1240PS |  |
|  | 10 | 1/1 | 1256 | 419 | 2 | C→U | ACU→AUU | T→I | ndhDeU1256TI |  |
|  | 11 | 1/1 | 1298 | 433 | 2 | C→U | UCA→UUA | S→L | ndhDeU1298SL |  |
|  | 12 | 1/1 | 1310 | 437 | 2 | C→U | UCA→UUA | S→L | ndhDeU1310SL |  |
|  | 13 | 1/1 | 1355 | 452 | 2 | C→U | UCU→UUU | S→F | ndhDeU1355SF |  |
|  | 14 | 1/1 | 1405 | 469 | 1 | U→C | UUU→CUU | F→L | ndhDeC1405FL |  |
| **ndhE** | 1 | 1/2 | 43 | 15 | 1 | U→C | UCU→CUU | S→L | ndhEeCU43SL |  |
|  | 1 | 2/2 | 44 | 15 | 2 | C→U | UCU→CUU | S→L | ndhEeCU43SL |  |
|  | 2 | 1/1 | 233 | 78 | 2 | C→U | UCA→UUA | S→L | ndhEeU233SL |  |
| **ndhE** | 1 | 1/1 | 145 | 49 | 1 | U→C | UUC→CUC | F→L | ndhGeC145FL |  |
|  | 2 | 1/1 | 329 | 110 | 2 | C→U | UCG→UUG | S→L | ndhGeU329SL |  |
|  | 3 | 1/1 | 386 | 129 | 2 | C→U | UCA→UUA | S→L | ndhGeU386SL |  |
|  | 4 | 1/1 | 472 | 158 | 1 | U→C | UUC→CUC | F→L | ndhGeC472FL |  |
| **ndhI** | 1 | 1/1 | 4 | 2 | 1 | U→C | UUC→CUC | F→L | ndhIeC4FL |  |
|  | 2 | 1/1 | 253 | 85 | 1 | U→C | UUU→CUU | F→L | ndhIeC253FL |  |
|  | 3 | 1/1 | 379 | 127 | 1 | C→U | CUU→UUU | L→F | ndhIeU379LF |  |
| **ndhA** | 1 | 1/1 | 566 | 189 | 2 | C→U | UCA→UUA | S→L | ndhAeU566SL |  |
|  | 2 | 1/1 | 860 | 287 | 2 | U→C | UUC→UCC | F→S | ndhAeC860FS |  |
|  | 3 | 1/1 | 866 | 289 | 2 | C→U | CCU→CUU | P→L | ndhAeU866PL |  |
|  | 4 | 1/1 | 1072 | 358 | 1 | C→U | CUU→UUU | L→F | ndhAeU1072LF |  |
|  | 1 | 1/1 | 122 | 41 | 2 | C→U | ACA→AUA | T→I | ndhAeU122TI |  |
|  | 2 | 1/1 | 125 | 42 | 2 | U→C | AUA→ACA | I→T | ndhAeC125IT |  |
|  | 3 | 1/1 | 341 | 114 | 2 | C→U | UCA→UUA | S→L | ndhAeU341SL |  |
| **ndhH** | 1 | 1/1 | 10 | 4 | 1 | U→C | UCA→CCA | S→P | ndhHeC10SP |  |
|  | 2 | 1/1 | 14 | 5 | 2 | C→U | GCU→GUU | A→V | ndhHeU14AV |  |
|  | 3 | 1/1 | 964 | 322 | 1 | C→U | CCG→UCG | P→S | ndhHeU964PS |  |
|  | 4 | 1/1 | 983 | 328 | 2 | C→U | GCG→GUG | A→V | ndhHeU983AV |  |
| **rps15** | 1 | 1/1 | 137 | 46 | 2 | C→U | UCA→UUA | S→L | rps15eU137SL |  |
|  | 2 | 1/1 | 254 | 85 | 2 | C→U | UCA→UUA | S→L | rps15eU254SL |  |
| **ycf1** | 1 | 1/1 | 26 | 9 | 2 | C→U | CCA→CUA | P→L | ycf1eU26PL |  |
|  | 2 | 1/1 | 217 | 73 | 1 | U→C | UUC→CUC | F→L | ycf1eC217FL |  |
|  | 3 | 1/1 | 245 | 82 | 2 | U→C | GUG→GCG | V→A | ycf1eC245VA |  |
|  | 4 | 1/1 | 602 | 201 | 2 | U→C | UUU→UCU | F→S | ycf1eC602FS |  |
|  | 5 | 1/1 | 658 | 220 | 1 | U→C | UUU→CUU | F→L | ycf1eC658FL |  |
|  | 6 | 1/1 | 764 | 255 | 2 | U→C | AUU→ACU | I→T | ycf1eC764IT |  |
|  | 7 | 1/1 | 800 | 267 | 2 | C→U | GCA→GUA | A→V | ycf1eU800AV |  |
|  | 8 | 1/1 | 821 | 274 | 2 | C→U | UCU→UUU | S→F | ycf1eU821SF |  |
|  | 9 | 1/1 | 823 | 275 | 1 | U→C | UCU→CCU | S→P | ycf1eC823SP |  |
|  | 10 | 1/1 | 854 | 285 | 2 | C→U | CCG→CUG | P→L | ycf1eU854PL |  |
|  | 11 | 1/2 | 922 | 308 | 1 | C→U | CCC→UUC | P→F | ycf1eUU922PF |  |
|  | 11 | 2/2 | 923 | 308 | 2 | C→U | CCC→UUC | P→F | ycf1eUU922PF |  |
|  | 12 | 1/2 | 967 | 323 | 1 | U→C | UCU→CUU | S→L | ycf1eCU967SL |  |
|  | 12 | 2/2 | 968 | 323 | 2 | C→U | UCU→CUU | S→L | ycf1eCU967SL |  |
|  | 13 | 1/1 | 1027 | 343 | 1 | U→C | UAU→CAU | Y→H | ycf1eC1027YH |  |
|  | 14 | 1/1 | 1051 | 351 | 1 | C→U | CUU→UUU | L→F | ycf1eU1051LF |  |
|  | 15 | 1/1 | 1253 | 418 | 2 | U→C | UUU→UCU | F→S | ycf1eC1253FS |  |
|  | 16 | 1/1 | 1277 | 426 | 2 | C→U | UCU→UUU | S→F | ycf1eU1277SF |  |
|  | 17 | 1/1 | 1303 | 435 | 1 | U→C | UUU→CUU | F→L | ycf1eC1303FL |  |
|  | 18 | 1/1 | 1379 | 460 | 2 | U→C | UUU→UCU | F→S | ycf1eC1379FS |  |
|  | 19 | 1/2 | 1501 | 501 | 1 | U→C | UUU→CCU | F→P | ycf1eCC1501FP |  |
|  | 19 | 2/2 | 1502 | 501 | 2 | U→C | UUU→CCU | F→P | ycf1eCC1501FP |  |
|  | 20 | 1/1 | 1504 | 502 | 1 | U→C | UCA→CCA | S→P | ycf1eC1504SP |  |
|  | 21 | 1/1 | 1507 | 503 | 1 | C→U | CCU→UUU | P→F | ycf1eUU1507PF |  |
|  | 21 | 1/2 | 1508 | 503 | 2 | C→U | CCU→UUU | P→F | ycf1eUU1507PF |  |
|  | 22 | 1/1 | 1553 | 518 | 2 | U→C | UUU→UCU | F→S | ycf1eC1553FS |  |
|  | 23 | 1/1 | 1589 | 530 | 2 | C→U | ACU→AUU | T→I | ycf1eU1589TI |  |
|  | 24 | 1/1 | 1624 | 542 | 1 | C→U | CUC→UUC | L→F | ycf1eU1624LF |  |
|  | 25 | 1/1 | 1646 | 549 | 2 | C→U | ACA→AUA | T→I | ycf1eU1646TI |  |
|  | 26 | 1/1 | 1718 | 573 | 2 | U→C | AUU→ACU | I→T | ycf1eC1718IT |  |
|  | 27 | 1/1 | 1753 | 585 | 1 | C→U | CCC→UCC | P→S | ycf1eU1753PS |  |
|  | 28 | 1/1 | 1781 | 594 | 2 | C→U | UCU→UUU | S→F | ycf1eU1781SF |  |
|  | 29 | 1/1 | 1955 | 652 | 2 | C→U | ACU→AUU | T→I | ycf1eU1955TI |  |
|  | 30 | 1/1 | 1984 | 662 | 1 | C→U | CCU→UCU | P→S | ycf1eU1984PS |  |
|  | 31 | 1/1 | 2150 | 717 | 2 | C→U | CCC→CUC | P→L | ycf1eU2150PL |  |
|  | 32 | 1/1 | 2152 | 718 | 1 | C→U | CUC→UUC | L→F | ycf1eU2152LF |  |
|  | 33 | 1/1 | 2180 | 727 | 2 | C→U | CCG→CUG | P→L | ycf1eU2180PL |  |
|  | 34 | 1/1 | 2302 | 768 | 1 | C→U | CUU→UUU | L→F | ycf1eU2302LF |  |
|  | 35 | 1/1 | 2372 | 791 | 2 | C→U | UCA→UUA | S→L | ycf1eU2372SL |  |
|  | 36 | 1/1 | 2396 | 799 | 2 | C→U | GCC→GUC | A→V | ycf1eU2396AV |  |
|  | 37 | 1/1 | 2732 | 911 | 2 | U→C | UUA→UCA | L→S | ycf1eC2732LS |  |
|  | 38 | 1/1 | 2783 | 928 | 2 | C→U | GCU→GUU | A→V | ycf1eU2783AV |  |
|  | 39 | 1/1 | 2894 | 965 | 2 | C→U | CCA→CUA | P→L | ycf1eU2894PL |  |
|  | 40 | 1/1 | 2902 | 968 | 1 | U→C | UUU→CUU | F→L | ycf1eC2902FL |  |
|  | 41 | 1/2 | 2911 | 971 | 1 | U→C | UUG→CCG | L→P | ycf1eCC2911LP | X |
|  | 41 | 2/2 | 2912 | 971 | 2 | U→C | UUG→CCG | L→P | ycf1eCC2911LP |  |
|  | 42 | 1/1 | 2981 | 994 | 2 | C→U | UCA→UUA | S→L | ycf1eU2981SL |  |
|  | 43 | 1/1 | 3050 | 1017 | 2 | C→U | ACU→AUU | T→I | ycf1eU3050TI |  |
|  | 44 | 1/1 | 3179 | 1060 | 2 | U→C | AUA→ACA | I→T | ycf1eC3179IT |  |
|  | 45 | 1/1 | 3197 | 1066 | 2 | U→C | AUU→ACU | I→T | ycf1eC3197IT |  |
|  | 46 | 1/1 | 3215 | 1072 | 2 | U→C | UUC→UCC | F→S | ycf1eC3215FS |  |
|  | 47 | 1/1 | 3262 | 1088 | 1 | C→U | CUU→UUU | L→F | ycf1eU3262LF |  |
|  | 48 | 1/1 | 3290 | 1097 | 2 | C→U | ACC→AUC | T→I | ycf1eU3290TI |  |
|  | 49 | 1/1 | 3307 | 1103 | 1 | C→U | CUU→UUU | L→F | ycf1eU3307LF |  |
|  | 50 | 1/1 | 3385 | 1129 | 1 | C→U | CAU→UAU | H→Y | ycf1eU3385HY |  |
|  | 51 | 1/1 | 3434 | 1145 | 2 | U→C | UUU→UCU | F→S | ycf1eC3434FS |  |
|  | 52 | 1/1 | 3488 | 1163 | 2 | C→U | ACC→AUC | T→I | ycf1eU3488TI |  |
|  | 53 | 1/1 | 3511 | 1171 | 1 | U→C | UUC→CUC | F→L | ycf1eC3511FL |  |
|  | 54 | 1/1 | 3550 | 1184 | 1 | C→U | CUU→UUU | L→F | ycf1eU3550LF |  |
|  | 55 | 1/1 | 3623 | 1208 | 2 | U→C | AUC→ACC | I→T | ycf1eC3623IT |  |
|  | 56 | 1/2 | 3688 | 1230 | 1 | U→C | UUU→CCU | F→P | ycf1eCC3688FP |  |
|  | 56 | 2/2 | 3689 | 1230 | 2 | U→C | UUU→CCU | F→P | ycf1eCC3688FP |  |
|  | 57 | 1/1 | 3755 | 1252 | 2 | C→U | GCU→GUU | A→V | ycf1eU3755AV |  |
|  | 58 | 1/1 | 3889 | 1297 | 1 | U→C | UUU→CUU | F→L | ycf1eC3889FL |  |
|  | 59 | 1/1 | 3965 | 1322 | 2 | C→U | CCU→CUU | P→L | ycf1eU3965PL |  |
|  | 60 | 1/1 | 4057 | 1353 | 1 | C→U | CUU→UUU | L→F | ycf1eU4057LF |  |
|  | 61 | 1/1 | 4070 | 1357 | 2 | C→U | GCC→GUC | A→V | ycf1eU4070AV |  |
|  | 62 | 1/1 | 4081 | 1361 | 1 | U→C | UCC→CCC | S→P | ycf1eC4081SP |  |
|  | 63 | 1/2 | 4189 | 1397 | 1 | C→U | CCU→UUU | P→F | ycf1eUU4189PF |  |
|  | 63 | 2/2 | 4190 | 1397 | 2 | C→U | CCU→UUU | P→F | ycf1eUU4189PF |  |
|  | 64 | 1/1 | 4249 | 1417 | 1 | C→U | CUU→UUU | L→F | ycf1eU4249LF |  |
|  | 65 | 1/1 | 4387 | 1463 | 1 | C→U | CUU→UUU | L→F | ycf1eU4387LF |  |
|  | 66 | 1/1 | 4396 | 1466 | 1 | C→U | CCG→UCG | P→S | ycf1eU4396PS |  |
|  | 67 | 1/1 | 4450 | 1484 | 1 | C→U | CCC→UCC | P→S | ycf1eU4450PS |  |
|  | 68 | 1/2 | 4498 | 1500 | 1 | U→C | UUA→CCA | L→P | ycf1eCC4498LP | X |
|  | 68 | 2/2 | 4499 | 1500 | 2 | U→C | UUA→CCA | L→P | ycf1eCC4498LP |  |
|  | 69 | 1/1 | 4508 | 1503 | 2 | U→C | GUU→GCU | V→A | ycf1eC4508VA |  |
|  | 70 | 1/1 | 4514 | 1505 | 2 | C→U | UCA→UUA | S→L | ycf1eU4514SL |  |
|  | 71 | 1/1 | 4615 | 1539 | 1 | U→C | UUC→CUC | F→L | ycf1eC4615FL |  |
|  | 72 | 1/1 | 4634 | 1545 | 2 | U→C | UUG→UCG | L→S | ycf1eC4634LS |  |
|  | 73 | 1/1 | 4661 | 1554 | 2 | C→U | UCU→UUU | S→F | ycf1eU4661SF |  |
|  | 74 | 1/1 | 4736 | 1579 | 2 | U→C | GUU→GCU | V→A | ycf1eC4736VA |  |
|  | 75 | 1/1 | 4801 | 1601 | 1 | C→U | CUU→UUU | L→F | ycf1eU4801LF |  |
|  | 76 | 1/1 | 4873 | 1625 | 1 | U→C | UUU→CUU | F→L | ycf1eC4873FL |  |
|  | 77 | 1/1 | 4901 | 1634 | 2 | U→C | UUA→UCA | L→S | ycf1eC4901LS |  |
|  | 78 | 1/1 | 4993 | 1665 | 1 | C→U | CAU→UAU | H→Y | ycf1eU4993HY |  |
|  | 79 | 1/1 | 5105 | 1702 | 2 | C→U | UCA→UUA | S→L | ycf1eU5105SL |  |
|  | 80 | 1/1 | 5134 | 1712 | 1 | C→U | CCA→UCA | P→S | ycf1eU5134PS |  |
|  | 81 | 1/1 | 5234 | 1745 | 2 | U→C | UUC→UCC | F→S | ycf1eC5234FS |  |
