## Supplementary Fig. S1 for "Complete chloroplast genome of African Baobab (*Adansonia digitata* L.): structural characterization, comparative genomics, and phylogenetic placement within Malvaceae"

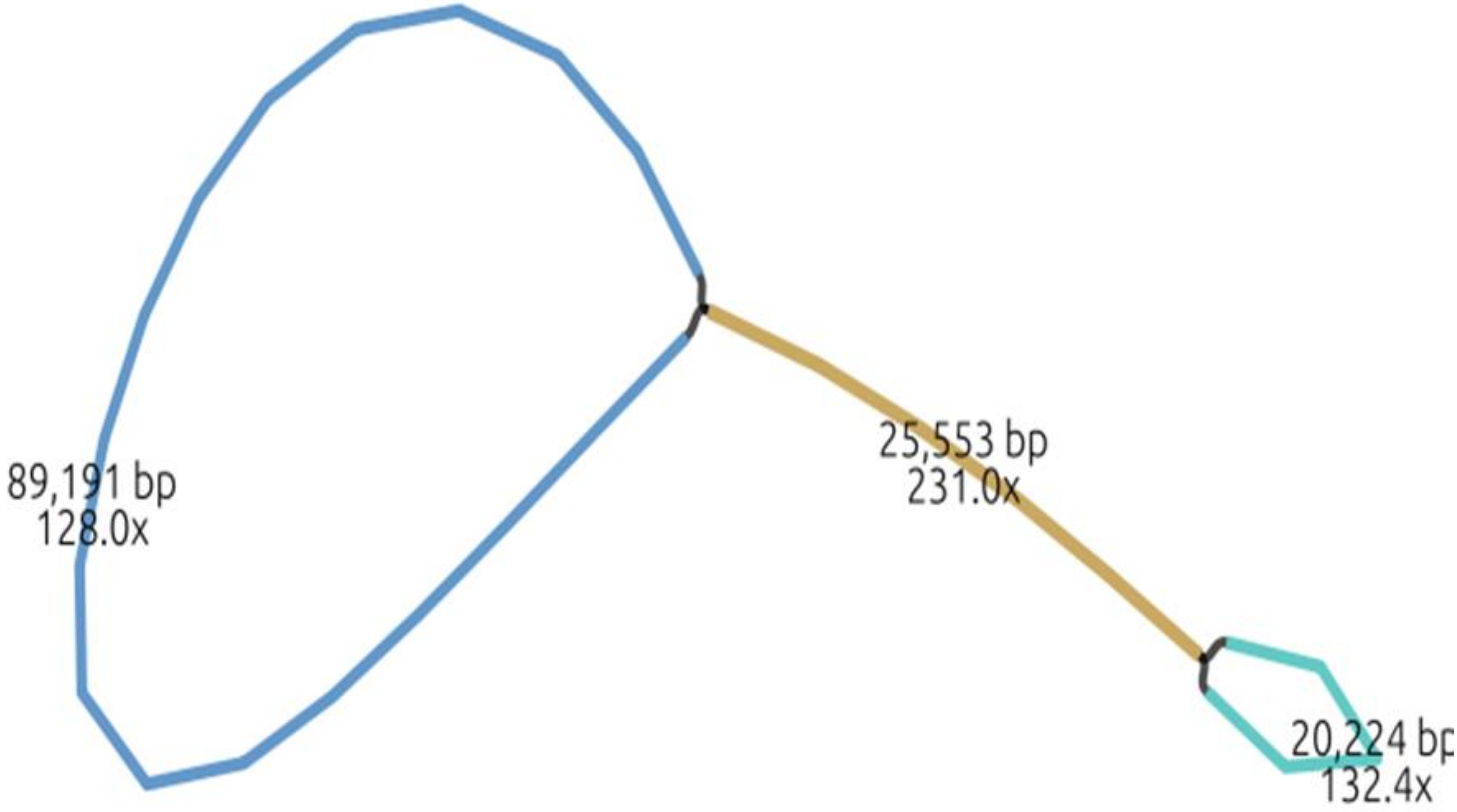


**Supplementary Figure 1.** Quadripartite architecture of the assembled *Adansonia digitata* chloroplast genome and read coverage across genomic regions. The graphical representation illustrates the organization of the chloroplast genome into its four structural regions: the large single-copy (LSC) region, the small single-copy (SSC) region, and two inverted repeat regions (IRa and IRb). Read coverage depth across each genomic region is displayed to indicate assembly depth.
