## Supplementary Fig. S2 for "Complete chloroplast genome of African Baobab (*Adansonia digitata* L.): structural characterization, comparative genomics, and phylogenetic placement within Malvaceae"

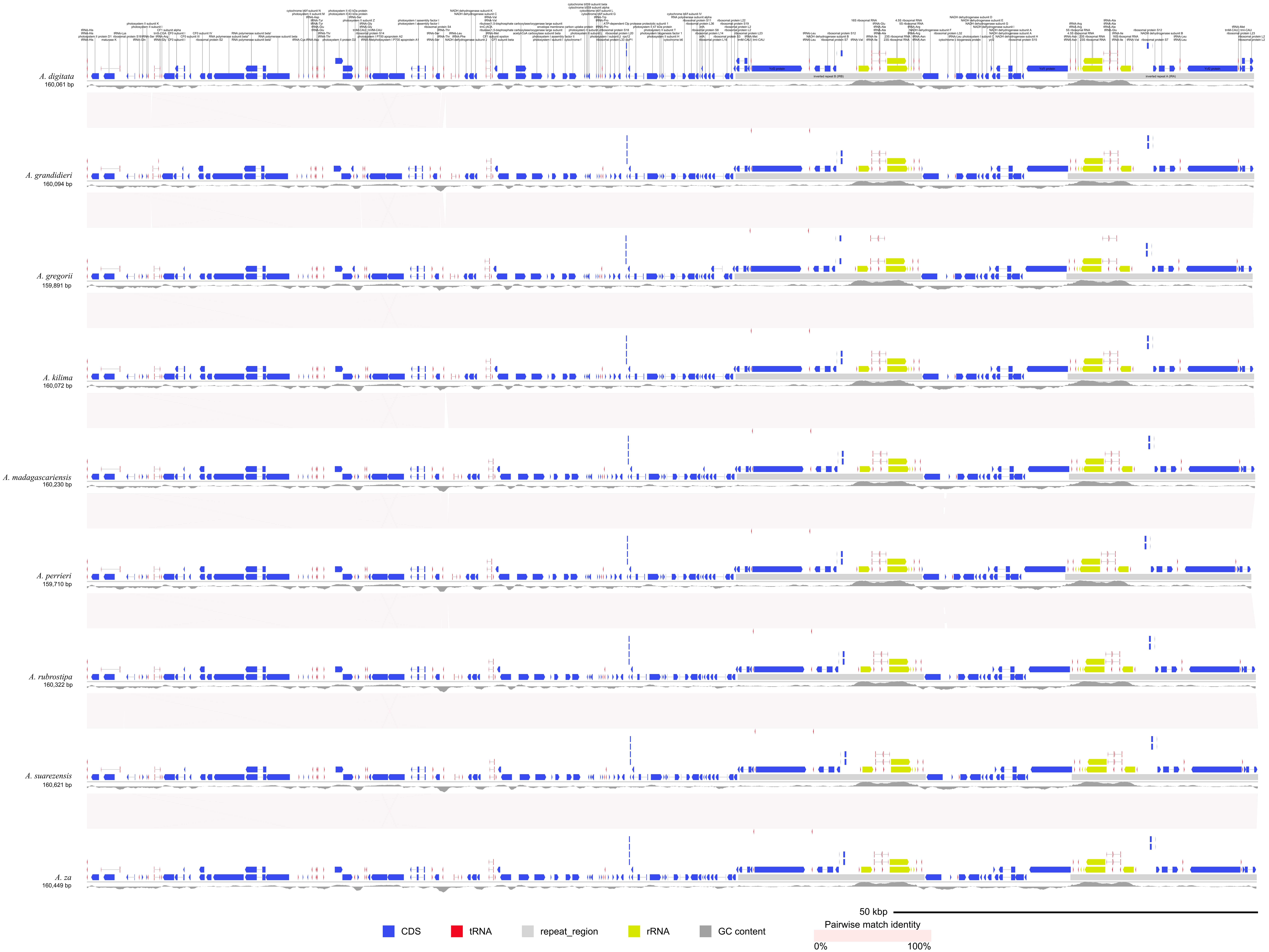


**Supplemntary Figure S2a**: Comparative schematic diagram illustrating the structural organization of the assembled *Adansonia digitata* chloroplast genome alongside known *Adansonia* species and the putative *A. kilima*. Gene overlaps across the large single-copy (LSC), small single-copy (SSC), and inverted repeat (IR) regions are displayed to highlight patterns of structural conservation and lineage-specific variation among the compared plastomes.

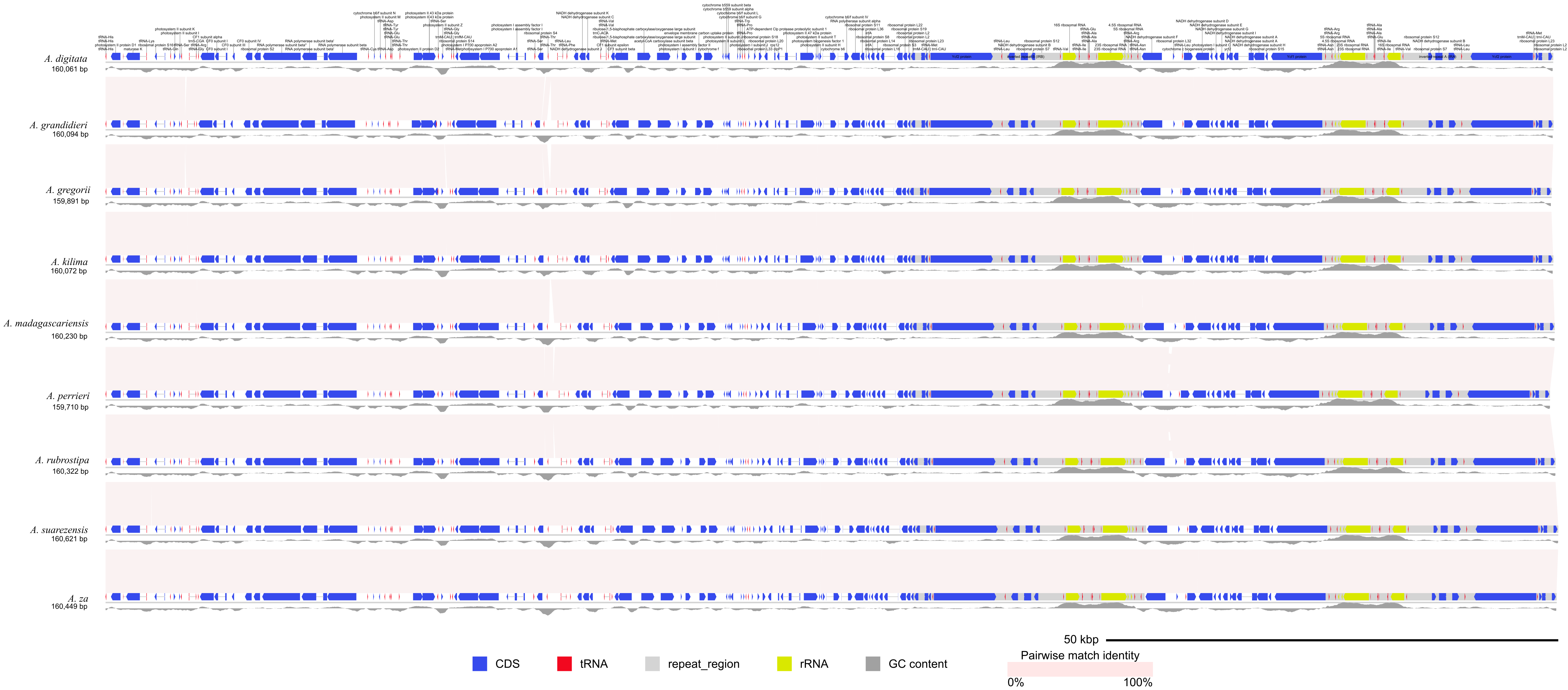


**Supplementary Figure 2b**: Comparative synteny analysis illustrating genomic collinearity between the assembled *Adansonia digitata* chloroplast genome, known *Adansonia* species, and the putative *A. kilima*. Syntenic blocks are shown to reveal patterns of structural conservation and lineage-specific rearrangements among the compared plastomes.
