## Supplementary Fig. S1 for "Complete chloroplast genome of African Baobab (*Adansonia digitata* L.): structural characterization, comparative genomics, and phylogenetic placement within Malvaceae"

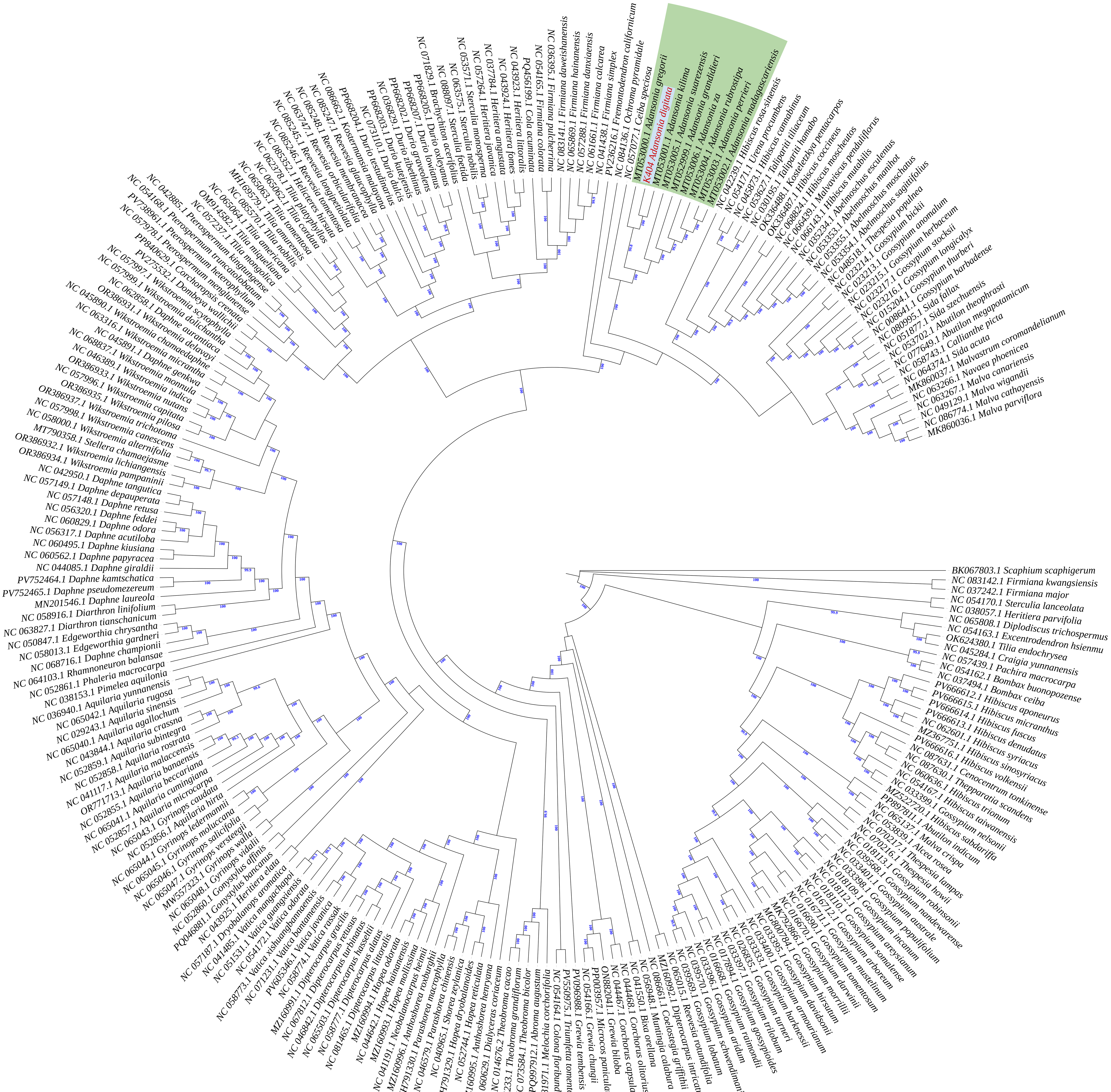

**Supplementary Figure 3a**. Maximum likelihood phylogenetic tree reconstructed from 248 Malvaceae chloroplast genomes, including the assembled *Adansonia digitata* plastome. The *Adansonia* clade is highlighted in green, and the assembled *A. digitata* genome is indicated in red. Bootstrap support values (1,000 replicates) are shown at each node. The tree was rooted on the branch leading to *Ceiba speciosa*.

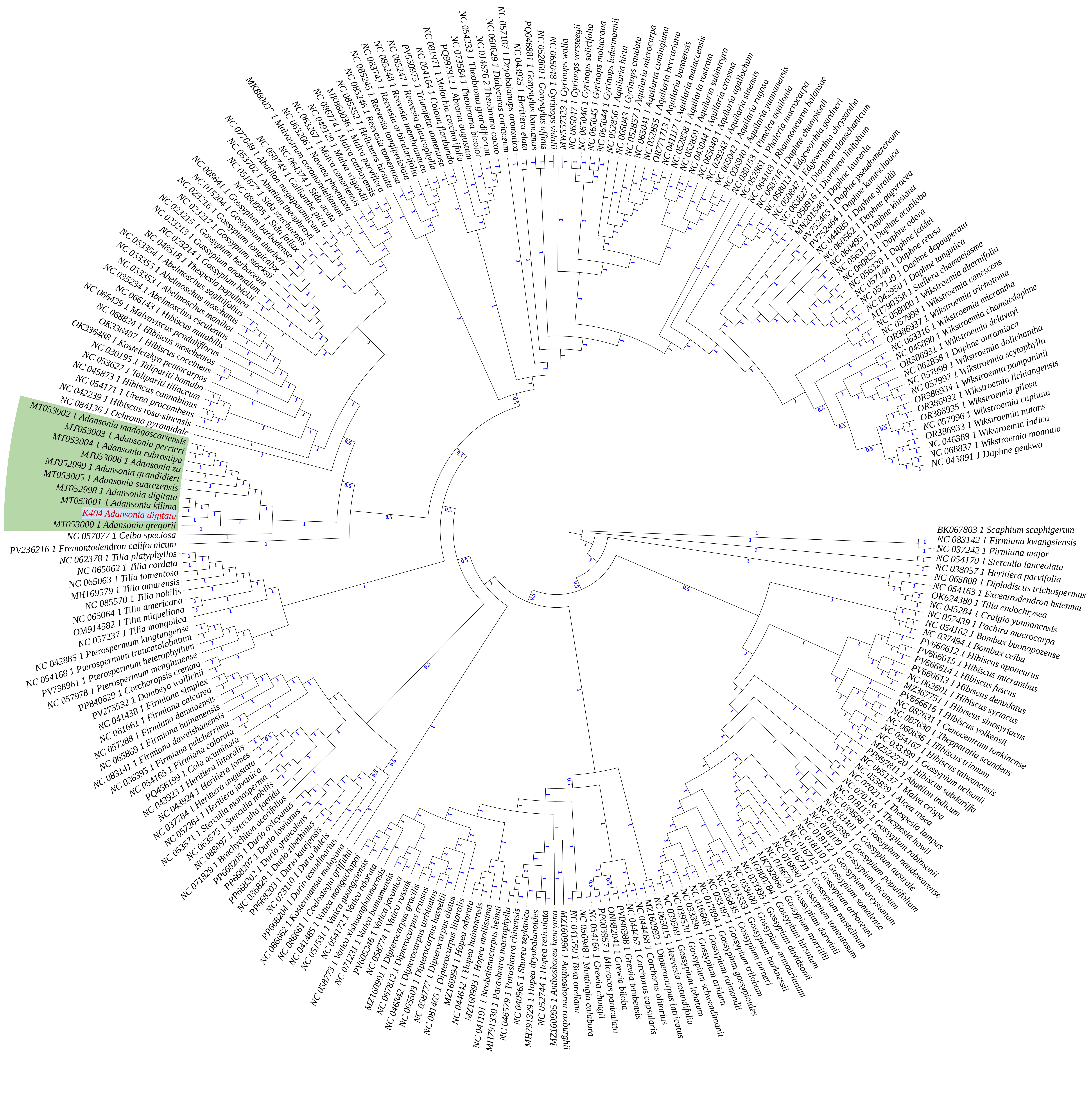

**Supplementary Figure 3b**. Bayesian inference phylogenetic tree reconstructed from 248 Malvaceae chloroplast genomes, including the assembled *Adansonia digitata* plastome. The *Adansonia* clade is highlighted in green, and the assembled *A. digitata* genome is indicated in red. Posterior probability support values are shown at each node. The tree was rooted on the branch leading to *Ceiba speciosa*.
